# Measurement of organ-specific and acute-phase blood protein levels in early Lyme disease

**DOI:** 10.1101/795344

**Authors:** Yong Zhou, Shizhen Qin, Mingjuan Sun, Li Tang, Xiaowei Yan, Taek-Kyun Kim, Juan Caballero, Gustavo Glusman, Mary E. Brunkow, Mark J. Soloski, Alison W. Rebman, Carol Scavarda, Denise Cooper, Gilbert S. Omenn, Robert L. Moritz, Gary P. Wormser, Nathan D. Price, John N. Aucott, Leroy Hood

## Abstract

Lyme disease results from infection of humans with the spirochete *Borrelia burgdorferi*. The first and most common clinical manifestation is the circular, inflamed skin lesion referred to as erythema migrans; later manifestations result from infections of other body sites. Laboratory diagnosis of Lyme disease can be challenging in patients with erythema migrans because of the time delay in the development of specific diagnostic antibodies against Borrelia. Reliable blood biomarkers for the early diagnosis of Lyme disease in patients with erythema migrans are needed. Here, we performed selected reaction monitoring, a targeted mass spectrometry-based approach, to measure selected proteins that 1) are known to be predominantly expressed in one organ (i.e., organ-specific blood proteins) and whose blood concentrations may change as a result of Lyme disease, or 2) are involved in acute immune responses. In a longitudinal cohort of 40 Lyme disease patients and 20 healthy controls, we identified 10 proteins with significantly altered serum levels in patients at the time of diagnosis, and we also developed a 10-protein panel identified through multivariate analysis. In an independent cohort of patients with erythema migrans, six of these proteins, APOA4, C9, CRP, CST6, PGLYRP2 and S100A9, were confirmed to show significantly altered serum levels in patients at time of presentation. Nine of the 10 proteins from the multivariate panel were also verified in the second cohort. These proteins, primarily innate immune response proteins or proteins specific to liver, skin or white blood cells, may serve as candidate blood biomarkers requiring further validation to aid in the laboratory diagnosis of early Lyme disease.

## INTRODUCTION

Lyme disease (LD), or Lyme borreliosis, is the most common tick-borne infectious disease in North America. According to estimates from the Centers for Disease Control and Prevention (CDC), LD accounted for 82% of all tick-borne disease reports during 2004–2016 ^1^. New cases of LD in the United States exceed 300,000 per year ^2-4^, and the geographic range of LD has been expanding recently in this country as well ^1^. The most common (70–90% of cases) and earliest clinical manifestation of LD is the skin lesion referred to as erythema migrans (EM), which typically occurs at the tick bite site at which the etiologic agent *Borrelia burgdorferi* was inoculated into the skin. *B. burgdorferi* may spread hematogenously from the initial site to other skin sites resulting in additional EM skin lesions, and/or resulting in the infection of other organs including the joints, nervous system and heart ^5, 6^. About 27–40% of LD patients with EM have abnormal liver function tests, but the cause of these abnormalities has not been determined ^7-9^.

Early diagnosis and treatment of patients with EM skin lesions is critical for preventing the development of extracutaneous manifestations of LD, such as Lyme arthritis and Lyme neuroborreliosis ^6, 10^. Due to a delay in the development of specific antibodies, laboratory diagnosis of LD in patients with or without EM often lacks optimal sensitivity during the first few weeks of infection—the time at which broad spectrum antibiotic treatment should be initiated ^11^. Direct diagnostic testing by culture or polymerase chain reaction (PCR) have several limitations including lower than ideal sensitivity, false negative test results (e.g., due to mutations in the sites of PCR amplification) and absence of effective assays that have been approved by the United States Food and Drug Administration (FDA).

There are currently over 40 diagnostic tests for LD in humans, which are available either commercially or from private laboratories ^11^. All of them fall into two main categories; indirect methods that detect the host immune responses against the pathogen, mainly the detection of IgM/IgG antibodies against *B. burgdorferi*, and direct methods to detect the pathogen itself.

The CDC-recommended two-step serological testing algorithm—which detects the presence of IgG and/or IgM antibodies against *B. burgdorferi* using first an enzyme immunoassay (typically an ELISA) and then Western blotting—is currently the most widely applied indirect approach to test for LD ^12, 13^. It performs well in later-stage disseminated LD but is not a sensitive test for early, localized LD, since IgM antibodies usually take 2 to 3 weeks to develop to detectable levels ^11, 14^. In addition, prompt antibiotic treatment of some patients with EM may prevent the development of diagnostic levels of anti-*B. burgdorferi* antibodies during the convalescent phase. Furthermore, both IgG and IgM antibodies specific for *B. burgdorferi* may persist long after symptoms have disappeared; therefore, their presence may not distinguish active current disease from historic earlier exposures ^11^. Since the 1990s, the specificity of first-tier serologic tests has improved slightly with the introduction of the C6 test ^15^ and the OspC7 ELISA ^16^. However, these newer tests still target IgG and/or IgM antibodies and do not improve the sensitivity or specificity to a satisfactory level for the early stage of disease ^11, 17^.

The main direct detection modalities in use are *B. burgdorferi* spirochete cultures and detection of bacterial DNA by PCR, but both of these methods can suffer from low sensitivity. Although the isolation and culturing of *B. burgdorferi* spirochetes is the gold standard to confirm diagnosis, culture is not a routine laboratory diagnostic test for LD due to its low success rate (40–60%), long incubation time (up to 8–12 weeks), susceptibility to contamination, and the level of expertise required ^18, 19^. Direct detection of outer surface membrane proteins of *B. burgdorferi*, e.g., OspA, by mass spectrometry has been reported in serum, but thus far only from three untreated LD patients ^20^.

Currently, no PCR-based assay has been cleared by the FDA for the diagnosis of LD. In patients with EM and a short duration of disease (<14 days), a *B. burgdorferi*-16S ribosomal RNA-based PCR on clinical specimens may be a useful diagnostic supplement ^21^. The blood-borne, i.e., bacteremic phase of *B. burgdorferi* is relatively brief and the concentration of spirochetes is generally quite low, therefore directly detecting the presence of *B. burgdorferi* DNA by PCR in blood from LD patients can be challenging ^22^. An assay employing a larger volume of blood coupled with a pre-enrichment step for *B. burgdorferi* DNA improved net sensitivity from 34% to only 62% ^23^. The bottom line is that none of these assays are suitable for the effective early detection of acute Lyme disease.

In this study, we employed a different strategy by targeting two classes of host proteins in blood whose blood concentration changes may reflect early Lyme infection: 1) acute-phase proteins (APP) involved in innate immune responses as the first line of defense against microorganisms; and 2) proteins synthesized specifically from organs known to be affected by *B. burgdorferi* infection (i.e., organ-specific proteins from liver and skin). APPs constitute part of the innate immune system that can respond within a few hours after bacteria enter the skin. Examples of APPs are C-reactive protein (CRP), the complement factors, ferritin, ceruloplasmin, serum amyloid A and haptoglobin. In response to injury or infection, local inflammatory cells secrete cytokines into the bloodstream, stimulating liver cells to increase (or decrease) the production of APPs that help the host to clear or inhibit the growth of pathogens.

Organ-specific transcripts are transcripts highly enriched in only one organ or expressed at least 10-fold over their expression levels in all other organs ^24^. Most proteins translated from organ-specific transcripts are involved in important organ-related functions and a few of these will be secreted into the blood through classic secretion, apoptosis, or shedding from the cell membrane by enzymatic cleavage. These blood organ-specific proteins may change their concentration levels in the blood when their cognate biological networks become disease perturbed—thus identifying the organ that is disease perturbed and pointing towards relevant pathology through the identification of their cognate biological networks. Thus, the levels of organ-specific blood proteins can be fingerprints of the health status of their corresponding organs. We have created a comprehensive list of organ-specific proteins that includes >2,500 proteins from 23 organs by mining more than 20 RNA and protein expression datasets for their expression patterns of these proteins (https://github.com/caballero/GeneAtlas). We have applied the highly sensitive and accurate targeted proteomics approach of selective reaction monitoring (LC-MS-SRM) ^25^ to measure the concentration of organ-specific proteins in blood and used this strategy successfully to identify candidate biomarkers for chronic and acute liver injury and other diseases ^26, 27^.

In this study, we measured the concentration of selected innate immune response and organ-specific proteins by LC-MS-SRM in LD samples in blood from the SLICE study (Study of Lyme Disease Immunology and Clinical Events) at Johns Hopkins University School of Medicine, and in an independent cohort from New York Medical College. We identified and verified serum proteins with altered abundances in patients during the early stage of *B. burgdorferi* infection, relative to healthy controls.

## MATERIALS AND METHODS

### ETHICS STATEMENT

This study was approved by the Institutional Review Boards of Johns Hopkins University and New York Medical College, as well as Western Institutional Review Board (WIRB), which approved the work performed at Institute for Systems Biology. It was conducted according to the principles of the Declaration of Helsinki. All patients provided written informed consent to participate.

### PATIENT CHARACTERISTICS

#### 1. DISCOVERY COHORT (SLICE)

The discovery cohort was obtained from Johns Hopkins University School of Medicine. These banked serum samples were previously collected under the project “Study of Lyme Disease Immunology and Clinical Events” (SLICE). This longitudinal sample set contained 40 participants with an EM skin lesion (greater than 5 cm in diameter); all patients included had either multiple EM skin lesions (n = 13) or a single lesion plus one or more systemic symptoms (n = 27). Patients were excluded if they had a history of LD, had received a LD vaccine, had a concomitant, objective extracutaneous manifestation of LD (such as neuro, cardiac, or arthritic complications), had an autoimmune condition, cancer, or a condition associated with non-specific fatigue and pain such as chronic fatigue syndrome, fibromyalgia, or major depression. All patients were untreated at the time of enrollment and were treated with antibiotic regimens consistent with current guidelines ^5^. The mean duration of illness prior to the first study visit was 9.5 ± 8.7 days.

The sample set included in the current study also comprised sera from 20 healthy controls. All controls were negative for prior LD by both self-report and two-tier serology performed at study entry; they were screened for the same exclusionary conditions as the cases with LD. The patients with LD and the healthy controls did not differ significantly in age or gender (**Table 1**). We analyzed serum samples collected from patients with LD at four time points: 1) pre-treatment (diagnosis/baseline), 2) 4 weeks post-diagnosis upon completion of antimicrobial treatment, 3) 6 months post-treatment, and 4) 12 months post-treatment, and from the control subjects we obtained samples at two time points (at the initial visit and 6 months later). Additional information on the study subjects can be found in **Table S1**. Serum was prepared according to standard protocol including clotting for 15-30 min at room temp before spinning at 3,000 RPM for 10 min, and storage at −80°C until analysis. Two-tier antibody testing, including a whole cell sonicate-based first-tier enzyme immunoassay, was performed by QUEST Diagnostics (Secaucus, NJ), a large commercial laboratory, following CDC interpretation guidelines at both the baseline and the 4 weeks post-treatment study visits.

**Table 1.** Demographic and clinical characteristics of patients with Lyme disease (n=40 in SLICE-JHU cohort, and n=30 in NYMC cohort) and healthy controls (n=20 for each of the cohorts) in the discovery and verification cohorts.

|  | n | Age, years | Female | Acute seropositive<sup>2</sup> at first visit | Ever seropositive<sup>2</sup> (acute/convalescent) | Multiple erythema migrans |
| --- | --- | --- | --- | --- | --- | --- |
| **SLICE-JHU** |  |  |  |  |  |  |
| Control | 20 | 54.9 ± 15.2 | 10 (50.0%) |  |  |  |
| Lyme | 40 | 48.9 ± 13.9 | 21 (52.5%) | 17 (42.5%) | 30 (76.9%) | 13 (32.5%) |
| *p*-value<sup>1</sup> |  | 0.132 | 0.855 |  |  |  |
| **NYMC** |  |  |  |  |  |  |
| Control | 20 | 49.1 ± 11.3 | 10 (50.0%) |  |  |  |
| Lyme | 30 | 49.1±10.9 | 18 (60.0%) | 16 (53.3%) | 20 (66.7%) | 10 (33.3%) |
| *p*-value<sup>1</sup> |  | 0.995 | 0.143 |  |  |  |

<sup>1</sup>Group comparisons for the SLICE-JHU, and separately for the NYMC cohorts, were performed using *chi*-squared test or Fisher's exact test for categorical variables, and *t*-test or Wilcoxon rank sum test of medians, where appropriate.
<sup>2</sup>Based on two-tier antibody testing conducted by large commercial laboratory following CDC interpretation of results for the SLICE-JHU cohort; based on a first-tier enzyme immunoassay performed at NYMC for the NYMC cohort.

#### 2. VERIFICATION COHORT (NYMC)

A similar sized LD patient cohort from the “Culture Confirmed Lyme Disease” and “Outcome of Lyme Disease” studies at New York Medical College (NYMC) was used to validate altered abundances of serum proteins in the early stage of LD. This verification cohort included serum samples from 30 patients with systemic LD symptoms in addition to the EM skin lesion at diagnosis (**Table 1**). Patients with concomitant, objective extracutaneous manifestations of LD were not included. Blood was drawn from all LD patients at 4 time points: 1) baseline (diagnosis), 2) convalescence (11.9 ± 5.9 days after antibiotic treatment started), 3) one-year post-treatment, and 4) 4-6 years post-treatment; blood was obtained from 20 healthy controls at 2 time points: baseline and one year later. No information on the duration of illness prior to the baseline visit was available. Blood was drawn into serum separator tubes (SST) and allowed to clot for 15-30 min at room temp. Tubes were spun at 3,000 RPM for 10 min, and serum poured off into tubes before being stored at −80°C until analysis. A whole cell sonicate-based first-tier enzyme immunoassay (EIA) was performed at NYMC for all of the patients at both the baseline and convalescence time points ^28^. A C6 test was performed for the controls (Immunetics, Inc., now Oxford Immunotec USA, Marlborough, MA).

All patients from NYMC were untreated at the time of enrollment and were then treated with antibiotic regimens consistent with current guidelines ^5^. Selected subject demographic and clinical characteristics can be found in **Table S1**. The sample distributions in the two LD cohorts are summarized in **Table S2**.

### SERUM SAMPLE PREPARATION FOR SRM ANALYSIS

Each serum sample was diluted 1:4 (v/v) in phosphate-buffered saline and spun through a .22 μm filter (Costar® Spin-X® centrifuge tube, Corning) at 10,000 x g to remove debris before being transferred to a clean polypropylene tube and stored at −80°C. To reduce the complexity of the serum proteome, an immuno-affinity depletion LC column (Multiple Affinity Removal Column Human 14 column, 4.6 × 100 mm, Agilent, Santa Clara, CA) was used to remove the 14 most abundant serum proteins. This procedure generally results in a 20-fold enrichment of low-abundance proteins. Samples were processed in random order. A reference pooled serum sample was added after every 50 sample runs.

Aliquots of 100 µL of 1:4 (v/v) diluted and filtered serum samples were loaded on the depletion column. Manufacturer recommended LC program was applied. The flow-through fractions (1.2 mL) were collected and concentrated to 200 µL using Amicon Ultra Centrifugal Filters (4 mL 3K MWCO, MilliporeSigma, Burlington, MA). BCA protein assay was performed to measure protein concentrations before and after depletion. Proteins were denatured in 50% (v/v) TFE (2,2,2-Trifluoroethanol) at 37°C for 30 min before reduction (TCEP), alkylation (iodoacetamide) and digestion with trypsin (1:25 trypsin/protein ratio) for 14 hours. After desalting using 1 cc C18 and MCX cartridges (Waters, Milford, MA), the proper amounts of stable C^13^N^15^ isotope-labeled (SIL) synthetic standards for target proteins were spiked into experimental samples before analysis in Agilent 6490 triple-quadrupole mass spectrometers (Agilent, Santa Clara, CA). A yeast protein (enolase 1, P00924) was spiked in as internal standard before depletion. All chemical reagents were obtained from Sigma-Aldrich (St. Louis, MO).

### TARGET PROTEIN LIST FOR SRM ANALYSIS

We have developed Gene Atlas Interface, a discovery system that facilitates the identification of transcripts/genes that are highly expressed in one particular organ or tissue. Currently, this system stores 21 publicly available microarray and RNA-seq gene expression datasets for more than 30 human and mouse tissues (body atlas). Microarray data were processed from the raw data (CEL files) with the R:Bioconductor “affy” and “limma” packages. All samples were normalized using the Robust Multi-Array Average (RMA) method and gene expression values were stored in individual tables. RNA-seq data were processed from raw reads (FASTQ files), after removing low-quality reads, repeats and ribosomal sequences. The reads were mapped to the human reference genome GRCh37 with an optimized version of Blat, then converted to BAM and transcript quantification was achieved with Cufflinks using the gene models of Ensembl r64 as described in the following references ^26, 29^.

Our pipeline allows comparison of expression levels of all transcripts in one or more tissues/organs of interest with all other organs in the same dataset, providing two methods to evaluate significance: 1) normalized value of the log-enriched-ratio multiplied by the normalized expression level, and 2) the lower bound of the Wilson score confidence interval for a Bernoulli parameter (95% confidence). Both methods can use the maximal value observed or an average of values. Candidate transcripts from a specific organ or tissue are compared with all other organs/tissues in the same dataset, retaining only those showing > 10-fold enrichment over all other organs summed and *P* value < 0.005. The transcript lists thus generated from each dataset are then compared. Because each transcriptional dataset may discover a different group of “organ-specific transcripts” that overlap to varying extents, we determined that, in general, a gene is most likely organ-specific if four or more out of the 21 independent datasets support the call. Genes (transcripts) identified by three or fewer datasets were evaluated by their log-enriched ratio, agreement of mRNA expression data with protein expression data as available, and manual inspection (for example, determining if it is a known protein with organ-specific function or alternatively a product of a housekeeping gene for basic cell functions).

The GeneAtlas pipeline, including software and datasets, is publicly available at https://github.com/caballero/GeneAtlas.

### SRM ASSAYS

Serum samples were analyzed in a triple-quadrupole mass spectrometer (Agilent 6490, Agilent, Santa Clara, CA) integrated with a nanospray ion source and Chip Cube nano-HPLC. Three to four pairs of light (endogenous) and heavy (stable C^13^N^15^ isotope-labeled (SIL) synthetic standards) transitions were monitored for each target peptide. A 90-min gradient of acetonitrile from 3% to 40% was used to elute peptides from a high-capacity nano-HPLC Chip (160 nL, 150 mm X 75 μm ID, Agilent, Santa Clara, CA) as described elsewhere ^26^. Other settings included operating the nano-HPLC separation chip at 0.3 μL/min nano pump flow rate, 2 μg peptide loading amount, 1820 V capillary voltage and Dynamic MRM with 200 Delta EMV (+). Duplicate runs were performed for each sample. All SRM parameters and results are deposited in the SRM chromatographic repository at ISB and are publicly available (http://www.srmatlas.org and http://www.peptideatlas.org/passel/, PASS01453). The final SRM methods used for 174 proteotypic peptides are summarized in **Supplement Table 3** in PeptideAtlas SRM Experiment Library (PASSEL) format.

### DATA ANALYSIS

Raw SRM mass spectrometry data were processed using the Skyline targeted proteomics environment ^30^. For each peptide, the total peak area and light/heavy (L/H) ratio of all monitored transitions were calculated and manually checked before export. The quantifier transition(s) were selected and used in further calculations in cases of high background and overlapping peaks from co-eluted peptides. Although the same amount of total peptide (2 μg) from each depleted sample was loaded into the mass spectrometer, the L/H ratios were adjusted by serum volume such that the relative peptide abundances from the same volume of serum were compared. The peptide L/H ratios were then log_2_-transformed for a more normal-like distribution. The peptide levels were normalized against the mean of the 20 control samples from two time points since there was no significant difference in controls between the two time points. Significantly differentially expressed proteins were identified by both Student’s *t*-test and multivariate analysis (MVA).

*T*-tests (Benjamini-Hochberg adjusted *p*-values) were used to identify differentially expressed proteins ^31^. The area under the curve (AUC) for the receiver operating characteristics (ROC) curve were generated in Prism 6.0 (GraphPad Software). Unsupervised principal component analysis (PCA) was performed in Perseus (1.6) ^32^ and hierarchical cluster analysis (HCA) heatmaps were generated in MultiExperiment Viewer (MeV, 4.9.0) ^33^.

#### Partial AUC (pAUC)

Partial AUC was applied to summarize a portion of the AUC curve over the pre-specified range of interest (leading to a low False Discovery Rate) such that only a portion of cases would be identified but with high confidence (>90% specificity) ^34^.

#### Combined AUC

Logistic regression modeling was applied to establish a prediction model with multiple individual peptides identified by Student’s *t*-test. The AUCs for both whole-set and in 10-fold cross-validations (CV) were then calculated to assess the effectiveness of this model.

#### Multivariate analysis (MVA)

In addition to the *t*-test, a multivariate algorithm similar to that used for an earlier study in a proteomic characterization of pulmonary nodules ^35^ was developed to identify the most informative proteins. In the discovery SLICE set, Monte Carlo Cross-Validation (MCCV) was performed on 10^6^ panels with 5-fold cross-validation. Each panel was composed of 10 features randomly selected from all measured peptides and fitted to a logistic regression model, using a 20% holdout rate and 100 sample permutations. In the verification NYMC set, the same model from the discovery SLICE set was applied to calculate the ROC curve.

### WESTERN BLOT ANALYSIS OF PATIENT SERUM

Aliquots of 0.5 μL of serum (not depleted) from each sample were loaded on 4-12% SDS-PAGE Bis-Tris gels (Life Technologies, Grand Island, NY USA). Following separation, the proteins were transferred to PVDF membranes using an iBlot® dry blotting system from Life Technologies (Thermo Fisher Scientific, Carlsbad, CA). The membrane was blocked for non-specific binding sites in 5% non-fat milk, 0.1% (v/v) Tween 20 in PBS at room temperature for 1 hour, then incubated with primary antibody overnight at 4°C. The membrane was washed three times in 0.1% Tween in PBS for 5 minutes each time and then incubated with horseradish peroxidase (HRP)-conjugated secondary antibodies for 1 hour at room temperature. Detection was carried out by SuperSignal West Dura enhanced Duration substrate (Thermo Scientific, cat # 34076, Rockford, IL) with FluorchemE (ProteinSimple, Santa Clara, CA). The images were analyzed using ImageJ ^36^. Commercial rabbit polyclonal antibodies against ALDOB (Cat # 18065-1-AP, 1:1000), CRP (Cat # 24175-1-AP, 1:500), CST6 (Cat # 17076-1-AP, 1:200), and FBP1 (Cat # 12842-1-AP, 1:500) were obtained from ProteinTech (Rosemont, IL).

## RESULTS AND DISCUSSION

The objective of this study was to identify blood proteins with altered abundances in patients with early stage Lyme disease as compared to healthy controls. Such proteins are candidate biomarkers requiring further validation that may ultimately lead to the development of improved diagnostic tests for early Lyme disease, especially in patients with a negative antibody-based serologic test. We used the targeted LC-MS-SRM proteomics technique to monitor selected proteins that fall within two categories: 1) key innate immunity regulators including acute phase proteins, reflecting the fact that LD is an acute inflammatory disease; and 2) organ-specific blood proteins predominantly expressed in organs and tissues known to be affected by *B. burgdorferi*, i.e., liver, heart, brain and skin ^5, 37^. We hypothesize that at least a subset of these organ-specific proteins may serve as surrogate indicators of the health status of their cognate organs (i.e., when networks are disease-perturbed, blood levels of their individual component proteins frequently change).

We designed SRM assays for 456 proteins with 2–3 proteotypic peptides per protein, using the SRMAtlas, a database containing SRM assays for more than 99.7% of the human proteome ^38^. Two pooled sera, one from the discovery LD sample set of 40 patients at the baseline time point and the other from the healthy controls, were used to pre-screen the detectability of all selected peptides in LC-MS-SRM. In all, 174 peptides (representing 124 human proteins) were detected and selected to be measured in individual serum samples. In the discovery sample set of 40 LD patients (each with 4 time points) and 20 healthy controls (each with 2 time points) from baseline to 12 months post-treatment, 90 proteotypic peptides from 73 targeted proteins were successfully detected and quantified.

### 1. Protein Changes Associated with Early Stage Lyme Disease

#### 1a. Proteins Identified by Student’s *t*-test

In the discovery cohort, we first looked for proteins with significantly different serum levels in LD patients at the baseline visit (i.e., time of diagnosis) relative to healthy controls by *t*-test (two-tailed distribution, homoscedastic) with a Benjamini–Hochberg adjusted P value cutoff < 0.05 to adjust for multiple hypothesis comparisons, corresponding to roughly an uncorrected *t*-test *P* < 0.005. At this level of significance, we identified 10 proteins (represented by 13 proteotypic peptides) with serum levels significantly elevated (ALDOB, C9, CRP, FBP1 and S100A9) or reduced (AFM, APOA4, CST6, ITIH2 and PGLYRP2) in LD patients (**Figure 1A** and **Table 2**). We refer to these proteins as the ***t*-Test Set**. Before treatment, in response to inflammatory stimuli by *B. burgdorferi* infection, CRP serum levels increased the most, from 3 to 276-fold in 30 of the 40 LD patients (75%), a finding consistent with other studies ^39, 40^.

**Table 2.**
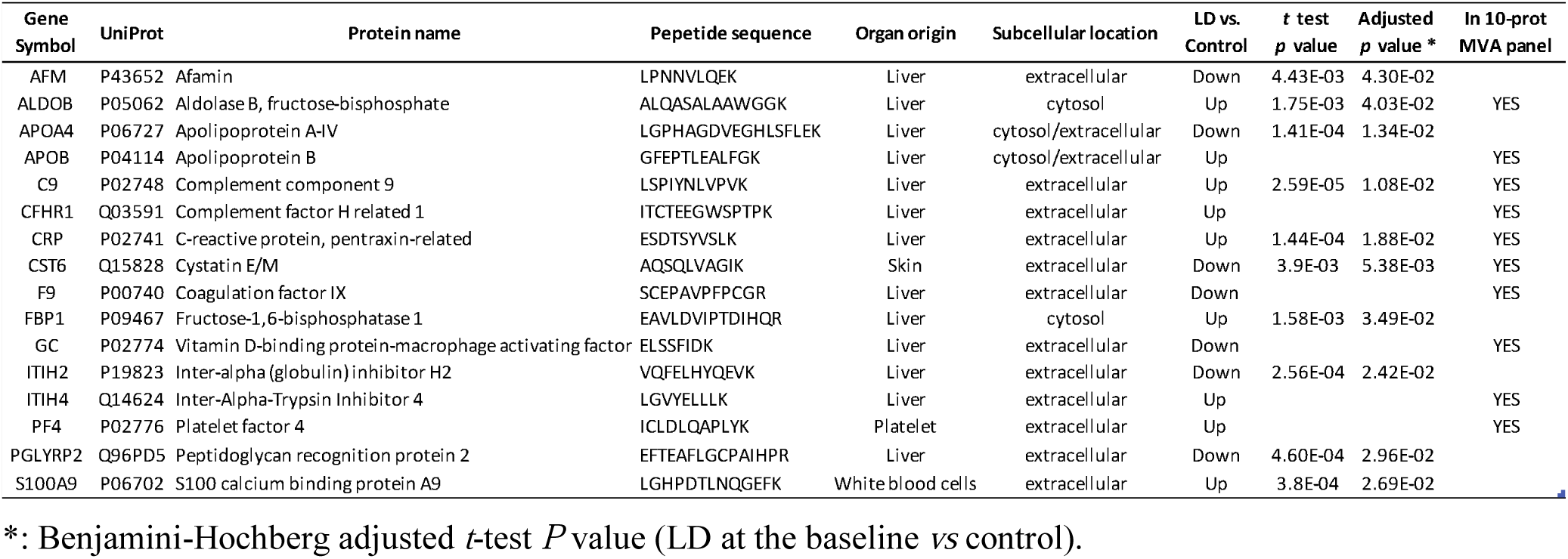
Summarized information of the 16 LD-associated blood proteins discovered in the discovery SLICE cohort.

**Figure 1.**
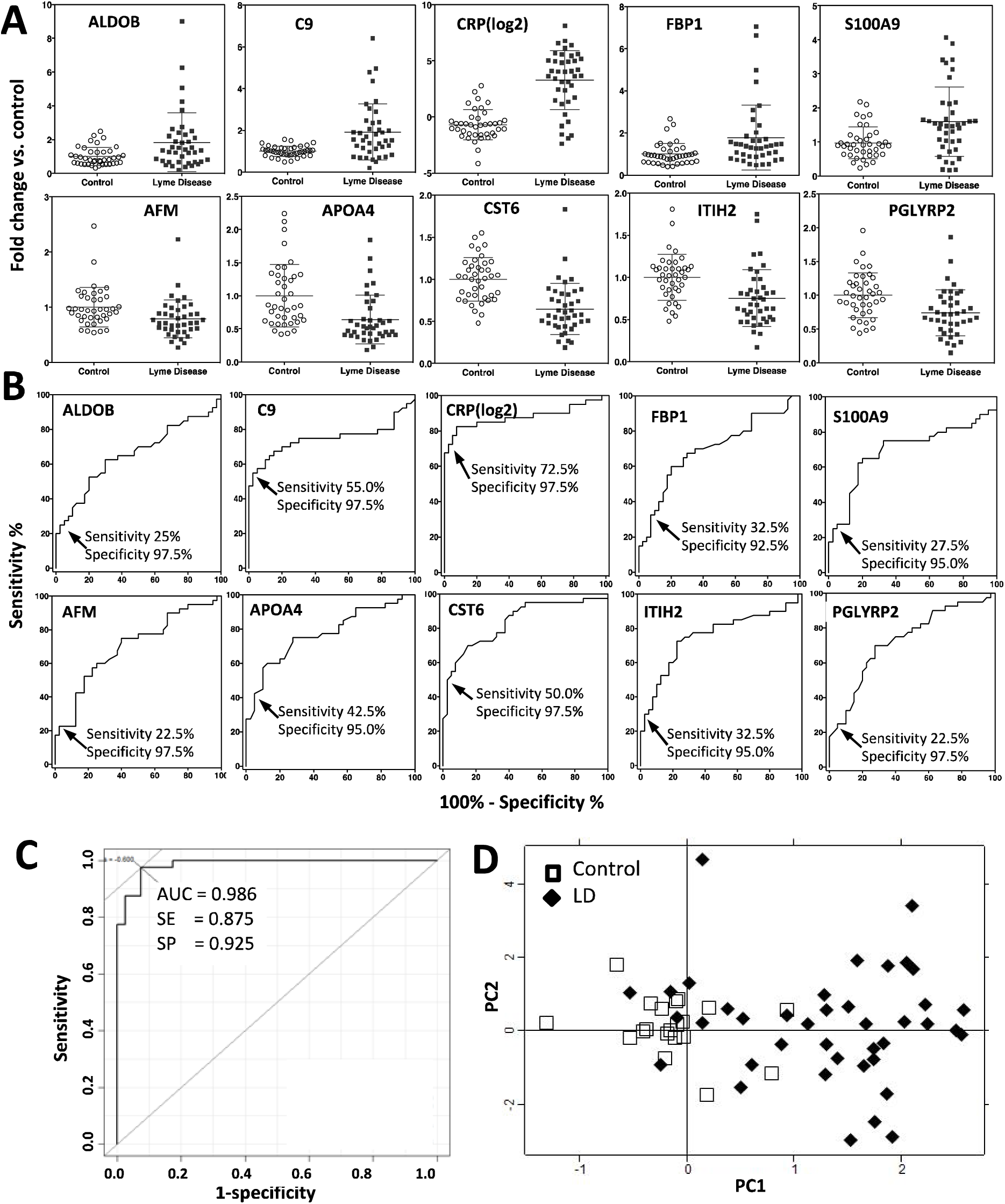
The 10 early Lyme disease-associated proteins identified by *t*-test in the discovery cohort (***t*-Test Set**). **1A**, Dot plots showing the distribution and mean of relative serum levels with standard deviation for 10 proteins with altered serum levels associated with LD by *t*-test (Benjamini–Hochberg P < 0.05), at the baseline time point (i.e., at LD diagnosis). The average of control = 1. **1B**, Individual AUCs. Cutoff points are shown where low false positive rates (2.5% − 7.5%) are achieved with variable sensitivities. **1C**, Combined AUC (whole set) of the 10 proteins to distinguish LD patients from healthy controls. **1D**, Biplots of PC1 and PC2 scores of 40 LD patients and 20 healthy controls from unsupervised PCA analyses, computed using the 10 ***t*-Test Set** proteins (13 proteotypic peptides). Filled diamonds and open squares represent PC scores for LD patients and controls, respectively.

We computed the area under the curve (AUC) for the receiver operating characteristics (ROC) curve for each of the 10 proteins (**Figure 1B**). Then we computed partial AUCs to identify the cut-off points of each protein in serum to reach a high specificity, i.e., between a 2.5% and 7.5% false positive rate in the ROC curve, with variable sensitivities (22.5% to 72.5%) in all 40 patients with LD (**Figure 1B**).

At these cutoff levels with high specificity but relatively low sensitivity, only a variable subset of LD patients could be classified by each of the 10 proteins. However, when all 10 ***t*-Test Set** proteins were combined, all 40 patients with LD had at least one protein exceeding the cutoff threshold (right column in **Table 3**). When considering patients as LD positive by at least two of the 10 proteins, 36 out of the 40 patients still could be identified at baseline, i.e., a sensitivity of 90%. The combined AUC of 10 proteins, when combined into a single prediction model using logistic regression, was 0.938 in 10-fold CV (0.986 whole set), with 0.85 in sensitivity and 0.93 in specificity (**Figure 1C**). To visualize the performance of these 10 proteins (represented by 13 proteotypic peptides) in identifying patients with LD, we also performed unsupervised principal component analysis (PCA). The distinction between LD patients and healthy controls was apparent at baseline (**Figure 1D**).

**Table 3.** LD patients identified at baseline by the 10 proteins as described in **Figures 1** and **2**. Each of the 10 proteins identifies patients with various sensitivity at the indicated cutoff level of fold change compared to average of healthy controls as shown in **Figure 1B**, with low false positive rates (2.5-- 7.5%). A combination of 10 proteins identified all 40 LD patients at the indicated cutoff protein levels. In the righthand column, the number of proteins with serum level exceeding the cutoff fold change is listed for every LD patient.

| Patient ID | *AFM*<br>< 0.55* | *ALDOB*<br>> 2.2 | *APOA4*<br>< 0.46 | *C9*<br>> 1.45 | *CRP*<br>> 2.7 | *CST6*<br>< 0.58 | *FBP1*<br>> 1.6 | *ITIH2*<br>< 0.60 | *PGLYRP2*<br>< 0.58 | *S100A9*<br>>1.8 | # of "+"<br>proteins |
| --- | --- | --- | --- | --- | --- | --- | --- | --- | --- | --- | --- |
| LD01 |  |  |  | + |  |  |  |  |  | + | 2 |
| LD02 |  |  |  | + |  |  | + |  |  |  | 2 |
| LD03 |  | + | + | + | + |  | + | + |  | + | 7 |
| LD04 |  | + | + |  |  | + | + | + |  |  | 5 |
| LD05 |  |  |  | + | + |  |  |  |  | + | 3 |
| LD06 |  | + | + | + |  | + | + |  |  |  | 5 |
| LD07 |  |  |  | + | + | + |  | + |  |  | 4 |
| LD08 |  |  |  |  | + | + |  |  |  | + | 3 |
| LD09 |  |  |  | + | + | + |  |  |  |  | 3 |
| LD10 |  |  | + | + | + |  |  |  |  |  | 3 |
| LD11 |  |  | + |  | + |  |  |  |  |  | 2 |
| LD12 |  |  |  |  | + | + |  |  | + |  | 3 |
| LD13 |  |  |  | + | + | + |  |  |  |  | 3 |
| LD14 |  |  |  | + | + |  |  |  |  |  | 2 |
| LD15 | + |  |  | + |  |  |  |  |  | + | 3 |
| LD16 |  |  | + |  |  |  |  |  |  | + | 2 |
| LD17 | + |  |  |  |  |  |  | + | + |  | 3 |
| LD18 |  |  | + |  | + |  |  |  |  |  | 2 |
| LD19 |  |  |  |  | + |  |  |  |  |  | 1 |
| LD20 |  |  |  |  | + | + |  |  |  |  | 2 |
| LD21 |  |  | + |  |  | + |  | + | + |  | 4 |
| LD22 |  |  |  |  | + | + |  |  |  |  | 2 |
| LD23 |  |  |  | + | + |  | + |  |  |  | 3 |
| LD24 | + |  |  |  | + |  |  |  |  |  | 2 |
| LD25 | + | + |  | + |  | + | + |  |  | + | 6 |
| LD26 |  |  |  |  | + |  |  |  |  |  | 1 |
| LD27 |  |  |  |  | + | + |  | + | + | + | 5 |
| LD28 | + |  | + |  | + | + |  | + | + |  | 6 |
| LD29 |  |  | + | + | + |  |  | + | + |  | 5 |
| LD30 |  |  | + | + | + | + |  | + |  |  | 5 |
| LD31 |  |  | + | + | + | + | + |  |  |  | 5 |
| LD32 |  | + |  | + | + |  | + |  |  | + | 5 |
| LD33 |  | + | + | + | + |  | + |  |  | + | 6 |
| LD34 |  | + |  | + | + |  | + |  |  | + | 5 |
| LD35 |  | + | + | + |  | + | + |  |  |  | 5 |
| LD36 |  |  | + |  | + | + |  |  |  |  | 3 |
| LD37 | + |  |  | + |  | + |  | + | + |  | 5 |
| LD38 | + | + | + | + |  | + | + |  | + |  | 7 |
| LD39 | + | + | + | + | + | + | + | + | + |  | 9 |
| LD40 | + |  | + |  |  | + |  | + | + |  | 5 |
| # LD patient | 9 | 11 | 18 | 23 | 31 | 21 | 13 | 15 | 13 | 11 |  |
| % LD patient | 22.5% | 27.5% | 45.0% | 57.5% | 77.5% | 52.5% | 32.5% | 37.5% | 32.5% | 27.5% |  |

\*: the cut-off fold change of serum concentration compared to average of healthy controls.
+: patient identified by each of the 10 proteins with relative serum concentrations beyond the cut-off protein values.

#### 1b. Proteins Identified by Multivariate Analysis (MVA)

An alternative way to discover disease-associated molecules with changing concentration levels is to identify features that have synergistic effects ^35, 41^. In this approach we started with the 90 peptides quantified by LC-MS-SRM and randomly generated one million panels of 10 features. We assessed each of these panels for their ability to distinguish LD from healthy controls. We then defined a protein as “cooperative” if it was found more frequently in best performing panels than expected by chance alone. To identify the cooperative proteins, we implemented a multivariate algorithm similar to that used in a proteomic blood diagnosis to distinguish benign from neoplastic pulmonary nodules ^35^. The MVA analysis revealed 10 proteins (utilizing 11 proteotypic peptides) that were cooperative in distinguishing LD patients at baseline from healthy controls: ALDOB, APOB, C9, CFHR, CRP, CST6, GC, F9, ITIH4 and PF4 (**Figure 2** and **Table 2**). The AUC of this 10-protein panel (the **MVA Panel**) was computed using logistic regression modeling, which reached 0.983 in distinguishing LD from controls at diagnosis, with 0.95 in sensitivity and 0.925 in specificity, respectively (**Figure 2A**). Using 2*10^4^ Monte Carlo cross-validation (MCCV) permutations with a 33% holdout rate, we assessed the performance of this **MVA Panel** and got an average AUC of 0.944. In fact, a combination of 4 proteins (ALDOB, CST6, FBP1 and PGLYRP2) from this 10-protein panel performed at a relatively high accuracy of 0.83 (0.85 sensitivity, 0.80 specificity) with an AUC = 0.92.

**Figure 2.**
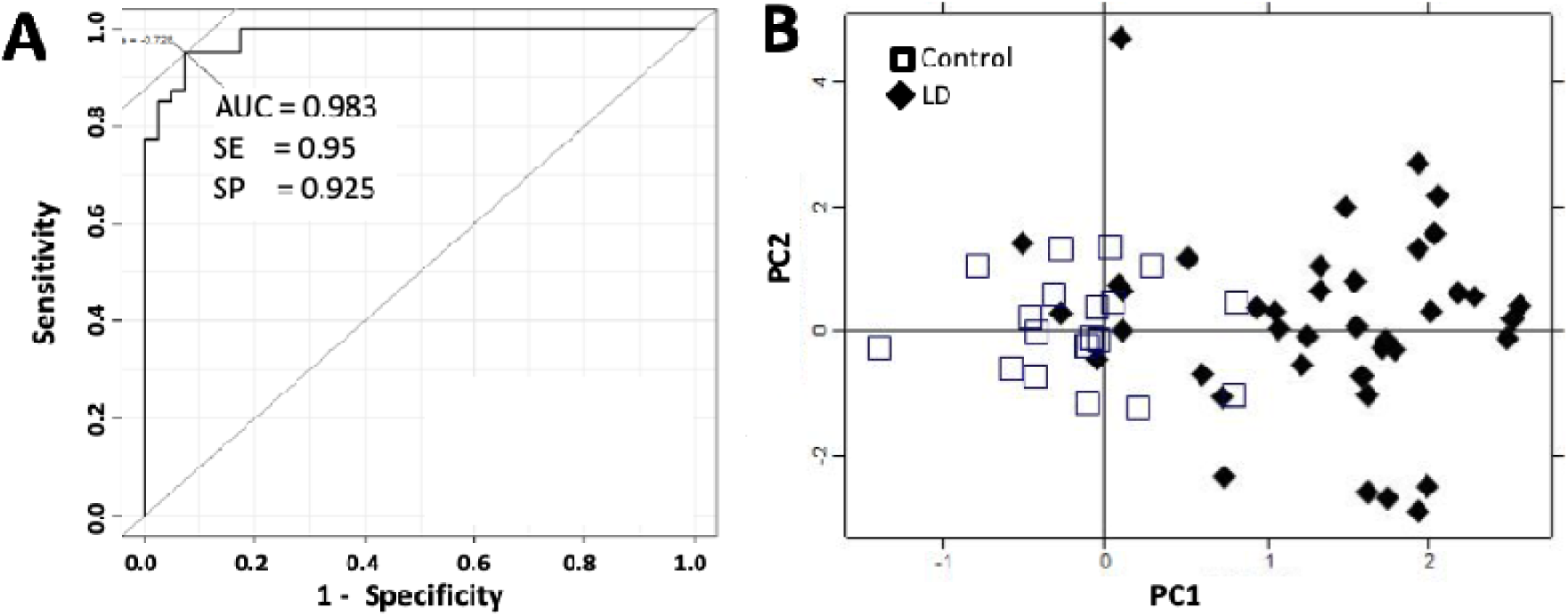
Performance of a group of 10 proteins discovered by multivariate analysis in the discovery cohort to identify patients with LD at baseline (MVA Panel). **2A**, AUC, and **2B**, Biplots of PC1 and PC2 scores of samples (40 LD patients and 20 healthy controls) from unsupervised PCA analyses, computed using the 10 **MVA Panel** proteins (11 proteotypic peptides). Filled diamonds and open squares represent PC scores for LD patients and controls, respectively.

Interestingly, four of these **MVA Panel** proteins (ALDOB, C9, CRP and CST6) overlapped with the ***t*-Test Set** of proteins, although the most cooperative proteins may not necessarily be the proteins with best individual performance identified by *t*-test ^35^. In unsupervised PCA analysis, similar to the 10 ***t*-Test Set** proteins, the PC scores of these 10 **MVA Panel** proteins presented a distinctive distribution between LD patients at baseline and healthy controls (**Figure 2B**).

**Figure S1A** illustrates the relative serum abundances of these 16 LD-associated proteins (10 ***t*-Test Set** proteins and 10 proteins in **MVA Panel**, with 4 overlapping), in LD individuals at baseline and in healthy controls.

### 2. Seronegative *versus* Seropositive Samples in Early Stage Lyme Disease

Next, we analyzed the performance of the 10 individual proteins of the ***t*-Test Set**, as well as the **MVA Panel**, in stratifying LD patients in seronegative (Sero-, n = 23) and seropositive (Sero+, n = 17) subgroups at the baseline visit.

Interestingly, all 10 individual proteins (13 proteotypic peptides) presented significant serum level changes in the Sero-subgroup *versus* controls (*t*-test *P* < 0.05, **Table S4**), with no significant difference between the 2 LD subgroups (*t*-test *P* > 0.05). The **MVA Panel** also returned similar AUC scores from multivariate analysis in both Sero- and Sero+ subgroups versus healthy controls (**Table S5**), performing slightly better in Sero-(AUC 0.911) compared to the Sero+ subgroup (AUC 0.865).

These findings suggest that these LD-associated serum proteins and the protein panel recognize acute LD in both seronegative and seropositive patients, when compared to healthy controls. If further validated, these proteins could potentially be used to aid the diagnosis of LD in patients with a negative serologic test result at the early stage of infection.

### 3. Protein Level Changes After Antimicrobial Treatment

At 4 weeks post-antimicrobial treatment, the perturbed serum levels of these 16 proteins returned to normal (fold change < 0.5 and *P* > 0.05 *vs*. healthy controls) in more than 80% of patients (**Figure S1B**), indicating diminished host immune responses and the reversibly acute effects in *B. burgdorferi* affected organs. This observation is consistent with multiple reports indicating that LD-associated hepatic dysfunction usually resolves within 2–3 weeks after starting treatment ^7-9^.

### 4. Network Analysis of Proteins Associated with Lyme Disease

We investigated the functional relationships among the 16 proteins discovered in this study and their potential involvement in response to *B. burgdorferi* infection and LD progression, using Gene Ontology (DAVID) and pathway-based (KEGG) analyses. As illustrated in **Figure S2**, many of the 16 proteins are involved in multiple extracellular pathways, including proteolysis, the immune response, and defense responses to bacteria. The remaining organ-specific proteins, for example, ALDOB (aldolase, fructose-bisphosphate B) and FBP1 (fructose-bisphosphatase 1), are members of intracellular metabolic pathways.

Pathways and functions related to innate immune reaction and acute phase response to pathogen infections are highly enriched (**Figure S2**). At least five proteins are directly involved in host responses to *B. burgdorferi* infection. For example, CRP (C-reactive protein), an acute reactant protein produced and secreted by hepatocytes, is a component of first line innate host defense, and serves as a sensitive signal of inflammation or infection ^42^. Two other proteins, C9 (complement component 9) and CFHR1 (complement factor H-related protein 1), are members of complement component cascades. C9, which can polymerize into complexes with C5, C6, C7, C8 or itself to neutralize a pathogen’s plasma membrane ^43^, also plays a key role in the adaptive immune response by enhancing the ability of antibodies and phagocytic cells to clear microbes. CFHR1 is a complement regulator and a member of factor H-related proteins that *B. burgdorferi* can bind to via outer surface proteins like *Erp*—an interaction that allows spirochetes to evade host complement during the initial stages of mammalian infection ^44^. PGLYRP2 (peptidoglycan recognition protein 2) belongs to the peptidoglycan recognition protein (PGRP) family, which recognizes and binds to peptidoglycan, an essential cell wall component of bacteria ^45^. It hydrolyzes the link between N-acetylmuramoyl residues and L-amino acid residues in peptidoglycan and may play a scavenger role by digesting biologically active peptidoglycan into biologically inactive fragments. S100A9 (S100 calcium-binding protein A9), a damage-associated molecular pattern molecule, is expressed specifically in monocytes and granulocytes. In association with its dimerization partner S100A8, S100A9 plays a prominent role in the regulation of inflammatory processes and immune response. Its differential expression has been associated with acute and chronic inflammation, and in sterile inflammatory conditions and cancers ^46^.

Whether the altered pattern of acute-phase related proteins discovered in this study is unique to *B. burgdorferi* infection, or may also reflect other types of infections, remains to be determined. We have observed that serum levels of complement factors C9 and CFHR1 changed significantly in LD at baseline, while some others, e.g., C4BPB, C5, C6, C8A/B/G, CFB, CFH, and CFP did not. The explanation for this behavior is not clear.

### 5. Organ-Specific Proteins Associated with Lyme Disease

Among the 16 LD-associated blood proteins, 13 are expressed predominantly in the liver. These 13 liver proteins consist of seven acute immune response proteins (C9, CFHR1, CRP, GC, ITIH2, ITIH4 and PGLYRP2), and four secreted proteins (AFM, APOA4, APOB and F9) that are involved in protein/lipid/polysaccharide transport or coagulation in the blood. The remaining two proteins (ALDOB and FBP1) are intracellular proteins involved in hepatocyte glycolysis/gluconeogenesis metabolism. Intracellular liver proteins are normally present in blood at low concentrations due to leakage. Blood levels of secreted liver proteins may decline under disease conditions due to disturbed protein production and secretion caused by organ injuries or infection. We have found that the elevation of intracellular proteins and the reduction of extracellular proteins in the blood may serve as sensitive markers of tissue injury or disease-perturbed tissue function in specific organs ^25^.

CST6 (Cystatin E/M) is the only skin-specific protein in our acute LD-associated protein list. It is secreted specifically from differentiated epidermal keratinocytes and serves as a skin-specific proteinase inhibitor. Cystatins and cystatin-like molecules belong to a new category of immunomodulatory molecules that act as inhibitors of lysosomal cysteine proteases, i.e., the cathepsins. Cathepsins are involved in processing and presentation of antigens, and have been associated with pathological conditions such as inflammation and cancers ^47^. An imbalance between cathepsins and cystatins may attenuate immune cell functions and thus facilitate tumor cell invasion ^48^. Decreased CST6 levels have been observed in inflamed skin ^49, 50^, which may reflect inflammatory processes in the skin area affected by EM in early LD patients.

We wish to emphasize the power of monitoring the concentration of organ-specific proteins in the blood to identify organs that have been perturbed in the course of a disease. We envision a time when at least 10 or more organ-specific proteins for the 25 most common organs will be in-hand—thus affording the ability, for a given disease, to determine which organs have been affected throughout the course of that disease. Examination of the disease-perturbed networks related to these sets of organ-specific proteins also provides insights into the pathophysiology of the disease. A key to increasing the number of detectable organ-specific blood proteins may lie in the ability to enrich and hence quantify low-abundance blood proteins (see discussion below).

### 6. Verification of SRM Results by Western Blotting

To verify the SRM results, we performed Western blot analysis with antibodies against four candidate proteins: ALDOB, CRP, CST6, and FBP1. For each protein, we measured serum levels in four randomly chosen LD patients (with 4 time points) and four healthy controls (2 time points) from the same cohort. Characteristic bands for ALDOB and CRP were observed, whereas antibodies against CST6 and FBP1 did not work well presumably due to their low antigen abundance in serum. Western blot results of both ALDOB and CRP agreed with SRM results, with Pearson’s correlation coefficient *r* = 0.83 for CRP and *r* = 0.80 for ALDOB (representative Western blot results are shown in **Figure S3**).

### 7. Verification of LD-associated Proteins in an Independent LD Cohort

We further evaluated the levels of these 16 LD-associated proteins by SRM in an independent LD cohort from NYMC, as described in Materials and Methods. This verification cohort included 30 patients with symptoms in addition to EM rash at diagnosis. Demographic and clinical characteristics of the NYMC LD patients are similar to those of the discovery SLICE cohort (**Table S1**).

Sample preparation, the SRM runs and data analyses in the verification cohort were exactly the same as for the discovery cohort. Fourteen proteins (18 peptides) out of the 16 LD-associated proteins were successfully detected and measured. The abundances of the remaining two proteins, ALDOB and FBP1, were too low to be reliably quantified in most samples, and were consequently removed from statistical analysis. Coagulation Factor IX (F9), a protein involved in the blood coagulation cascade, was also eliminated from further statistical analysis due to its abnormally low serum concentrations (∼50% lower) in all LD samples compared to healthy controls. A heatmap of the 16 quantified proteins in LD patients over the 4 time points is presented in **Figure S4**.

#### Six early LD-associated serum proteins were verified

Six of the eight quantified proteins (APOA4, C9, CRP, CST6, PGLYRP2 and S100A9) in the ***t*-Test Set** were successfully verified in the second verification patient cohort (Student’s *t*-test *P* < 0.05). Similar cut-off levels were identified by pAUC, with similar sensitivities and specificities as in the discovery cohort (**Table 4**). Two proteins, AFM and ITIH2, had no power for discriminating LD from healthy controls and could not be verified.

**Table 4:** LD patients identified at baseline by the six individual proteins in the verification cohort.

| Protein<br>Peptide | APOA4<br>LGPHAGDVEGHLSFLEK | C9<br>LSPIYNLVPVK | CRP<br>ESDTSYVSLK | CST6<br>AQSQLVAGIK | PGLYRP2<br>EFTEAFLGCPAIHPR | S100A9<br>LGHPDTLNQGEFK |
| --- | --- | --- | --- | --- | --- | --- |
| *P* value (*t*-test) | 2.5E-05** | 3.0E-03** | 5.0E-04** | 4.0E-02* | 7.0E-03* | 4.0E-03** |
| FC cutoff | < 0.33 | > 1.6 | > 4.5 | < 0.60 | < 0.35 | > 2.2 |
| *Partial AUC at cutoff* |  |  |  |  |  |  |
| sensitivity | 40.0% | 41.2% | 64.7% | 24.3% | 22.9% | 27.0% |
| specificity | 92.5% | 92.5% | 97.5% | 90.0% | 97.5% | 92.5% |
| *# of patients detected at cutoff* |  |  |  |  |  |  |
| Lyme (n=30) | 8 (26.7%) | 12 (40.0%) | 22 (73.3%) | 8 (26.7%) | 6 (20.0%) | 11 (36.7%) |
| *Average FC to controls* |  |  |  |  |  |  |
| Sero - | 0.42 ± 0.29 | 1.61 ± 0.98 | 24.47 ± 36.86 | 0.81 ± 0.37 | 0.78 ± 0.31 | 1.69 ± 0.95 |
| Sero + | 0.57 ± 0.47 | 1.59 ± 0.73 | 14.00 ± 15.74 | 0.83 ± 0.4 | 0.81 ± 0.38 | 1.76 ± 1.36 |
| Healthy control | 1 ± 0.6 | 1 ± 0.43 | 1 ± 1.17 | 1 ± 0.32 | 1 ± 0.41 | 1 ± 0.75 |
| *t-test P value* |  |  |  |  |  |  |
| Sero - vs. Control | 1.4E-05** | 3.2E-02* | 2.2E-02* | 7.8E-02* | 4.3E-02* | 1.6E-02* |
| Sero + vs. Control | 9.2E-03* | 1.1E-02* | 4.7E-03** | 1.4E-01 | 1.1E-01 | 5.4E-02 |
| Sero + vs. Sero - | 3.3E-01 | 9.4E-01 | 3.1E-01 | 8.7E-01 | 7.9E-01 | 8.7E-01 |

\* cutoff value is the fold change to average of healthy controls in the same sample set.
Sero-: LD patient group with negative first-tier serology test (EIA) at diagnosis
Sero+: LD patient group with positive serology test at diagnosis. \*: *t*-test $P < 0.05$ , \*\*: $P < 0.005$ .

The combined ROC of these six verified proteins was calculated using logistic regression modeling, which reached an AUC = 0.893 in 10-fold CV (whole set = 0.925), with 0.88 in sensitivity, 0.83 in specificity and 0.90 in accuracy, all scores slightly lower than in the discovery cohort (**Figure 3A**). At the prediction probability cutoff value of 0.696, the combination of these six proteins correctly predicted 23 out of the 30 LD patients in the verification cohort (**Figure 3B**).

**Figure 3.**
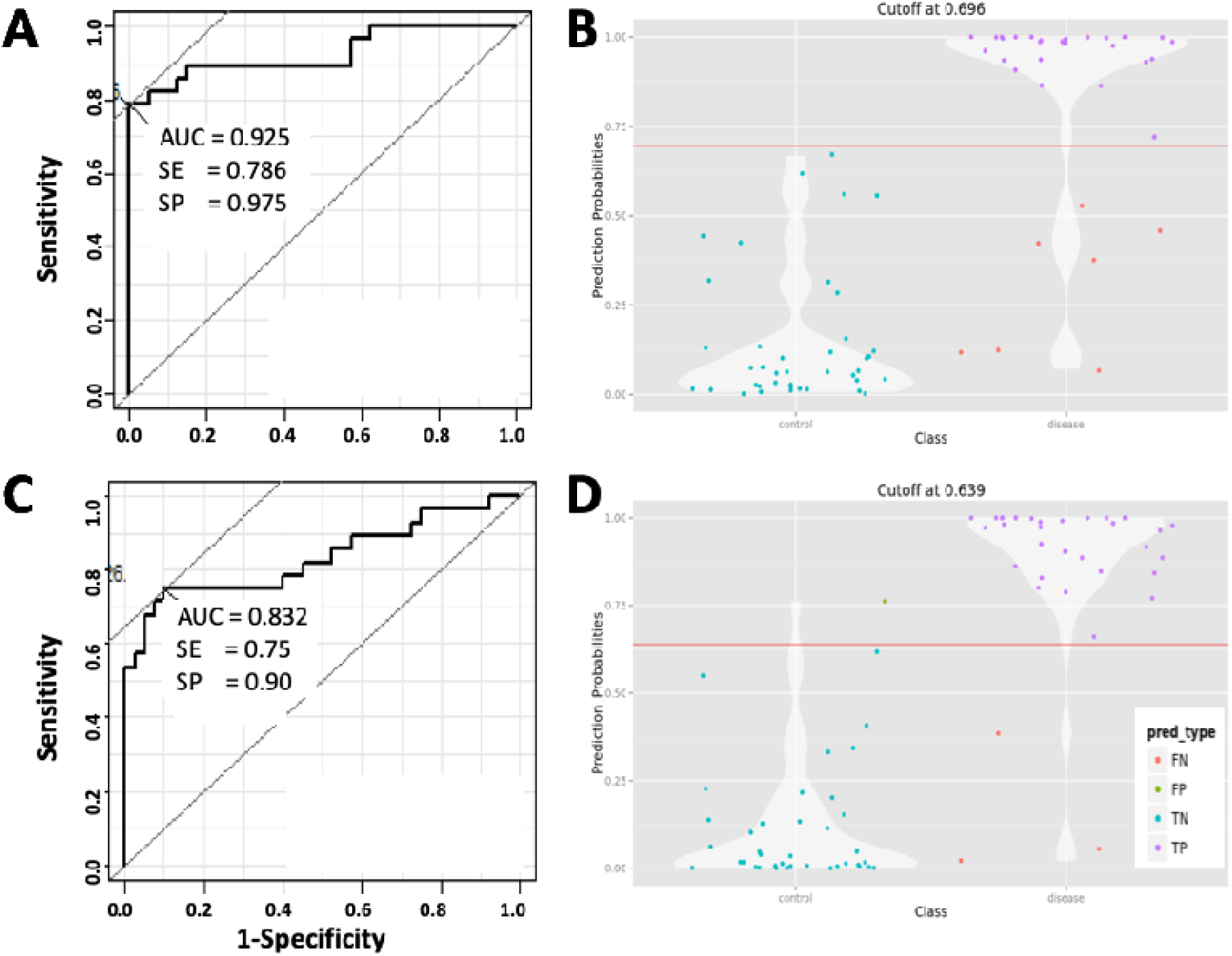
Serum levels of early Lyme disease associated proteins in the verification cohort. **3A**, Combined AUC (whole set), sensitivity and specificity of the 6 verified blood proteins showing statistically altered serum levels in LD at the baseline time point. **3B**, Violin plot combines a box plot and a kernel density plot, showing the probability density to classify LD patients by the six proteins at the indicated cutoff value of AUC. The four possible categories: False Negative (FN); False Positives (FP); True Negative (TN); True Positive (TP). **3C**, AUC, sensitivity and specificity of the 9-protein panel identified by multivariate analysis. **3D**, Violin plot showing the probability density to classify LD patients by the 9-protein panel at the indicated cutoff level of AUC.

#### The 10-protein panel from multivariate analysis performs well in verification cohort

The 10-protein **MVA Panel** discovered by MVA was also verified. As discussed in the previous section, F9 was removed because of its abnormally low concentrations in LD samples which seemed likely to be technical error, thus artificially increasing accuracy as a result. Subsequently, the ROC of the 9-protein panel was computed using the model “locked-down” from the discovery cohort, except the contributions of F9 were set as zero in the formula. This model resulted in an AUC = 0.832 (sensitivity 0.75, specificity 0.9 and accuracy 0.84) (**Figure 3C**). Interestingly, serum levels of two proteins, APOB (*p* < 0.001) and CFHR1 (*p* = 0.05), were also significantly increased in this verification cohort, in contrast to results in the discovery cohort. As illustrated in **Figure 3D**, at a cutoff value of 0.639, this 9-protein panel correctly classified 27 out of the 30 LD patients in the verification cohort with one false-positive among the 20 controls.

#### Seronegative versus seropositive samples

As in the first cohort, all six verified individual proteins from the ***t*-Test Set** demonstrated significant serum level changes in Sero-patients (n=16) *versus* healthy controls (n=20, *t* test *P* < 0.05, **Table 4**), with no significant difference between the 2 LD subgroups (*t* test *P* > 0.05, **Table 4**). The **MVA Panel** (nine proteins) also reached similar AUC scores from multivariate analysis in both Sero-and Sero+ (n=14) subgroups *versus* healthy controls (**Table S5**), performing slightly better in Sero-(AUC 0.904) compared to the Sero+ subgroup (AUC 0.842).

As shown in **Figure S4**, after an average of 11.9 ± 5.9 days of antimicrobial treatment, the perturbed serum levels of these proteins returned to normal, similar to the discovery samples.

It should be noted that, as most of the patient samples in this cohort were collected in the 1990s, some of the candidate proteins could have degraded during storage, which may explain why they were apparently below the level of detection in this verification sample set.

### 8. Limitations of the Study

The major limitation of this study is the relatively small sample size (70 LD and 40 healthy controls) from two cohorts. Thus, findings herein should not be considered as biomarkers, but as candidates that will require further validation in appropriately controlled studies. Also, the possibility of misdiagnosed patients in those without microbiologic confirmation, and the different levels of serology tests performed in the two cohorts, could impact the observed outcomes. We therefore plan to validate these acute LD-associated blood proteins in additional sample sets, ultimately aiming to develop a diagnostic protein panel for detection of early LD, prediction of disease progression, and assessment of the overall health of specific organ systems in patients who have persistent symptoms seemingly related to LD. We also need to evaluate our candidate proteins in blood from patients that have other types of infections as well to determine how specific these markers are for LD. That said, in geographic areas with high rates of LD, these blood proteins, if validated, could prove useful in informing early treatment with antibiotics in seronegative patients without an EM rash.

Another limitation of this study is that most organ-specific proteins that can be detected and quantified in depleted serum by LC-MS-SRM are from several large organs like the liver. It is still challenging to detect proteins originating from small organs (thus presumably contributing lower levels of proteins to the blood) or organs isolated from systemic blood flow, e.g., via the blood brain barrier. These less detectable proteins could be very informative to the study of many diseases. One approach to investigating these very low-abundance organ-specific proteins in the blood is to specifically capture them using a multiplex affinity column of antibodies or other types of protein or peptide binding reagents. In theory, such reusable positive enrichment methods could improve our ability to study dozens of target proteins present at very low levels in the blood from specific organs that are involved in early or late manifestations of *B. burgdorferi* infections. Equalizing approaches to reduce high-abundance proteins and enrich low-abundance proteins in biological liquids, for example, ProteoMiner hexapeptide beads, could also enhance our ability to detect and measure low-abundant proteins in blood secreted from particularly small organs ^51, 52^. We plan to quantify and assess additional organ-specific proteins, especially proteins enriched in other *B. burgdorferi*-affected organs such as the bladder, brain, heart, and muscle, with improved sample enrichment procedures.

## CONCLUSIONS

In this study, we focused on specific blood proteins drawn from two key categories related to *B. burgdorferi* infection: 1) proteins involved in the host innate immune response during the acute phase of infection and 2) proteins specifically originating from single organs that are possible targets of the infection, for example, liver, brain, heart and skin. With this focused approach we aimed to identify serum proteins whose altered blood abundances, relative to healthy controls, could identify individuals infected with *B. burgdorferi* at the earliest stage possible.

In two independent LD cohorts, we identified a set of proteins that may fulfill that aim:

1. In the discovery cohort, we identified 10 proteins by *t*-test (ALDOB, AFM, APOA4, C9, CRP, CST6, FBP1, ITIH2, PGLYRP2 and S100A9), each of which has significantly altered protein levels in serum and is capable of distinguishing LD patients from healthy controls at the time of diagnosis. We have also identified an **MVA Panel** of 10 proteins (ALDOB, APOB, C9, CFHR, CRP, CST6, GC, F9, ITIH4 and PF4) by multivariate analysis that distinguished LD patients from healthy controls with high accuracy.
2. In an independent LD cohort, we verified i) 6 proteins from the ***t*-Test Set** of 10 proteins, and ii) 9 of the 10 proteins in the **MVA Panel**, with similar sensitivity and specificity as in the discovery cohort.
3. These LD-associated serum proteins (***t*-Test Set**) and protein panel (**MVA Panel**) identified acute LD in both seronegative and seropositive patients.

We plan to further validate these early LD-associated serum proteins by testing additional independent LD and healthy control cohorts, along with other appropriate disease control cohorts.

## AUTHOR INFORMATION

### Author Contributions

The manuscript was written through contributions of all authors. All authors have given approval to the final version of the manuscript.

## ACKNOWLEDGMENTS

This work was supported by Bay Area Lyme Foundation (www.bayarealyme.org), The Steven and Alexandra Cohen Foundation (http://www.steveandalex.org), NIH grant P30AR05350 (to JNA and MJS), P50GM076547 (LH and RLM), the National Institute of Allergy and Infectious Diseases under Award Number R21AI133335 (RLM), and The Wilke Family Foundation. GSO acknowledges NIH grants P30ES017885 and U24CA210967. The authors would like to thank the individuals with LD and healthy controls who participated in this study. We thank our colleagues at ISB: Dr. Simon Evans who helpfully read and commented on the manuscript, and Dr. Max Robinson for helpful discussions on the statistics and analytical aspects of this study.

## FUNDING SOURCES

Bay Area Lyme Foundation (www.bayarealyme.org)

The Steven and Alexandra Cohen Foundation (http://www.steveandalex.org)

The Wilke Family Foundation

NIH P50GM076547 (LH and RLM)

NIH grant P30AR05350 (to JNA and MJS)

the National Institute of Allergy and Infectious Diseases R21AI133335 (RLM)

NIH grants P30ES017885 and U24CA210967 GSO acknowledges

## ABBREVIATIONS

AFM: afamin;
ALDOB: aldolase B,
AUC: area under the curve;
C9: complement component 9;
CFHR1: complement factor H-related protein 1;
CRP: C-reactive protein;
CST6: cystatin E/M;
F9: Coagulation Factor IX;
FBP1: fructose-bisphosphate 1;
GC: vitamin D-binding protein-macrophage activating factor;
ITIH2: inter-alpha (globulin) inhibitor H2;
ITIH4: inter-alpha-trypsin inhibitor 4;
LD: Lyme disease;
MVA: multivariate analysis;
MS: mass spectrometry;
pAUC: partial AUC;
PF4: platelet factor 4;
PGLYRP2: peptidoglycan recognition protein 2;
ROC curve: Receiver operating characteristic curve;
SRM: selective reaction monitoring

## Supporting Information

### 1. Supplemental Figures

**Figure S1. A)** Heatmap of serum levels of the 16 LD-associated blood proteins identified in the discovery cohort with 40 LD (LD01 – LD40) and 20 healthy controls (C01 – C20) at the baseline time point. **B)** Heatmap of average serum levels of the 16 LD-associated blood proteins identified in the discovery cohort over the 4 time points from baseline to 12 months post-treatment (**Lyme 1**, baseline; **Lyme 2**, 4-wk post-treatment; **Lyme 3**, 6-mo post-treatment; and **Lyme 4**, 12-mo post-treatment). The log_2_ fold changes against average of healthy controls are shown. Red: up, Blue: down, compared to average of controls. Proteins with two peptides: *ApoA4_L*, LGPHAGDVEGHLSFLEK; *ApoA4_S*, SELTQQLNALFQDK; *CRP_E*, ESDTSYVSLK; *CRP_G*, GYSIFSYATK; *PGLYRP2_E*, EFTEAFLGCPAIHPR, *PGLYRP2_G*, GCPDVQASLPDAK.

**Figure S2**. Network analysis of 16 early LD-associated proteins revealed by *t*-test and multivariate analysis in the discovery cohort.

The intracellular pathway glycolysis/gluconeogenesis, extracellular pathways of proteolysis, immune response, and defense response to bacteria are highly enriched among these proteins. Node colors: RED, up in LD serum; BLUE, down in LD serum; Green, varies; GRAY: proteins not measured in this study.

**Figure S3**. Western blot verification of LD-associated blood proteins identified in LC-MS-SRM analysis. **2A**, ALDOB; **2B**, CRP. The patient IDs, time points post-diagnosis (TP) and patient groups are labeled on the bottom of gel images. The average abundance of healthy controls = 1 (dot line).

**Figure S4**. Verification of LD-associated proteins in the second independent Lyme disease cohort. Heatmap shows the serum levels of the 16 proteins measured in the 30 Lyme disease patients in this verification cohort by LC-MS-SRM. Average fold changes in serum abundance are shown. Time points: **Lyme 1**, baseline; **Lyme 2**, convalescence; **Lyme 3**, one-year post-treatment; and **Lyme 4**, 4-6 years post-treatment. Red: up, Blue: down, relative to average of healthy controls (**Cntl 1**: at the initial visit, and **Cntl 2**: one year later).

### 2. Supplemental Tables

(file name: YZhou etal_Lyme-associated protein_Suppl tables.xlsx)

**Table S1**. Detailed demographic and clinical characteristics in both SLICE (JHU) and New York Medical College (NYMC) sample sets

**Table S2**. Sample distributions in two LD cohorts

**Table S3**. Summarization of SRM methods for 174 monitored proteotypic peptides

**Table S4**. Serum level changes of the 10 individual proteins (***t*-Test Set**) in seronegative and seropositive subgroups in the SLICE LD cohort

**Table S5**. The performance of **MVA Panel** in stratifying seronegative and seropositive LD subgroups from healthy controls in both SLICE and NYMC sets

## TOC

Organ-specific & Acute-phase Blood Proteins in Early Lyme Disease

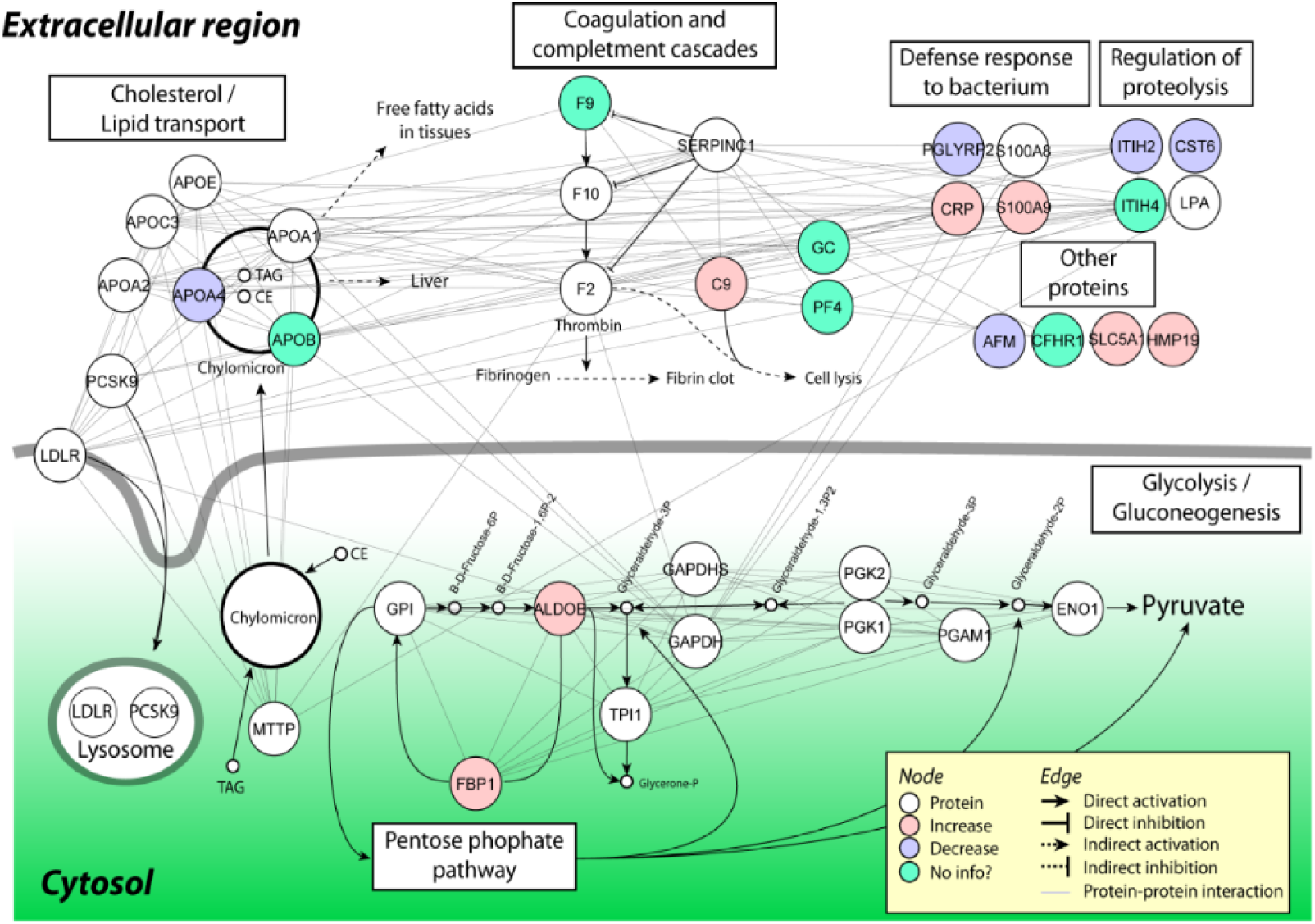

