## Supplemental Figs and Tables for "Measurement of organ-specific and acute-phase blood protein levels in early Lyme disease"

**Table of contents**

**1. Supplemental Figures:**

**Figure S1.** Heatmap of serum levels of the 16 LD-associated blood proteins identified in the discovery cohort

**Figure S2.** Network analysis of 16 early LD-associated proteins revealed by *t*-test and multivariate analysis in the discovery cohort.

**Figure S3.** Western blot verification of LD-associated blood proteins identified in LC-MS-SRM analysis.

**Figure S4.** Verification of LD-associated proteins in the second independent Lyme disease cohort.

**2. Supplemental Tables:** (Also in file: YZhou etal_blood proteins in early Lyme_Suppl tables.xlsx)

**Table S1.** Detailed demographic and clinical characteristics in both SLICE (JHU) and New York Medical College (NYMC) sample sets

**Table S2.** Sample distributions in two LD cohorts

**Table S3.** Summarization of SRM methods for 174 monitored proteotypic peptides

**Table S4.** Serum level changes of the 10 individual proteins (***t*-Test Set**) in seronegative and seropositive subgroups in the SLICE LD cohort

**Table S5.** The performance of **MVA Panel** in stratifying seronegative and seropositive LD subgroups from healthy controls in both SLICE and NYMC sets

### SUPPLEMENTAL FIGURES

**Figure S1**. **A)** Heatmap of serum levels of the 16 LD-associated blood proteins identified in the discovery cohort with 40 LD (LD01 – LD40) and 20 healthy controls (C01 – C20) at the baseline time point (i.e., at LD diagnosis). **B)** Heatmap of average serum levels of the 16 LD-associated blood proteins identified in the discovery cohort over the four time points from baseline to 12 months post-treatment in patients (**Lyme 1**, Baseline; **Lyme 2**, 4-wk post-treatment; **Lyme 3**, 6-mo post-treatment; and **Lyme 4**, 12-mo post-treatment). The $\log_2$ fold changes relative to the average of controls at baseline are shown. Red: up, Blue: down, relative to average of healthy controls (**Cntl 1**, at the initial visit; and **Cntl 2**, 6 months later). Proteins with two peptides: *ApoA4_L*, LGPHAGDVEGHLSFLEK; *ApoA4_S*, SELTQQLNALFQDK; *CRP_E*, ESDTSYVSLK; *CRP_G*, GYSIFSYATK; *PGLYRP2_E*, EFTEAFLGCPAIHPR, *PGLYRP2_G*, GCPDVQASLPDAK.

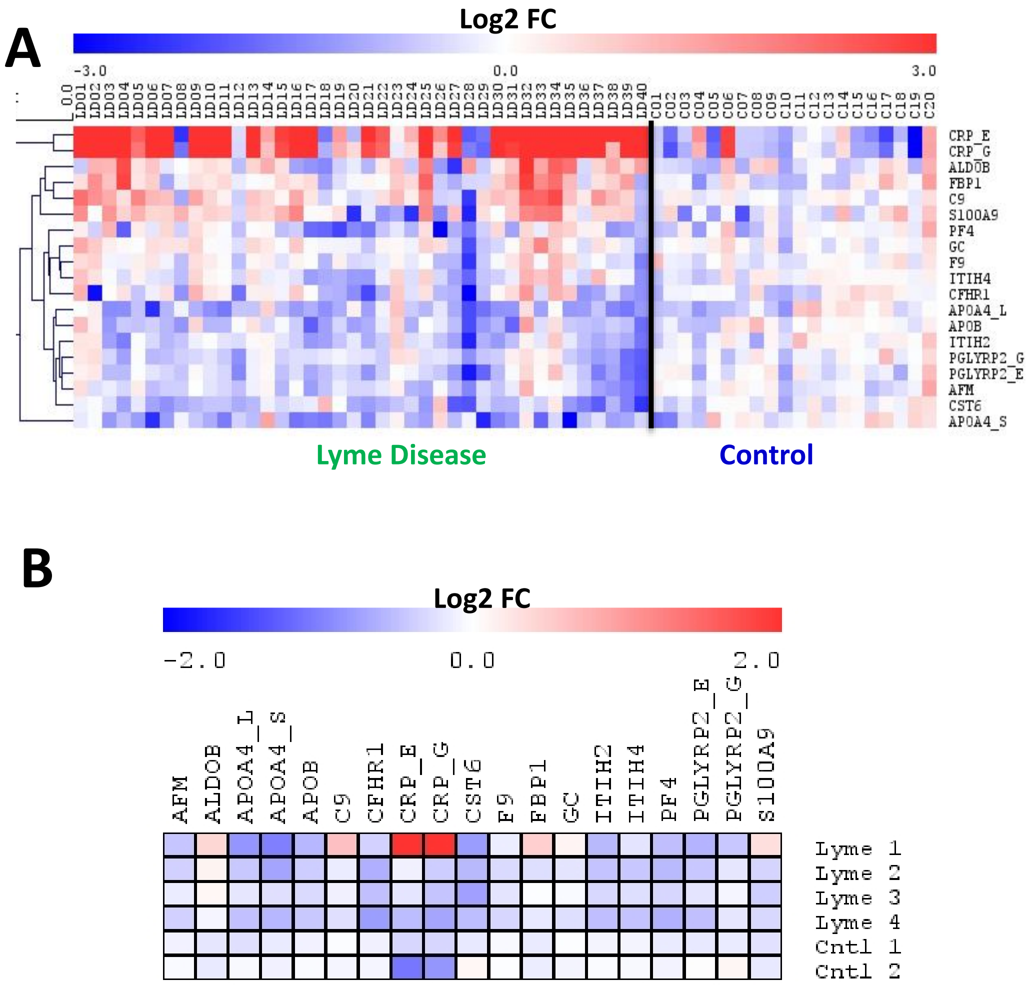
A
Log2 FC
-3.0
0.0
3.0
CRP_E
CRP_G
ALDOB
FBP1
C9
S100A9
PF4
GC
F9
ITIH4
CFHR1
APOA4_L
APOB
ITIH2
PGLYRP2_G
PGLYRP2_E
AFM
CST6
APOA4_S
Lyme Disease
Control
B
Log2 FC
-2.0
0.0
2.0
AFM
ALDOB
APOA4_L
APOA4_S
APOB
C9
CFHR1
CRP_E
CRP_G
CST6
F9
FBP1
GC
ITIH2
ITIH4
PF4
PGLYRP2_E
PGLYRP2_G
S100A9
Lyme 1
Lyme 2
Lyme 3
Lyme 4
Cntl 1
Cntl 2

**Figure S2.** Network analysis of 16 early LD-associated proteins revealed by *t*-test and multivariate analysis in the discovery cohort.

The intracellular pathway glycolysis/gluconeogenesis, extracellular pathways of proteolysis, immune response, and defense response to bacteria are highly enriched among these proteins. Node colors: RED, up in LD serum; BLUE, down in LD serum; Green, varies; GRAY: proteins not measured in this study.

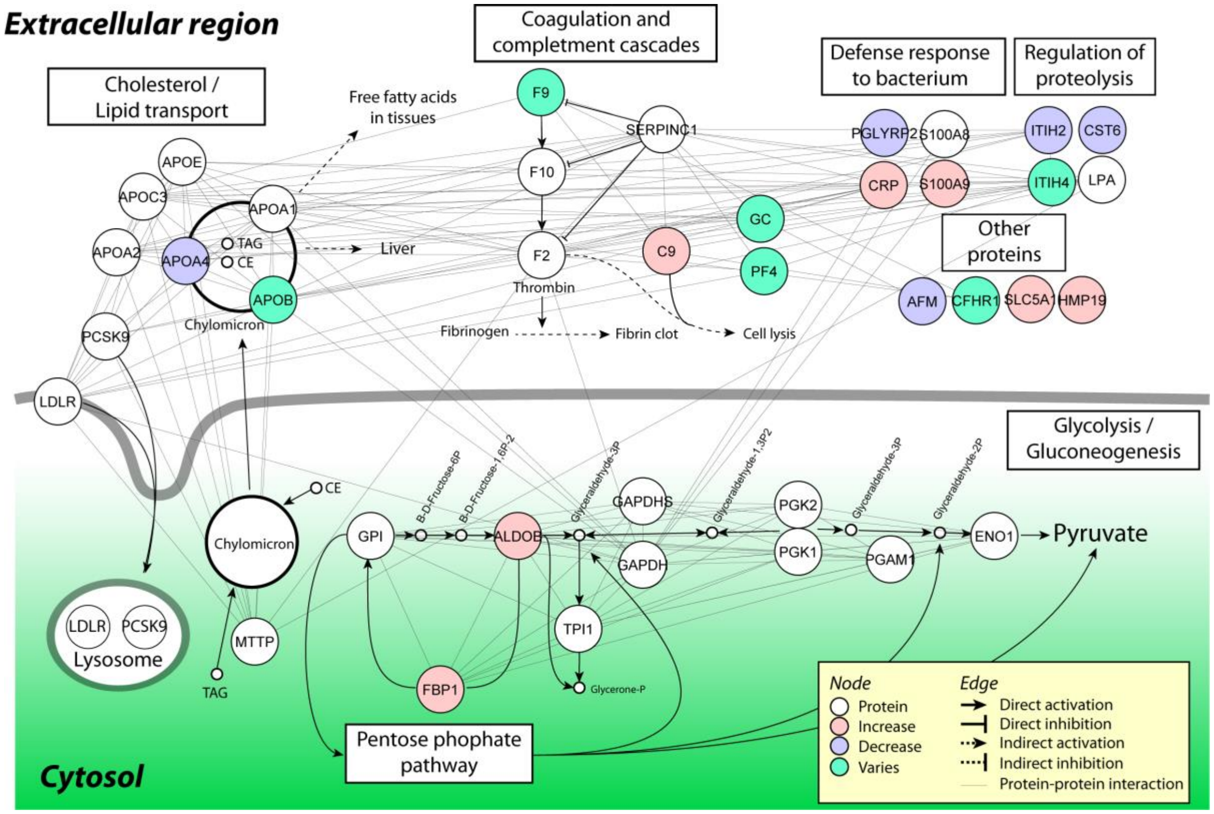

**Figure S3.** Western blot verification of *B. burgdorferi* infection affected proteins identified from SRM analysis in selected sera from the discovery cohort. **A**, ALDOB; **B**, CRP. The patient IDs, time points post-diagnosis (TP) and patient groups are labeled on the bottom of gel images. The average of healthy controls = 1 (dot line).

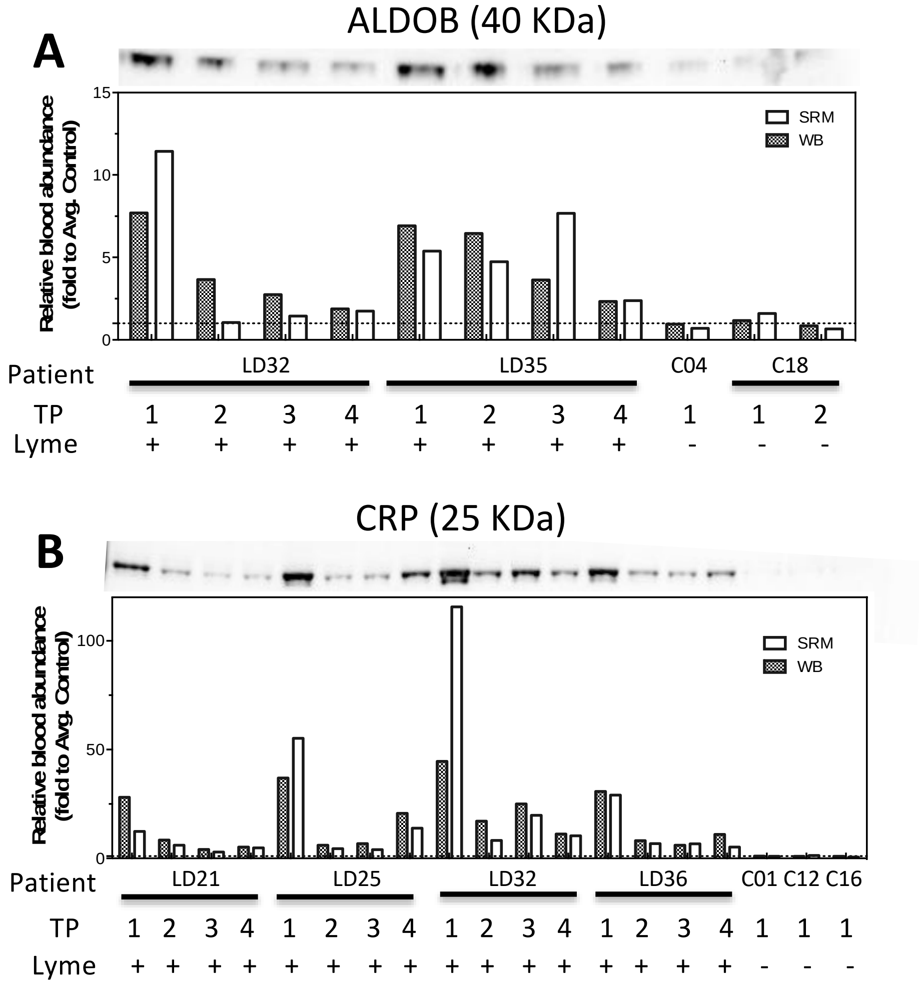

**Figure S4.** Verification of candidate biomarkers in the second independent Lyme disease cohort. Heatmap shows the serum levels of the 16 proteins measured in the 30 Lyme disease patients in this verification cohort by LC-MS-SRM. Average fold changes in serum abundance are shown for LD patients. Time points: **Lyme 1**, baseline; **Lyme 2**, convalescence; **Lyme 3**, one-year post-treatment; and **Lyme 4**, 4-6 years post-treatment. Red: up, Blue: down, relative to average of healthy controls (**Cntl 1**, at the initial visit; and **Cntl 2**, one year later).

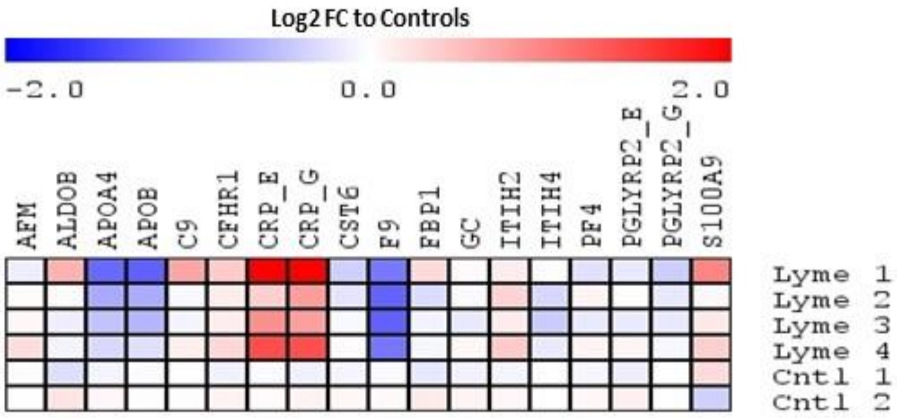

**Table S1.** Detailed demographic and clinical characteristics in both SLICE (JHU) and New York Medical College (NYMC) sample sets

| ISB PID | Lyme Borreliosis | AGE | GENDER | RASHSIZE | DISSEM | SEROGRP | DURAT |
|---|---|---|---|---|---|---|---|
| *SLICE (JHU)* | | | | | | | |
| LD01 | 1 | 54 | 1 | 160 | 0 | 1 | 4 |
| LD02 | 1 | 57 | 1 | 120 | 0 | 1 | 3 |
| LD03 | 1 | 20 | 1 | 280 | 0 | 2 | 4 |
| LD04 | 1 | 60 | 0 | 88 | 0 | 1 | 5 |
| LD05 | 1 | 61 | 1 | 208 | 1 | 1 | 10 |
| LD06 | 1 | 71 | 0 | 161 | 1 | 2 | 4 |
| LD07 | 1 | 51 | 0 | 256 | 0 | 2 | 13 |
| LD08 | 1 | 38 | 1 | 180 | 1 | 2 | 14 |
| LD09 | 1 | 53 | 0 | 144 | 0 | 2 | 5 |
| LD10 | 1 | 55 | 1 | 49 | 0 | 1 | 8 |
| LD11 | 1 | 63 | 0 | 49 | 0 | 0 | 14 |
| LD12 | 1 | 43 | 0 | 153 | 0 | 2 | 21 |
| LD13 | 1 | 33 | 1 | 495 | 0 | 2 | 28 |
| LD14 | 1 | 69 | 1 | 270 | 0 | 0 | 10 |
| LD15 | 1 | 35 | 1 | 32 | 0 | 1 | 3 |
| LD16 | 1 | 44 | 0 | 144 | 0 | 2 | 2 |
| LD17 | 1 | 26 | 1 | 225 | 1 | 1 | 4 |
| LD18 | 1 | 29 | 1 | 36 | 0 | 0 | 2 |
| LD19 | 1 | 67 | 1 | 324 | 1 | 2 | 14 |
| LD20 | 1 | 52 | 0 | 25 | 0 | 0 | 7 |
| LD21 | 1 | 64 | 0 | 135 | 0 | 1 | 3 |
| LD22 | 1 | 54 | 0 | 30 | 0 | . | 35 |
| LD23 | 1 | 69 | 0 | 99 | 1 | 0 | 8 |
| LD24 | 1 | 26 | 1 | 20 | 0 | 0 | 3 |
| LD25 | 1 | 43 | 1 | 52 | 0 | 1 | 3 |
| LD26 | 1 | 37 | 0 | 25 | 1 | 2 | 8 |
| LD27 | 1 | 38 | 0 | 21 | 0 | 0 | 4 |
| LD28 | 1 | 53 | 0 | 21 | 1 | 0 | 42 |
| LD29 | 1 | 20 | 1 | 551 | 0 | 1 | 7 |
| LD30 | 1 | 36 | 1 | 154 | 1 | 2 | 14 |
| LD31 | 1 | 68 | 1 | 195 | 1 | 2 | 10 |
| LD32 | 1 | 50 | 0 | 54 | 1 | 2 | 11 |
| LD33 | 1 | 42 | 1 | 112 | 0 | 1 | 3 |
| LD34 | 1 | 45 | 1 | 132 | 0 | 2 | 14 |
| LD35 | 1 | 56 | 0 | 289 | 1 | 2 | 10 |
| LD36 | 1 | 50 | 0 | 90 | 0 | 1 | 5 |
| LD37 | 1 | 49 | 0 | 77 | 1 | 2 | 10 |
| LD38 | 1 | 59 | 0 | 35 | 0 | 2 | 5 |
| LD39 | 1 | 64 | 0 | 150 | 0 | 1 | 4 |
| LD40 | 1 | 49 | 0 | 176 | 0 | 0 | 7 |
| C01 | 0 | 70 | 1 | N/A | N/A | 0 | N/A |
| C02 | 0 | 61 | 0 | N/A | N/A | 0 | N/A |
| C03 | 0 | 50 | 0 | N/A | N/A | 0 | N/A |
| C04 | 0 | 50 | 0 | N/A | N/A | 0 | N/A |
| C05 | 0 | 22 | 1 | N/A | N/A | 0 | N/A |
| C06 | 0 | 40 | 0 | N/A | N/A | 0 | N/A |
| C07 | 0 | 46 | 1 | N/A | N/A | 0 | N/A |
| C08 | 0 | 65 | 0 | N/A | N/A | 0 | N/A |
| C09 | 0 | 54 | 0 | N/A | N/A | 0 | N/A |
| C10 | 0 | 60 | 0 | N/A | N/A | 0 | N/A |
| C11 | 0 | 73 | 1 | N/A | N/A | 0 | N/A |
| C12 | 0 | 70 | 1 | N/A | N/A | 0 | N/A |
| C13 | 0 | 72 | 0 | N/A | N/A | 0 | N/A |
| C14 | 0 | 55 | 1 | N/A | N/A | 0 | N/A |
| C15 | 0 | 51 | 1 | N/A | N/A | 0 | N/A |
| C16 | 0 | 63 | 0 | N/A | N/A | 0 | N/A |
| C17 | 0 | 66 | 1 | N/A | N/A | 0 | N/A |

| | | | | | | | |
|---|---|---|---|---|---|---|---|
| C18 | 0 | 35 | 0 | N/A | N/A | 0 | N/A |
| C19 | 0 | 25 | 1 | N/A | N/A | 0 | N/A |
| C20 | 0 | 69 | 1 | N/A | N/A | 0 | N/A |
| *NYMC* | | | | | | | |
| B01 | 1 | 55 | 0 | 1442 | 0 | 2 | |
| B02 | 1 | 47 | 0 | 327 | 0 | 1 | |
| B03 | 1 | 50 | 0 | 544 | 0 | 0 | |
| B04 | 1 | 63 | 0 | 265 | 0 | 1 | |
| B05 | 1 | 53 | 0 | 1585 | 1 | 1 | |
| B06 | 1 | 26 | 0 | 236 | 0 | 0 | |
| B07 | 1 | 55 | 1 | 214 | 0 | 0 | |
| B08 | 1 | 64 | 0 | 427 | 0 | 0 | |
| B09 | 1 | 49 | 1 | 864 | 0 | 2 | |
| B10 | 1 | 42 | 0 | 283 | 1 | 2 | |
| B11 | 1 | 70 | 1 | 377 | 0 | 0 | |
| B12 | 1 | 54 | 0 | 1357 | 0 | 1 | |
| B13 | 1 | 37 | 1 | 132 | 0 | 2 | |
| B14 | 1 | 40 | 0 | 2727 | 0 | 2 | |
| B15 | 1 | 50 | 1 | . | 0 | 0 | |
| B16 | 1 | 55 | 0 | 679 | 1 | 1 | |
| B17 | 1 | 55 | 1 | 199 | 0 | 0 | |
| B18 | 1 | 45 | 1 | 191 | 1 | 0 | |
| B19 | 1 | 45 | 0 | 492 | 0 | 1 | |
| B20 | 1 | 47 | 0 | 2268 | 0 | 2 | |
| B21 | 1 | 57 | 0 | 2790 | 1 | 2 | |
| B22 | 1 | 65 | 1 | 1382 | 1 | 2 | |
| B23 | 1 | 32 | 0 | 88 | 0 | 1 | |
| B24 | 1 | 49 | 1 | 542 | 1 | 2 | |
| B25 | 1 | 72 | 0 | 132 | 0 | 1 | |
| B26 | 1 | 42 | 0 | 481 | 1 | 1 | |
| B27 | 1 | 43 | 1 | 1590 | 1 | 2 | |
| B28 | 1 | 36 | 1 | 880 | 0 | 1 | |
| B29 | 1 | 40 | 1 | 729 | 1 | 2 | |
| B30 | 1 | 34 | 0 | 561 | 0 | 2 | |
| D01 | 0 | 69 | 1 | N/A | N/A | 0 | |
| D02 | 0 | 52 | 0 | N/A | N/A | 0 | |
| D03 | 0 | 37 | 1 | N/A | N/A | 0 | |
| D04 | 0 | 42 | 0 | N/A | N/A | 0 | |
| D05 | 0 | 50 | 1 | N/A | N/A | 0 | |
| D06 | 0 | 56 | 0 | N/A | N/A | 0 | |
| D07 | 0 | 47 | 1 | N/A | N/A | 0 | |
| D08 | 0 | 72 | 0 | N/A | N/A | 0 | |
| D09 | 0 | 58 | 0 | N/A | N/A | 0 | |
| D10 | 0 | 46 | 1 | N/A | N/A | 0 | |
| D11 | 0 | 41 | 1 | N/A | N/A | 0 | |
| D12 | 0 | 54 | 1 | N/A | N/A | 0 | |
| D13 | 0 | 46 | 0 | N/A | N/A | 0 | |
| D14 | 0 | 42 | 0 | N/A | N/A | 1 | |
| D15 | 0 | 49 | 0 | N/A | N/A | 0 | |
| D16 | 0 | 34 | 0 | N/A | N/A | 0 | |
| D17 | 0 | 70 | 1 | N/A | N/A | 0 | |
| D18 | 0 | 36 | 1 | N/A | N/A | 0 | |
| D19 | 0 | 43 | 1 | N/A | N/A | 0 | |
| D20 | 0 | 37 | 0 | N/A | N/A | 2 | |

**Lyme Borreliosis**: 0=no, 1=yes
**AGE**: age at study enrollment (v1)
**GENDER**: 0=female, 1=male
**RASHSIZE**: EM rash size (cm2)
**DISSEM**: disseminated (multiple) EM rashes, 0=no, 1=yes
**SEROGRP**: CDC serostatus group (0= negative v1/v2, 1= negative v1/converted to positive v2, 2= positive at v1/v2)
**DURAT**: duration of illness prior to study visit (days)

**Table S2.** Sample distributions in two LD cohorts

Demographic and clinical characteristics of the NYMC LD patients are similar to those of the SLICE cohort; the time points represented are somewhat different however, where the 3rd and 4th draws of samples were collected at 1 year and 4-6 years after antibiotic treatment, respectively. There was no 6-month time point as in the discovery cohort.

***SLICE (JHU)***

| Groups | Number of patients | Time points and number of serum samples | | | |
|---|---|---|---|---|---|
| | | Pre-treatment | 4 weeks post treatment | 6 months post treatment | 12 months post treatment |
| LD | 40 | 40 | 40 | 40 | 40 |
| Control | 20 | 20 | | 20 | |

***NYMC***

| Groups | Number of patients | Time points and number of serum samples | | | |
|---|---|---|---|---|---|
| | | Pre-treatment | Convalesce | 1-year post treatment | 4-5 years post treatment |
| LD - With symptom at acute | 30 | 30 | 30 | 30 | 30 |
| Control | 20 | 20 | | 20 | |

**Table S3.** Summarization of SRM methods for 174 monitored proteotypic peptides

Included information of gene symbol, UniProtKB ID, peptide sequence, Q1/Q3 m/z and transition_group_id for 66 target proteins.

| Gene symbol | recommended protein name | UniprotKB ID | Peptide sequence | Q1 | Q3 | prec_z | frg_type | frg_nr | frg_z |
|---|---|---|---|---|---|---|---|---|---|
| A1BG | Alpha-1B-glycoprotein | P04217 | CEGPIPDVTFELLR | 823.4 | 1089.6 | 2 | y | 9 | 1 |
| A1BG | Alpha-1B-glycoprotein | P04217 | CEGPIPDVTFELLR | 823.4 | 1089.6 | 2 | y | 9 | 1 |
| A1BG | Alpha-1B-glycoprotein | P04217 | CEGPIPDVTFELLR | 823.4 | 778.4 | 2 | y | 6 | 1 |
| A1BG | Alpha-1B-glycoprotein | P04217 | CEGPIPDVTFELLR | 823.4 | 545.3 | 2 | y | 9 | 2 |
| A1BG | Alpha-1B-glycoprotein | P04217 | SGLSTGWTQLSK | 632.8 | 1007.5 | 2 | y | 9 | 1 |
| A1BG | Alpha-1B-glycoprotein | P04217 | SGLSTGWTQLSK | 632.8 | 920.5 | 2 | y | 8 | 1 |
| A1BG | Alpha-1B-glycoprotein | P04217 | SGLSTGWTQLSK | 632.8 | 819.4 | 2 | y | 7 | 1 |
| A1BG | Alpha-1B-glycoprotein | P04217 | SGLSTGWTQLSK | 632.8 | 234.1 | 2 | y | 2 | 1 |
| ACTC1 | Actin, alpha cardiac muscle 1 | P68032 | DSYVGDEAQSK | 599.8 | 833.4 | 2 | y | 8 | 1 |
| ACTC1 | Actin, alpha cardiac muscle 1 | P68032 | DSYVGDEAQSK | 599.8 | 734.3 | 2 | y | 7 | 1 |
| ACTC1 | Actin, alpha cardiac muscle 1 | P68032 | DSYVGDEAQSK | 599.8 | 234.1 | 2 | y | 2 | 1 |
| ACTC1 | Actin, alpha cardiac muscle 1 | P68032 | DSYVGDEAQSK | 599.8 | 498.7 | 2 | y | 9 | 2 |
| ACTC1 | Actin, alpha cardiac muscle 1 | P68032 | IIAPPER | 398.2 | 682.4 | 2 | y | 6 | 1 |
| ACTC1 | Actin, alpha cardiac muscle 1 | P68032 | IIAPPER | 398.2 | 569.3 | 2 | y | 5 | 1 |
| ACTC1 | Actin, alpha cardiac muscle 1 | P68032 | IIAPPER | 398.2 | 498.3 | 2 | y | 4 | 1 |
| ACTC1 | Actin, alpha cardiac muscle 1 | P68032 | IIAPPER | 398.2 | 249.6 | 2 | y | 4 | 2 |
| ADPRHL1 | ADP-ribosylarginine] hydrolase-like protein 1 | Q8NDY3 | AIFPDNYDAEER | 720.3 | 1108.5 | 2 | y | 9 | 1 |
| ADPRHL1 | ADP-ribosylarginine] hydrolase-like protein 1 | Q8NDY3 | AIFPDNYDAEER | 720.3 | 896.4 | 2 | y | 7 | 1 |
| ADPRHL1 | ADP-ribosylarginine] hydrolase-like protein 1 | Q8NDY3 | AIFPDNYDAEER | 720.3 | 554.7 | 2 | y | 9 | 2 |
| ADPRHL1 | [Protein ADP-ribosylarginine] hydrolase-like protein 1 | Q8NDY3 | LEDLGAALYR | 560.8 | 878.5 | 2 | y | 8 | 1 |
| ADPRHL1 | [Protein ADP-ribosylarginine] hydrolase-like protein 1 | Q8NDY3 | LEDLGAALYR | 560.8 | 763.4 | 2 | y | 7 | 1 |
| ADPRHL1 | [Protein ADP-ribosylarginine] hydrolase-like protein 1 | Q8NDY3 | LEDLGAALYR | 560.8 | 650.4 | 2 | y | 6 | 1 |
| ADPRHL1 | [Protein ADP-ribosylarginine] hydrolase-like protein 1 | Q8NDY3 | LEDLGAALYR | 560.8 | 593.3 | 2 | y | 5 | 1 |
| AFM | Afamin | P43652 | LPNNVLQEK | 527.8 | 941.5 | 2 | y | 8 | 1 |
| AFM | Afamin | P43652 | LPNNVLQEK | 527.8 | 844.5 | 2 | y | 7 | 1 |
| AFM | Afamin | P43652 | LPNNVLQEK | 527.8 | 730.4 | 2 | y | 6 | 1 |
| AFM | Afamin | P43652 | LPNNVLQEK | 527.8 | 616.4 | 2 | y | 5 | 1 |
| AGXT | Serine--pyruvate aminotransferase, mitochondrial | P21549 | ALNAPPGTSLISFSDK | 809.4 | 628.8 | 2 | y | 12 | 2 |
| AGXT | Serine--pyruvate aminotransferase, mitochondrial | P21549 | ALNAPPGTSLISFSDK | 809.4 | 1256.7 | 2 | y | 12 | 1 |
| AGXT | Serine--pyruvate aminotransferase, mitochondrial | P21549 | ALNAPPGTSLISFSDK | 809.4 | 1159.6 | 2 | y | 11 | 1 |
| AGXT | Serine--pyruvate aminotransferase, mitochondrial | P21549 | ALNAPPGTSLISFSDK | 809.4 | 591.3 | 2 | y | 5 | 1 |
| AGXT | Serine--pyruvate aminotransferase | P21549 | LQALGLQLFVK | 615.4 | 988.6 | 2 | y | 9 | 1 |
| AGXT | Serine--pyruvate aminotransferase | P21549 | LQALGLQLFVK | 615.4 | 917.6 | 2 | y | 8 | 1 |
| AGXT | Serine--pyruvate aminotransferase | P21549 | LQALGLQLFVK | 615.4 | 804.5 | 2 | y | 7 | 1 |
| AGXT | Serine--pyruvate aminotransferase | P21549 | LQALGLQLFVK | 615.4 | 506.3 | 2 | y | 4 | 1 |
| AHSG | Alpha-2-HS-glycoprotein | P02765 | CNLLAEK | 424.2 | 687.4 | 2 | y | 6 | 1 |
| AHSG | Alpha-2-HS-glycoprotein | P02765 | CNLLAEK | 424.2 | 573.4 | 2 | y | 5 | 1 |
| AHSG | Alpha-2-HS-glycoprotein | P02765 | CNLLAEK | 424.2 | 460.3 | 2 | y | 4 | 1 |
| AHSG | Alpha-2-HS-glycoprotein | P02765 | CNLLAEK | 424.2 | 347.2 | 2 | y | 3 | 1 |
| AK5 | Adenylate kinase isoenzyme 5 | Q9Y6K8 | GFLIDGYPR | 519.3 | 833.5 | 2 | y | 7 | 1 |
| AK5 | Adenylate kinase isoenzyme 5 | Q9Y6K8 | GFLIDGYPR | 519.3 | 720.4 | 2 | y | 6 | 1 |
| AK5 | Adenylate kinase isoenzyme 5 | Q9Y6K8 | GFLIDGYPR | 519.3 | 607.3 | 2 | y | 5 | 1 |
| AK5 | Adenylate kinase isoenzyme 5 | Q9Y6K8 | GFLIDGYPR | 519.3 | 492.3 | 2 | y | 4 | 1 |
| AKR1C1 | Aldo-keto reductase family 1 member C1 | Q04828 | HIDSAHLYNNEEQVGLAIR | 727.0 | 1014.6 | 3 | y | 9 | 1 |
| AKR1C1 | Aldo-keto reductase family 1 member C1 | Q04828 | HIDSAHLYNNEEQVGLAIR | 727.0 | 885.5 | 3 | y | 8 | 1 |
| AKR1C1 | Aldo-keto reductase family 1 member C1 | Q04828 | HIDSAHLYNNEEQVGLAIR | 727.0 | 756.5 | 3 | y | 7 | 1 |
| AKR1C1 | Aldo-keto reductase family 1 member C1 | Q04828 | HIDSAHLYNNEEQVGLAIR | 727.0 | 529.3 | 3 | y | 5 | 1 |
| ALDOB | Fructose-bisphosphate aldolase B | P05062 | ALQASALAAWGGK | 622.3 | 931.5 | 2 | y | 10 | 1 |
| ALDOB | Fructose-bisphosphate aldolase B | P05062 | ALQASALAAWGGK | 622.3 | 860.5 | 2 | y | 9 | 1 |
| ALDOB | Fructose-bisphosphate aldolase B | P05062 | ALQASALAAWGGK | 622.3 | 589.3 | 2 | y | 6 | 1 |
| ALDOB | Fructose-bisphosphate aldolase B | P05062 | ALQASALAAWGGK | 622.3 | 447.2 | 2 | y | 4 | 1 |
| ALDOB | Fructose-bisphosphate aldolase B | P05062 | ELSEIAQSIVANGK | 729.9 | 887.5 | 2 | y | 9 | 1 |
| ALDOB | Fructose-bisphosphate aldolase B | P05062 | ELSEIAQSIVANGK | 729.9 | 688.4 | 2 | y | 7 | 1 |

| | | | | | | | | | |
|---|---|---|---|---|---|---|---|---|---|
| ALDOB | Fructose-bisphosphate aldolase B | P05062 | ELSEIAQSIVANGK | 729.9 | 488.3 | 2 | y | 5 | 1 |
| ALDOB | Fructose-bisphosphate aldolase B | P05062 | ELSEIAQSIVANGK | 729.9 | 389.2 | 2 | y | 4 | 1 |
| AMBP | Protein AMBP | P02760 | GECVPGEQEPEPILIPR | 640.7 | 934.6 | 3 | y | 8 | 1 |
| AMBP | Protein AMBP | P02760 | GECVPGEQEPEPILIPR | 640.7 | 708.5 | 3 | y | 6 | 1 |
| AMBP | Protein AMBP | P02760 | GECVPGEQEPEPILIPR | 640.7 | 737.9 | 3 | y | 13 | 2 |
| AMBP | Protein AMBP | P02760 | GECVPGEQEPEPILIPR | 640.7 | 354.7 | 3 | y | 6 | 2 |
| APCS | Serum amyloid P-component | P02743 | IVLGQEQDSYGGK | 697.4 | 1181.5 | 2 | y | 11 | 1 |
| APCS | Serum amyloid P-component | P02743 | IVLGQEQDSYGGK | 697.4 | 1068.5 | 2 | y | 10 | 1 |
| APCS | Serum amyloid P-component | P02743 | IVLGQEQDSYGGK | 697.4 | 883.4 | 2 | y | 8 | 1 |
| APCS | Serum amyloid P-component | P02743 | IVLGQEQDSYGGK | 697.4 | 511.3 | 2 | y | 5 | 1 |
| APOA1 | Apolipoprotein A-I | P02647 | DYVSQFEGSALGK | 700.8 | 1023.5 | 2 | y | 10 | 1 |
| APOA1 | Apolipoprotein A-I | P02647 | DYVSQFEGSALGK | 700.8 | 808.4 | 2 | y | 8 | 1 |
| APOA1 | Apolipoprotein A-I | P02647 | DYVSQFEGSALGK | 700.8 | 661.4 | 2 | y | 7 | 1 |
| APOA1 | Apolipoprotein A-I | P02647 | DYVSQFEGSALGK | 700.8 | 532.3 | 2 | y | 6 | 1 |
| APOA1 | Apolipoprotein A-I | P02647 | LLDNWDSVTSTFSK | 806.9 | 1386.6 | 2 | y | 12 | 1 |
| APOA1 | Apolipoprotein A-I | P02647 | LLDNWDSVTSTFSK | 806.9 | 856.4 | 2 | y | 8 | 1 |
| APOA1 | Apolipoprotein A-I | P02647 | LLDNWDSVTSTFSK | 806.9 | 670.3 | 2 | y | 6 | 1 |
| APOA1 | Apolipoprotein A-I | P02647 | LLDNWDSVTSTFSK | 806.9 | 569.3 | 2 | y | 5 | 1 |
| APOA2 | Apolipoprotein A-II | P02652 | SPELQAEAK | 486.8 | 788.4 | 2 | y | 7 | 1 |
| APOA2 | Apolipoprotein A-II | P02652 | SPELQAEAK | 486.8 | 659.4 | 2 | y | 6 | 1 |
| APOA2 | Apolipoprotein A-II | P02652 | SPELQAEAK | 486.8 | 418.2 | 2 | y | 4 | 1 |
| APOA2 | Apolipoprotein A-II | P02652 | SPELQAEAK | 486.8 | 443.2 | 2 | y | 8 | 2 |
| APOA4 | Apolipoprotein A-IV | P06727 | LGPHAGDVEGHLSFLEK | 452.2 | 465.8 | 4 | y | 8 | 2 |
| APOA4 | Apolipoprotein A-IV | P06727 | LGPHAGDVEGHLSFLEK | 452.2 | 564.9 | 4 | y | 16 | 3 |
| APOA4 | Apolipoprotein A-IV | P06727 | LGPHAGDVEGHLSFLEK | 452.2 | 545.9 | 4 | y | 15 | 3 |
| APOA4 | Apolipoprotein A-IV | P06727 | LGPHAGDVEGHLSFLEK | 452.2 | 513.6 | 4 | y | 14 | 3 |
| APOA4 | Apolipoprotein A-IV | P06727 | SELTQQLNALFQDK | 817.9 | 1076.6 | 2 | y | 9 | 1 |
| APOA4 | Apolipoprotein A-IV | P06727 | SELTQQLNALFQDK | 817.9 | 948.5 | 2 | y | 8 | 1 |
| APOA4 | Apolipoprotein A-IV | P06727 | SELTQQLNALFQDK | 817.9 | 835.4 | 2 | y | 7 | 1 |
| APOA4 | Apolipoprotein A-IV | P06727 | SELTQQLNALFQDK | 817.9 | 537.3 | 2 | y | 4 | 1 |
| APOB | Apolipoprotein B-100 | P04114 | GFEPTLEALFGK | 654.8 | 975.6 | 2 | y | 9 | 1 |
| APOB | Apolipoprotein B-100 | P04114 | GFEPTLEALFGK | 654.8 | 777.5 | 2 | y | 7 | 1 |
| APOB | Apolipoprotein B-100 | P04114 | GFEPTLEALFGK | 654.8 | 664.4 | 2 | y | 6 | 1 |
| APOC2 | Apolipoprotein C-II | P02655 | TYLPAVDEK | 518.3 | 771.4 | 2 | y | 7 | 1 |
| APOC2 | Apolipoprotein C-II | P02655 | TYLPAVDEK | 518.3 | 658.3 | 2 | y | 6 | 1 |
| APOC2 | Apolipoprotein C-II | P02655 | TYLPAVDEK | 518.3 | 386.2 | 2 | y | 7 | 2 |
| APOC2 | Apolipoprotein C-II | P02655 | TYLPAVDEK | 518.3 | 329.7 | 2 | y | 6 | 2 |
| APOC4 | Apolipoprotein C-IV | P55056 | AWFLESK | 440.7 | 623.3 | 2 | y | 5 | 1 |
| APOC4 | Apolipoprotein C-IV | P55056 | AWFLESK | 440.7 | 476.3 | 2 | y | 4 | 1 |
| APOC4 | Apolipoprotein C-IV | P55056 | AWFLESK | 440.7 | 363.2 | 2 | y | 3 | 1 |
| APOC4 | Apolipoprotein C-IV | P55056 | AWFLESK | 440.7 | 234.1 | 2 | y | 2 | 1 |
| APOC4 | Apolipoprotein C-IV | P55056 | ELLETVVNR | 536.8 | 830.5 | 2 | y | 7 | 1 |
| APOC4 | Apolipoprotein C-IV | P55056 | ELLETVVNR | 536.8 | 717.4 | 2 | y | 6 | 1 |
| APOC4 | Apolipoprotein C-IV | P55056 | ELLETVVNR | 536.8 | 588.3 | 2 | y | 5 | 1 |
| APOC4 | Apolipoprotein C-IV | P55056 | ELLETVVNR | 536.8 | 487.3 | 2 | y | 4 | 1 |
| ApoE | Apolipoprotein E | P02649 | CLAVYQAGAR | 554.8 | 764.405 | 2 | y | 7 | 1 |
| ApoE | Apolipoprotein E | P02649 | CLAVYQAGAR | 554.8 | 665.3365 | 2 | y | 6 | 1 |
| ApoE | Apolipoprotein E | P02649 | CLAVYQAGAR | 554.8 | 374.2146 | 2 | y | 4 | 1 |
| ApoE | Apolipoprotein E | P02649 | CLAVYQAGAR | 554.8 | 418.2247 | 2 | y | 8 | 2 |
| APOE | Apolipoprotein E | P02649 | LAVYQAGAR | 474.8 | 665.3 | 2 | y | 6 | 1 |
| APOE | Apolipoprotein E | P02649 | LAVYQAGAR | 474.8 | 502.3 | 2 | y | 5 | 1 |
| APOE | Apolipoprotein E | P02649 | LAVYQAGAR | 474.8 | 374.2 | 2 | y | 4 | 1 |
| APOE | Apolipoprotein E | P02649 | LAVYQAGAR | 474.8 | 333.2 | 2 | y | 6 | 2 |
| APOE | Apolipoprotein E | P02649 | LGADMEDVCGR | 611.8 | 981.4 | 2 | y | 8 | 1 |
| APOE | Apolipoprotein E | P02649 | LGADMEDVCGR | 611.8 | 866.3 | 2 | y | 7 | 1 |
| APOE | Apolipoprotein E | P02649 | LGADMEDVCGR | 611.8 | 735.3 | 2 | y | 6 | 1 |
| APOE | Apolipoprotein E | P02649 | LGADMEDVCGR | 611.8 | 491.2 | 2 | y | 8 | 2 |
| APOE | Apolipoprotein E | P02649 | LGADMEDVR | 503.2 | 764.3 | 2 | y | 6 | 1 |
| APOE | Apolipoprotein E | P02649 | LGADMEDVR | 503.2 | 649.3 | 2 | y | 5 | 1 |
| APOE | Apolipoprotein E | P02649 | LGADMEDVR | 503.2 | 518.3 | 2 | y | 4 | 1 |
| APOE | Apolipoprotein E | P02649 | LGADMEDVR | 503.2 | 382.7 | 2 | y | 6 | 2 |
| APOF | Apolipoprotein F | Q13790 | SGVQQLIQYYQDQK | 849.4 | 1085.5 | 2 | y | 8 | 1 |
| APOF | Apolipoprotein F | Q13790 | SGVQQLIQYYQDQK | 849.4 | 972.4 | 2 | y | 7 | 1 |
| APOF | Apolipoprotein F | Q13790 | SGVQQLIQYYQDQK | 849.4 | 681.3 | 2 | y | 5 | 1 |
| AST1 | Aspartate aminotransferase, cytoplasmic | P17174 | IGADFLAR | 431.7 | 749.4 | 2 | y | 7 | 1 |

| | | | | | | | | | |
|---|---|---|---|---|---|---|---|---|---|
| AST1 | Aspartate aminotransferase, cytoplasmic | P17174 | IGADFLAR | 431.7 | 692.4 | 2 | y | 6 | 1 |
| AST1 | Aspartate aminotransferase, cytoplasmic | P17174 | IGADFLAR | 431.7 | 621.3 | 2 | y | 5 | 1 |
| AST1 | Aspartate aminotransferase, cytoplasmic | P17174 | IGADFLAR | 431.7 | 506.3 | 2 | y | 4 | 1 |
| ASTN1 | Astrotactin-1 | O14525 | SITVSALPFLR | 602.4 | 1003.6 | 2 | y | 9 | 1 |
| ASTN1 | Astrotactin-1 | O14525 | SITVSALPFLR | 602.4 | 902.5 | 2 | y | 8 | 1 |
| ASTN1 | Astrotactin-1 | O14525 | SITVSALPFLR | 602.4 | 803.5 | 2 | y | 7 | 1 |
| ASTN1 | Astrotactin-1 | O14525 | SITVSALPFLR | 602.4 | 532.3 | 2 | y | 4 | 1 |
| BHMT | Betaine--homocysteine S-methyltransferase 1 | Q93088 | AGASIIGVNCHFDPTISLK | 667.3 | 901.0 | 3 | y | 16 | 2 |
| BHMT | Betaine--homocysteine S-methyltransferase 1 | Q93088 | AGASIIGVNCHFDPTISLK | 667.3 | 857.5 | 3 | y | 15 | 2 |
| BHMT | Betaine--homocysteine S-methyltransferase 1 | Q93088 | AGASIIGVNCHFDPTISLK | 667.3 | 800.9 | 3 | y | 14 | 2 |
| BHMT | Betaine--homocysteine S-methyltransferase 1 | Q93088 | AGASIIGVNCHFDPTISLK | 667.3 | 744.4 | 3 | y | 13 | 2 |
| BHMT | Betaine--homocysteine S-methyltransferase 1 | Q93088 | AIAEELAPER | 549.8 | 914.4578 | 2 | y | 8 | 1 |
| BHMT | Betaine--homocysteine S-methyltransferase 1 | Q93088 | AIAEELAPER | 549.8 | 714.3781 | 2 | y | 6 | 1 |
| BHMT | Betaine--homocysteine S-methyltransferase 1 | Q93088 | AIAEELAPER | 549.8 | 585.3355 | 2 | y | 5 | 1 |
| BHMT | Betaine--homocysteine S-methyltransferase 1 | Q93088 | AIAEELAPER | 549.8 | 472.2514 | 2 | y | 4 | 1 |
| BMP10 | Bone morphogenetic protein 10 | O95393 | GVCNYPLAEHLTPTK | 567.3 | 772.4 | 3 | y | 13 | 2 |
| BMP10 | Bone morphogenetic protein 10 | O95393 | GVCNYPLAEHLTPTK | 567.3 | 635.3 | 3 | y | 11 | 2 |
| BMP10 | Bone morphogenetic protein 10 | O95393 | GVCNYPLAEHLTPTK | 567.3 | 553.8 | 3 | y | 10 | 2 |
| BMP10 | Bone morphogenetic protein 10 | O95393 | GVCNYPLAEHLTPTK | 567.3 | 515.3 | 3 | y | 13 | 3 |
| BMP10 | Bone morphogenetic protein 10 | O95393 | TLNLSDIPTQDSAK | 751.9 | 1061.5 | 2 | y | 10 | 1 |
| BMP10 | Bone morphogenetic protein 10 | O95393 | TLNLSDIPTQDSAK | 751.9 | 974.5 | 2 | y | 9 | 1 |
| BMP10 | Bone morphogenetic protein 10 | O95393 | TLNLSDIPTQDSAK | 751.9 | 859.5 | 2 | y | 8 | 1 |
| BMP10 | Bone morphogenetic protein 10 | O95393 | TLNLSDIPTQDSAK | 751.9 | 746.4 | 2 | y | 7 | 1 |
| C4BPB | C4b-binding protein beta chain | P20851 | LIQEAPKPECEK | 481.3 | 662.3 | 3 | y | 5 | 1 |
| C4BPB | C4b-binding protein beta chain | P20851 | LIQEAPKPECEK | 481.3 | 608.3 | 3 | y | 10 | 2 |
| C4BPB | C4b-binding protein beta chain | P20851 | LIQEAPKPECEK | 481.3 | 544.3 | 3 | y | 9 | 2 |
| C4BPB | C4b-binding protein beta chain | P20851 | LIQEAPKPECEK | 481.3 | 444.2 | 3 | y | 7 | 2 |
| C5 | Complement C5 | P01031 | TDAPDLPEENQAR | 728.3 | 1168.6 | 2 | y | 10 | 1 |
| C5 | Complement C5 | P01031 | TDAPDLPEENQAR | 728.3 | 1071.5 | 2 | y | 9 | 1 |
| C5 | Complement C5 | P01031 | TDAPDLPEENQAR | 728.3 | 956.5 | 2 | y | 8 | 1 |
| C5 | Complement C5 | P01031 | TDAPDLPEENQAR | 728.3 | 843.4 | 2 | y | 7 | 1 |
| C6 | Complement component C6 | P13671 | DLHLSDVFLK | 396.2 | 621.4 | 3 | y | 5 | 1 |
| C6 | Complement component C6 | P13671 | DLHLSDVFLK | 396.2 | 506.3 | 3 | y | 4 | 1 |
| C6 | Complement component C6 | P13671 | DLHLSDVFLK | 396.2 | 407.3 | 3 | y | 3 | 1 |
| C6 | Complement component C6 | P13671 | DLHLSDVFLK | 396.2 | 479.8 | 3 | y | 8 | 2 |
| C8A | Complement component C8 alpha chain | P07357 | AIDEDCSQYEPIPGSQK | 968.9 | 1018.5 | 2 | y | 9 | 1 |
| C8A | Complement component C8 alpha chain | P07357 | AIDEDCSQYEPIPGSQK | 968.9 | 855.5 | 2 | y | 8 | 1 |
| C8A | Complement component C8 alpha chain | P07357 | AIDEDCSQYEPIPGSQK | 968.9 | 726.4 | 2 | y | 7 | 1 |
| C8A | Complement component C8 alpha chain | P07357 | AIDEDCSQYEPIPGSQK | 968.9 | 516.3 | 2 | y | 5 | 1 |
| C8B | Complement component C8 beta chain | P07358 | IPGIFELGISSQSDR | 809.9 | 1238.6 | 2 | y | 11 | 1 |
| C8B | Complement component C8 beta chain | P07358 | IPGIFELGISSQSDR | 809.9 | 962.5 | 2 | y | 9 | 1 |
| C8B | Complement component C8 beta chain | P07358 | IPGIFELGISSQSDR | 809.9 | 849.4 | 2 | y | 8 | 1 |
| C8B | Complement component C8 beta chain | P07358 | IPGIFELGISSQSDR | 809.9 | 679.3 | 2 | y | 6 | 1 |
| C8B | Complement component C8 beta chain | P07358 | SGFSFGFK | 438.7 | 789.4 | 2 | y | 7 | 1 |
| C8B | Complement component C8 beta chain | P07358 | SGFSFGFK | 438.7 | 732.4 | 2 | y | 6 | 1 |
| C8B | Complement component C8 beta chain | P07358 | SGFSFGFK | 438.7 | 585.3 | 2 | y | 5 | 1 |
| C8B | Complement component C8 beta chain | P07358 | SGFSFGFK | 438.7 | 498.3 | 2 | y | 4 | 1 |
| C8G | Complement component C8 gamma chain | P07360 | SLPVSDSVLSGFEQR | 810.9 | 1224.6 | 2 | y | 11 | 1 |
| C8G | Complement component C8 gamma chain | P07360 | SLPVSDSVLSGFEQR | 810.9 | 836.4 | 2 | y | 7 | 1 |
| C8G | Complement component C8 gamma chain | P07360 | SLPVSDSVLSGFEQR | 810.9 | 723.3 | 2 | y | 6 | 1 |
| C8G | Complement component C8 gamma chain | P07360 | SLPVSDSVLSGFEQR | 810.9 | 710.9 | 2 | y | 13 | 2 |
| C8G | Complement component C8 gamma chain | P07360 | VQEAHLTEDQIFYFPK | 655.7 | 814.4 | 3 | y | 6 | 1 |
| C8G | Complement component C8 gamma chain | P07360 | VQEAHLTEDQIFYFPK | 655.7 | 701.4 | 3 | y | 5 | 1 |
| C8G | Complement component C8 gamma chain | P07360 | VQEAHLTEDQIFYFPK | 655.7 | 391.2 | 3 | y | 3 | 1 |
| C8G | Complement component C8 gamma chain | P07360 | VQEAHLTEDQIFYFPK | 655.7 | 869.4 | 3 | y | 14 | 2 |
| C9 | Complement component C9 | P02748 | LSPIYNLVPVK | 621.9 | 832.5 | 2 | y | 7 | 1 |
| C9 | Complement component C9 | P02748 | LSPIYNLVPVK | 621.9 | 442.3 | 2 | y | 4 | 1 |
| C9 | Complement component C9 | P02748 | LSPIYNLVPVK | 621.9 | 521.8 | 2 | y | 9 | 2 |
| C9 | Complement component C9 | P02748 | LSPIYNLVPVK | 621.9 | 278.2 | 2 | y | 5 | 2 |
| CA1 | Carbonic anhydrase 1 | P00915 | LYPIANGNNQSPVDIK | 581.6 | 658.4 | 3 | y | 6 | 1 |
| CA1 | Carbonic anhydrase 1 | P00915 | LYPIANGNNQSPVDIK | 581.6 | 571.3 | 3 | y | 5 | 1 |
| CA1 | Carbonic anhydrase 1 | P00915 | LYPIANGNNQSPVDIK | 581.6 | 260.2 | 3 | y | 2 | 1 |
| CA1 | Carbonic anhydrase 1 | P00915 | LYPIANGNNQSPVDIK | 581.6 | 733.9 | 3 | y | 14 | 2 |
| CA1 | Carbonic anhydrase 1 | P00915 | VLDALQAIK | 485.8 | 758.4 | 2 | y | 7 | 1 |
| CA1 | Carbonic anhydrase 1 | P00915 | VLDALQAIK | 485.8 | 459.3 | 2 | y | 4 | 1 |

| | | | | | | | | | |
|---|---|---|---|---|---|---|---|---|---|
| CA1 | Carbonic anhydrase 1 | P00915 | VLDALQAIK | 485.8 | 331.2 | 2 | y | 3 | 1 |
| CA1 | Carbonic anhydrase 1 | P00915 | VLDALQAIK | 485.8 | 260.2 | 2 | y | 2 | 1 |
| CD5L | CD5 antigen-like | O43866 | LVGGDNLCSGR | 574.3 | 935.4 | 2 | y | 9 | 1 |
| CD5L | CD5 antigen-like | O43866 | LVGGDNLCSGR | 574.3 | 706.3 | 2 | y | 6 | 1 |
| CD5L | CD5 antigen-like | O43866 | LVGGDNLCSGR | 574.3 | 479.2 | 2 | y | 4 | 1 |
| CD5L | CD5 antigen-like | O43866 | LVGGDNLCSGR | 574.3 | 319.2 | 2 | y | 3 | 1 |
| CES1 | Carboxylesterase 1 | P23141 | EGYLQIGANTQAAQK | 796.4 | 1129.596 | 2 | y | 11 | 1 |
| CES1 | Carboxylesterase 1 | P23141 | EGYLQIGANTQAAQK | 796.4 | 1001.537 | 2 | y | 10 | 1 |
| CES1 | Carboxylesterase 1 | P23141 | EGYLQIGANTQAAQK | 796.4 | 888.4534 | 2 | y | 9 | 1 |
| CES1 | Carboxylesterase 1 | P23141 | EGYLQIGANTQAAQK | 796.4 | 417.2456 | 2 | y | 4 | 1 |
| CES1 | Carboxylesterase 1 | P23141 | GNWGHLDQVAALR | 479.6 | 529.3 | 3 | y | 5 | 1 |
| CES1 | Carboxylesterase 1 | P23141 | GNWGHLDQVAALR | 479.6 | 430.3 | 3 | y | 4 | 1 |
| CES1 | Carboxylesterase 1 | P23141 | GNWGHLDQVAALR | 479.6 | 633.3 | 3 | y | 11 | 2 |
| CES1 | Carboxylesterase 1 | P23141 | GNWGHLDQVAALR | 479.6 | 511.8 | 3 | y | 9 | 2 |
| CFB | Complement factor B | P00751 | LEDSVTYHCSR | 456.2 | 823.4 | 3 | y | 6 | 1 |
| CFB | Complement factor B | P00751 | LEDSVTYHCSR | 456.2 | 559.2 | 3 | y | 4 | 1 |
| CFB | Complement factor B | P00751 | LEDSVTYHCSR | 456.2 | 562.7 | 3 | y | 9 | 2 |
| CFB | Complement factor B | P00751 | LEDSVTYHCSR | 456.2 | 412.2 | 3 | y | 6 | 2 |
| CFH | Complement factor H | P08603 | RPYFPVAVGK | 378.6 | 570.4 | 3 | y | 6 | 1 |
| CFH | Complement factor H | P08603 | RPYFPVAVGK | 378.6 | 473.3 | 3 | y | 5 | 1 |
| CFH | Complement factor H | P08603 | RPYFPVAVGK | 378.6 | 374.2 | 3 | y | 4 | 1 |
| CFH | Complement factor H | P08603 | RPYFPVAVGK | 378.6 | 285.7 | 3 | y | 6 | 2 |
| CFHR1 | Complement factor H-related protein 1 | B1AKG0 | ITCTEEGWSPTPK | 753.4 | 1291.6 | 2 | y | 11 | 1 |
| CFHR1 | Complement factor H-related protein 1 | B1AKG0 | ITCTEEGWSPTPK | 753.4 | 901.4 | 2 | y | 8 | 1 |
| CFHR1 | Complement factor H-related protein 1 | B1AKG0 | ITCTEEGWSPTPK | 753.4 | 772.4 | 2 | y | 7 | 1 |
| CFHR1 | Complement factor H-related protein 1 | B1AKG0 | ITCTEEGWSPTPK | 753.4 | 529.3 | 2 | y | 5 | 1 |
| CFP | Properdin | P27918 | SISCQEIPGQQSR | 745.4 | 914.5 | 2 | y | 8 | 1 |
| CFP | Properdin | P27918 | SISCQEIPGQQSR | 745.4 | 785.4 | 2 | y | 7 | 1 |
| CFP | Properdin | P27918 | SISCQEIPGQQSR | 745.4 | 672.3 | 2 | y | 6 | 1 |
| CFP | Properdin | P27918 | SISCQEIPGQQSR | 745.4 | 601.8 | 2 | y | 10 | 2 |
| CFP | Properdin | P27918 | TCNHPVPQHGGPFCAGDATR | 545.5 | 590.3 | 4 | y | 6 | 1 |
| CFP | Properdin | P27918 | TCNHPVPQHGGPFCAGDATR | 545.5 | 519.3 | 4 | y | 5 | 1 |
| CFP | Properdin | P27918 | TCNHPVPQHGGPFCAGDATR | 545.5 | 462.2 | 4 | y | 4 | 1 |
| CFP | Properdin | P27918 | TCNHPVPQHGGPFCAGDATR | 545.5 | 347.2 | 4 | y | 3 | 1 |
| CLEC3B | Tetranectin | P05452 | LDTLAQEVALLK | 657.4 | 871.5 | 2 | y | 8 | 1 |
| CLEC3B | Tetranectin | P05452 | LDTLAQEVALLK | 657.4 | 800.5 | 2 | y | 7 | 1 |
| CLEC3B | Tetranectin | P05452 | LDTLAQEVALLK | 657.4 | 672.4 | 2 | y | 6 | 1 |
| CLEC3B | Tetranectin | P05452 | LDTLAQEVALLK | 657.4 | 444.3 | 2 | y | 4 | 1 |
| CLEC3B | Tetranectin | P05452 | TFHEASEDCISR | 484.5 | 650.3 | 3 | y | 5 | 1 |
| CLEC3B | Tetranectin | P05452 | TFHEASEDCISR | 484.5 | 535.3 | 3 | y | 4 | 1 |
| CLEC3B | Tetranectin | P05452 | TFHEASEDCISR | 484.5 | 375.2 | 3 | y | 3 | 1 |
| CLEC3B | Tetranectin | P05452 | TFHEASEDCISR | 484.5 | 262.2 | 3 | y | 2 | 1 |
| CLSTN3 | Calsyntenin-3 | Q9BQT9 | VNDVNEFAPVFVER | 817.9 | 1207.6 | 2 | y | 10 | 1 |
| CLSTN3 | Calsyntenin-3 | Q9BQT9 | VNDVNEFAPVFVER | 817.9 | 1093.6 | 2 | y | 9 | 1 |
| CLSTN3 | Calsyntenin-3 | Q9BQT9 | VNDVNEFAPVFVER | 817.9 | 964.5 | 2 | y | 8 | 1 |
| CLSTN3 | Calsyntenin-3 | Q9BQT9 | VNDVNEFAPVFVER | 817.9 | 746.4 | 2 | y | 6 | 1 |
| CLU | Clusterin | P10909 | LFDSDPITVTVPVEVSR | 937.5 | 1296.8 | 2 | y | 12 | 1 |
| CLU | Clusterin | P10909 | LFDSDPITVTVPVEVSR | 937.5 | 1086.6 | 2 | y | 10 | 1 |
| CLU | Clusterin | P10909 | LFDSDPITVTVPVEVSR | 937.5 | 886.5 | 2 | y | 8 | 1 |
| CLU | Clusterin | P10909 | LFDSDPITVTVPVEVSR | 937.5 | 686.4 | 2 | y | 6 | 1 |
| CNDP1 | Beta-Ala-His dipeptidase | Q96KN2 | ALEQDLPVNIK | 620.4 | 798.5 | 2 | y | 7 | 1 |
| CNDP1 | Beta-Ala-His dipeptidase | Q96KN2 | ALEQDLPVNIK | 620.4 | 683.4 | 2 | y | 6 | 1 |
| CNDP1 | Beta-Ala-His dipeptidase | Q96KN2 | ALEQDLPVNIK | 620.4 | 570.4 | 2 | y | 5 | 1 |
| CNDP1 | Beta-Ala-His dipeptidase | Q96KN2 | ALEQDLPVNIK | 620.4 | 374.2 | 2 | y | 3 | 1 |
| CNDP1 | Beta-Ala-His dipeptidase | Q96KN2 | EWVAIESDSVQPVPR | 571.3 | 596.4 | 3 | y | 5 | 1 |
| CNDP1 | Beta-Ala-His dipeptidase | Q96KN2 | EWVAIESDSVQPVPR | 571.3 | 468.3 | 3 | y | 4 | 1 |
| CNDP1 | Beta-Ala-His dipeptidase | Q96KN2 | EWVAIESDSVQPVPR | 571.3 | 272.2 | 3 | y | 2 | 1 |
| CNDP1 | Beta-Ala-His dipeptidase | Q96KN2 | EWVAIESDSVQPVPR | 571.3 | 234.7 | 3 | y | 4 | 2 |
| CPB2 | Carboxypeptidase B2 | Q96IY4 | SFYANNHCIGTDLNR | 594.6 | 675.3 | 3 | y | 6 | 1 |
| CPB2 | Carboxypeptidase B2 | Q96IY4 | SFYANNHCIGTDLNR | 594.6 | 774.4 | 3 | y | 13 | 2 |
| CPB2 | Carboxypeptidase B2 | Q96IY4 | SFYANNHCIGTDLNR | 594.6 | 692.8 | 3 | y | 12 | 2 |
| CPB2 | Carboxypeptidase B2 | Q96IY4 | SFYANNHCIGTDLNR | 594.6 | 657.3 | 3 | y | 11 | 2 |
| CPN1 | Carboxypeptidase N catalytic chain | P15169 | IVQLIQDTR | 543.3 | 873.5 | 2 | y | 7 | 1 |
| CPN1 | Carboxypeptidase N catalytic chain | P15169 | IVQLIQDTR | 543.3 | 745.4 | 2 | y | 6 | 1 |
| CPN1 | Carboxypeptidase N catalytic chain | P15169 | IVQLIQDTR | 543.3 | 632.3 | 2 | y | 5 | 1 |

| | | | | | | | | | |
|---|---|---|---|---|---|---|---|---|---|
| CPN1 | Carboxypeptidase N catalytic chain | P15169 | IVQLIQDTR | 543.3 | 519.3 | 2 | y | 4 | 1 |
| CPN2 | Carboxypeptidase N subunit 2 | P22792 | LLNIQTYCAGPAYLK | 863.0 | 1271.6 | 2 | y | 11 | 1 |
| CPN2 | Carboxypeptidase N subunit 2 | P22792 | LLNIQTYCAGPAYLK | 863.0 | 1143.6 | 2 | y | 10 | 1 |
| CPN2 | Carboxypeptidase N subunit 2 | P22792 | LLNIQTYCAGPAYLK | 863.0 | 879.4 | 2 | y | 8 | 1 |
| CPN2 | Carboxypeptidase N subunit 2 | P22792 | LLNIQTYCAGPAYLK | 863.0 | 648.4 | 2 | y | 6 | 1 |
| CRP | C-reactive protein | P02741 | ESDTSYVSLK | 564.8 | 797.4 | 2 | y | 7 | 1 |
| CRP | C-reactive protein | P02741 | ESDTSYVSLK | 564.8 | 696.4 | 2 | y | 6 | 1 |
| CRP | C-reactive protein | P02741 | ESDTSYVSLK | 564.8 | 609.4 | 2 | y | 5 | 1 |
| CRP | C-reactive protein | P02741 | ESDTSYVSLK | 564.8 | 446.3 | 2 | y | 4 | 1 |
| CRP | C-reactive protein | P02741 | GYSIFSYATK | 568.8 | 916.5 | 2 | y | 8 | 1 |
| CRP | C-reactive protein | P02741 | GYSIFSYATK | 568.8 | 829.4 | 2 | y | 7 | 1 |
| CRP | C-reactive protein | P02741 | GYSIFSYATK | 568.8 | 716.4 | 2 | y | 6 | 1 |
| CST6 | Cystatin-M | Q15828 | AQSQLVAGIK | 507.8 | 815.5 | 2 | y | 8 | 1 |
| CST6 | Cystatin-M | Q15828 | AQSQLVAGIK | 507.8 | 728.5 | 2 | y | 7 | 1 |
| CST6 | Cystatin-M | Q15828 | AQSQLVAGIK | 507.8 | 388.3 | 2 | y | 4 | 1 |
| CTSS | Cathepsin S | P25774 | GIDSDASYPYK | 608.3 | 1045.4 | 2 | y | 9 | 1 |
| CTSS | Cathepsin S | P25774 | GIDSDASYPYK | 608.3 | 930.4 | 2 | y | 8 | 1 |
| CTSS | Cathepsin S | P25774 | GIDSDASYPYK | 608.3 | 843.4 | 2 | y | 7 | 1 |
| CTSS | Cathepsin S | P25774 | GIDSDASYPYK | 608.3 | 407.2 | 2 | y | 3 | 1 |
| DBH | Dopamine beta-hydroxylase | P09172 | AFYYPEEAGLAFGGPGSSR | 659.3 | 764.4 | 3 | y | 8 | 1 |
| DBH | Dopamine beta-hydroxylase | P09172 | AFYYPEEAGLAFGGPGSSR | 659.3 | 617.3 | 3 | y | 7 | 1 |
| DBH | Dopamine beta-hydroxylase | P09172 | AFYYPEEAGLAFGGPGSSR | 659.3 | 560.3 | 3 | y | 6 | 1 |
| DBH | Dopamine beta-hydroxylase | P09172 | AFYYPEEAGLAFGGPGSSR | 659.3 | 252.1 | 3 | y | 5 | 2 |
| DBH | Dopamine beta-hydroxylase | P09172 | FQGEWNLQPLPK | 728.9 | 1181.6 | 2 | y | 10 | 1 |
| DBH | Dopamine beta-hydroxylase | P09172 | FQGEWNLQPLPK | 728.9 | 995.6 | 2 | y | 8 | 1 |
| DBH | Dopamine beta-hydroxylase | P09172 | FQGEWNLQPLPK | 728.9 | 582.4 | 2 | y | 5 | 1 |
| DBH | Dopamine beta-hydroxylase | P09172 | FQGEWNLQPLPK | 728.9 | 454.3 | 2 | y | 4 | 1 |
| EFEMP1 | EGF-containing fibulin-like extracellular matrix protein 1 | Q12805 | GSFACQCPPGYQK | 750.3 | 1137.5 | 2 | y | 9 | 1 |
| EFEMP1 | EGF-containing fibulin-like extracellular matrix protein 1 | Q12805 | GSFACQCPPGYQK | 750.3 | 849.4 | 2 | y | 7 | 1 |
| EFEMP1 | EGF-containing fibulin-like extracellular matrix protein 1 | Q12805 | GSFACQCPPGYQK | 750.3 | 689.4 | 2 | y | 6 | 1 |
| EFEMP1 | EGF-containing fibulin-like extracellular matrix protein 1 | Q12805 | GSFACQCPPGYQK | 750.3 | 592.3 | 2 | y | 5 | 1 |
| EFEMP1 | EGF-containing fibulin-like extracellular matrix protein 1 | Q12805 | LNCEDIDECR | 662.3 | 807.3 | 2 | y | 6 | 1 |
| EFEMP1 | EGF-containing fibulin-like extracellular matrix protein 1 | Q12805 | LNCEDIDECR | 662.3 | 692.3 | 2 | y | 5 | 1 |
| EFEMP1 | EGF-containing fibulin-like extracellular matrix protein 1 | Q12805 | LNCEDIDECR | 662.3 | 579.2 | 2 | y | 4 | 1 |
| EFEMP1 | EGF-containing fibulin-like extracellular matrix protein 1 | Q12805 | LNCEDIDECR | 662.3 | 464.2 | 2 | y | 3 | 1 |
| ENO1 | Enolase 1 | P00924 | AVDDFLISLDGTANK | 789.9 | 1178.6 | 2 | y | 11 | 1 |
| ENO1 | Enolase 1 | P00924 | AVDDFLISLDGTANK | 789.9 | 1031.6 | 2 | y | 10 | 1 |
| ENO1 | Enolase 1 | P00924 | AVDDFLISLDGTANK | 789.9 | 918.5 | 2 | y | 9 | 1 |
| ENO1 | Enolase 1 | P00924 | AVDDFLISLDGTANK | 789.9 | 805.4 | 2 | y | 8 | 1 |
| ENO1 | Alpha-enolase | P06733 | GNPTVEVELTTEK | 708.9 | 1047.6 | 2 | y | 9 | 1 |
| ENO1 | Alpha-enolase | P06733 | GNPTVEVELTTEK | 708.9 | 948.5 | 2 | y | 8 | 1 |
| ENO1 | Alpha-enolase | P06733 | GNPTVEVELTTEK | 708.9 | 819.4 | 2 | y | 7 | 1 |
| ENO1 | Alpha-enolase | P06733 | GNPTVEVELTTEK | 708.9 | 720.4 | 2 | y | 6 | 1 |
| F10 | Coagulation factor X | P00742 | ETYDFDIAVLR | 671.3 | 833.5 | 2 | y | 7 | 1 |
| F10 | Coagulation factor X | P00742 | ETYDFDIAVLR | 671.3 | 686.4 | 2 | y | 6 | 1 |
| F10 | Coagulation factor X | P00742 | ETYDFDIAVLR | 671.3 | 571.4 | 2 | y | 5 | 1 |
| F10 | Coagulation factor X | P00742 | ETYDFDIAVLR | 671.3 | 458.3 | 2 | y | 4 | 1 |
| F10 | Coagulation factor X | P00742 | NCELFTR | 470.2 | 665.4 | 2 | y | 5 | 1 |
| F10 | Coagulation factor X | P00742 | NCELFTR | 470.2 | 536.3 | 2 | y | 4 | 1 |
| F10 | Coagulation factor X | P00742 | NCELFTR | 470.2 | 423.2 | 2 | y | 3 | 1 |
| F10 | Coagulation factor X | P00742 | NCELFTR | 470.2 | 276.2 | 2 | y | 2 | 1 |
| F12 | Coagulation factor XII | P00748 | VVGGLVALR | 442.3 | 784.5 | 2 | y | 8 | 1 |
| F12 | Coagulation factor XII | P00748 | VVGGLVALR | 442.3 | 685.4 | 2 | y | 7 | 1 |
| F12 | Coagulation factor XII | P00748 | VVGGLVALR | 442.3 | 628.4 | 2 | y | 6 | 1 |
| F12 | Coagulation factor XII | P00748 | VVGGLVALR | 442.3 | 458.3 | 2 | y | 4 | 1 |
| F9 | Coagulation factor IX | P00740 | SALVLQYLR | 531.8 | 904.6 | 2 | y | 7 | 1 |
| F9 | Coagulation factor IX | P00740 | SALVLQYLR | 531.8 | 791.5 | 2 | y | 6 | 1 |
| F9 | Coagulation factor IX | P00740 | SALVLQYLR | 531.8 | 692.4 | 2 | y | 5 | 1 |
| F9 | Coagulation factor IX | P00740 | SALVLQYLR | 531.8 | 579.3 | 2 | y | 4 | 1 |
| F9 | Coagulation factor IX | P00740 | SCEPAVPFPCGR | 688.8 | 1000.5 | 2 | y | 9 | 1 |
| F9 | Coagulation factor IX | P00740 | SCEPAVPFPCGR | 688.8 | 832.4 | | y | 7 | 1 |
| F9 | Coagulation factor IX | P00740 | SCEPAVPFPCGR | 688.8 | 733.3 | | y | 6 | 1 |
| F9 | Coagulation factor IX | P00740 | SCEPAVPFPCGR | 688.8 | 489.2 | | y | 4 | 1 |
| FBP1 | Fructose-1,6-bisphosphatase 1 | P09467 | APVILGSPDDVLEFLK | 857.0 | 1162.599 | 2 | y | 10 | 1 |
| FBP1 | Fructose-1,6-bisphosphatase 1 | P09467 | APVILGSPDDVLEFLK | 857.0 | 1075.567 | 2 | y | 9 | 1 |

| | | | | | | | | | |
|---|---|---|---|---|---|---|---|---|---|
| FBP1 | Fructose-1,6-bisphosphatase 1 | P09467 | APVILGSPDDVLEFLK | 857.0 | 536.3079 | 2 | y | 4 | 1 |
| FBP1 | Fructose-1,6-bisphosphatase 1 | P09467 | APVILGSPDDVLEFLK | 857.0 | 268.6576 | 3 | y | 4 | 2 |
| FBP1 | Fructose-1,6-bisphosphatase 1 | P09467 | EAVLDVIPTDIHQR | 536.0 | 653.8593 | 3 | y | 11 | 1 |
| FBP1 | Fructose-1,6-bisphosphatase 1 | P09467 | EAVLDVIPTDIHQR | 536.0 | 597.3173 | 3 | y | 10 | 2 |
| FBP1 | Fructose-1,6-bisphosphatase 1 | P09467 | EAVLDVIPTDIHQR | 536.0 | 490.2696 | 3 | y | 8 | 2 |
| FBP1 | Fructose-1,6-bisphosphatase 1 | P09467 | EAVLDVIPTDIHQR | 536.0 | 433.7276 | 3 | y | 7 | 2 |
| GC | Vitamin D-binding protein | P02774 | ELSSFIDK | 469.7 | 809.4 | 2 | y | 7 | 1 |
| GC | Vitamin D-binding protein | P02774 | ELSSFIDK | 469.7 | 696.4 | 2 | y | 6 | 1 |
| GC | Vitamin D-binding protein | P02774 | ELSSFIDK | 469.7 | 609.3 | 2 | y | 5 | 1 |
| GC | Vitamin D-binding protein | P02774 | ELSSFIDK | 469.7 | 522.3 | 2 | y | 4 | 1 |
| GC | Vitamin D-binding protein | P02774 | HLSLLTTLSNR | 418.9 | 691.4 | 3 | y | 6 | 1 |
| GC | Vitamin D-binding protein | P02774 | HLSLLTTLSNR | 418.9 | 590.3 | 3 | y | 5 | 1 |
| GC | Vitamin D-binding protein | P02774 | HLSLLTTLSNR | 418.9 | 489.3 | 3 | y | 4 | 1 |
| GC | Vitamin D-binding protein | P02774 | HLSLLTTLSNR | 418.9 | 295.7 | 3 | y | 5 | 2 |
| GLUD2 | Glutamate dehydrogenase 2, mitochondrial | P49448 | LQHGSILGFPK | 399.6 | 561.3 | 3 | y | 5 | 1 |
| GLUD2 | Glutamate dehydrogenase 2, mitochondrial | P49448 | LQHGSILGFPK | 399.6 | 448.3 | 3 | y | 4 | 1 |
| GLUD2 | Glutamate dehydrogenase 2, mitochondrial | P49448 | LQHGSILGFPK | 399.6 | 281.2 | 3 | y | 5 | 2 |
| GLUD2 | Glutamate dehydrogenase 2, mitochondrial | P49448 | LQHGSILGFPK | 399.6 | 224.6 | 3 | y | 4 | 2 |
| GOT1 | Aspartate aminotransferase, cytoplasmic | P17174 | ITWSNPPAQGAR | 649.3 | 897.5 | 2 | y | 9 | 1 |
| GOT1 | Aspartate aminotransferase, cytoplasmic | P17174 | ITWSNPPAQGAR | 649.3 | 810.4 | 2 | y | 8 | 1 |
| GOT1 | Aspartate aminotransferase, cytoplasmic | P17174 | ITWSNPPAQGAR | 649.3 | 696.4 | 2 | y | 7 | 1 |
| GOT1 | Aspartate aminotransferase, cytoplasmic | P17174 | IVASTLSNPELFEEWTGNVK | 745.4 | 833.4 | 3 | y | 7 | 1 |
| GOT1 | Aspartate aminotransferase, cytoplasmic | P17174 | IVASTLSNPELFEEWTGNVK | 745.4 | 704.4 | 3 | y | 6 | 1 |
| GOT1 | Aspartate aminotransferase, cytoplasmic | P17174 | IVASTLSNPELFEEWTGNVK | 745.4 | 518.3 | 3 | y | 5 | 1 |
| GOT1 | Aspartate aminotransferase, cytoplasmic | P17174 | IVASTLSNPELFEEWTGNVK | 745.4 | 417.2 | 3 | y | 7 | 2 |
| GOT1 | Aspartate aminotransferase, cytoplasmic | P17174 | VGGVQSLGGTGALR | 636.4 | 959.5 | 2 | y | 10 | 1 |
| GOT1 | Aspartate aminotransferase, cytoplasmic | P17174 | VGGVQSLGGTGALR | 636.4 | 831.5 | 2 | y | 9 | 1 |
| GOT1 | Aspartate aminotransferase, cytoplasmic | P17174 | VGGVQSLGGTGALR | 636.4 | 744.4 | 2 | y | 8 | 1 |
| GOT1 | Aspartate aminotransferase, cytoplasmic | P17174 | VGGVQSLGGTGALR | 636.4 | 586.8 | 2 | y | 13 | 2 |
| GOT2 | Aspartate aminotransferase, mitochondrial | P00505 | ISVAGVTSSNVGYLAHAIHQVTK | 588.8 | 913.5 | 4 | y | 17 | 2 |
| GOT2 | Aspartate aminotransferase, mitochondrial | P00505 | ISVAGVTSSNVGYLAHAIHQVTK | 588.8 | 863.0 | 4 | y | 16 | 2 |
| GOT2 | Aspartate aminotransferase, mitochondrial | P00505 | ISVAGVTSSNVGYLAHAIHQVTK | 588.8 | 669.4 | 4 | y | 12 | 2 |
| GPT | Alanine aminotransferase 1 | P24298 | ALCVINPGNPTGQVQTR | 550.8 | 916.5 | 2 | y | 7 | 1 |
| GPT | Alanine aminotransferase 1 | P24298 | ALCVINPGNPTGQVQTR | 550.8 | 787.4 | 2 | y | 6 | 1 |
| GPT | Alanine aminotransferase 1 | P24298 | ALCVINPGNPTGQVQTR | 550.8 | 674.3 | 2 | y | 5 | 1 |
| GPT | Alanine aminotransferase 1 | P24298 | AWALDVAELHR | 427.6 | 839.4 | 3 | y | 7 | 1 |
| GPT | Alanine aminotransferase 1 | P24298 | AWALDVAELHR | 427.6 | 512.3 | 3 | y | 9 | 2 |
| GPT | Alanine aminotransferase 1 | P24298 | AWALDVAELHR | 427.6 | 476.8 | 3 | y | 8 | 2 |
| GPT | Alanine aminotransferase 1 | P24298 | AWALDVAELHR | 427.6 | 420.2 | 3 | y | 7 | 2 |
| GPT | Alanine aminotransferase 1 | P24298 | LLVAGEGHTR | 351.5 | 727.3 | 3 | y | 7 | 1 |
| GPT | Alanine aminotransferase 1 | P24298 | LLVAGEGHTR | 351.5 | 656.3 | 3 | y | 6 | 1 |
| GPT | Alanine aminotransferase 1 | P24298 | LLVAGEGHTR | 351.5 | 364.2 | 3 | y | 7 | 2 |
| GPT | Alanine aminotransferase 1 | P24298 | LLVAGEGHTR | 351.5 | 328.7 | 3 | y | 6 | 2 |
| GPT | Alanine aminotransferase 1 | P24298 | ALELEQELR | 550.8 | 916.5 | 2 | y | 7 | 1 |
| GPT | Alanine aminotransferase 1 | P24298 | ALELEQELR | 550.8 | 787.4 | 2 | y | 6 | 1 |
| GPT | Alanine aminotransferase 1 | P24298 | ALELEQELR | 550.8 | 674.3 | 2 | y | 5 | 1 |
| GPT2 | Alanine aminotransferase 2 | Q8TD30 | DFHINFLEK | 388.2 | 536.3 | 3 | y | 4 | 1 |
| GPT2 | Alanine aminotransferase 2 | Q8TD30 | DFHINFLEK | 388.2 | 450.8 | 3 | y | 7 | 2 |
| GPT2 | Alanine aminotransferase 2 | Q8TD30 | DFHINFLEK | 388.2 | 268.7 | 3 | y | 4 | 2 |
| GSTO1 | Glutathione S-transferase omega-1 | P78417 | GSAPPGPVPEGSIR | 660.8 | 1008.5 | 2 | y | 10 | 1 |
| GSTO1 | Glutathione S-transferase omega-1 | P78417 | GSAPPGPVPEGSIR | 660.8 | 658.4 | 2 | y | 6 | 1 |
| GSTO1 | Glutathione S-transferase omega-1 | P78417 | GSAPPGPVPEGSIR | 660.8 | 553.3 | 2 | y | 11 | 2 |
| GSTO1 | Glutathione S-transferase omega-1 | P78417 | GSAPPGPVPEGSIR | 660.8 | 504.8 | 2 | y | 10 | 2 |
| HABP2 | Hyaluronan-binding protein 2 | Q14520 | LIANTLCNSR | 581.3 | 935.4363 | 2 | y | 8 | 1 |
| HABP2 | Hyaluronan-binding protein 2 | Q14520 | LIANTLCNSR | 581.3 | 864.3992 | 2 | y | 7 | 1 |
| HABP2 | Hyaluronan-binding protein 2 | Q14520 | LIANTLCNSR | 581.3 | 649.3086 | 2 | y | 5 | 1 |
| HABP2 | Hyaluronan-binding protein 2 | Q14520 | LIANTLCNSR | 581.3 | 536.2246 | 2 | y | 4 | 1 |
| HGFAC | Hepatocyte growth factor activator | Q04756 | VANYVDWINDR | 682.8 | 917.4 | 2 | y | 7 | 1 |
| HGFAC | Hepatocyte growth factor activator | Q04756 | VANYVDWINDR | 682.8 | 818.4 | 2 | y | 6 | 1 |
| HGFAC | Hepatocyte growth factor activator | Q04756 | VANYVDWINDR | 682.8 | 703.4 | 2 | y | 5 | 1 |
| HGFAC | Hepatocyte growth factor activator | Q04756 | VANYVDWINDR | 682.8 | 517.3 | 2 | y | 4 | 1 |
| HGFAC | Hepatocyte growth factor activator | Q04756 | YEYLEGGDR | 551.2 | 938.4 | 2 | y | 8 | 1 |
| HGFAC | Hepatocyte growth factor activator | Q04756 | YEYLEGGDR | 551.2 | 809.4 | 2 | y | 7 | 1 |
| HGFAC | Hepatocyte growth factor activator | Q04756 | YEYLEGGDR | 551.2 | 646.3 | 2 | y | 6 | 1 |
| HGFAC | Hepatocyte growth factor activator | Q04756 | YEYLEGGDR | 551.2 | 404.2 | 2 | y | 4 | 1 |

| | | | | | | | | | |
|---|---|---|---|---|---|---|---|---|---|
| HP | Haptoglobin | P00738 | VTSIQDWVQK | 602.3 | 1003.5 | 2 | y | 8 | 1 |
| HP | Haptoglobin | P00738 | VTSIQDWVQK | 602.3 | 803.4 | 2 | y | 6 | 1 |
| HP | Haptoglobin | P00738 | VTSIQDWVQK | 602.3 | 560.3 | 2 | y | 4 | 1 |
| HPCAL4 | Hippocalcin-like protein 4 | Q9UM19 | DCPSGILNLEEFQQLYIK | 723.0 | 1068.6 | 3 | y | 8 | 1 |
| HPCAL4 | Hippocalcin-like protein 4 | Q9UM19 | DCPSGILNLEEFQQLYIK | 723.0 | 423.3 | 3 | y | 3 | 1 |
| HPCAL4 | Hippocalcin-like protein 4 | Q9UM19 | DCPSGILNLEEFQQLYIK | 723.0 | 260.2 | 3 | y | 2 | 1 |
| HPCAL4 | Hippocalcin-like protein 4 | Q9UM19 | DCPSGILNLEEFQQLYIK | 723.0 | 946.5 | 3 | y | 16 | 2 |
| HPX | Hemopexin | P02790 | LWWLDLK | 487.3 | 860.5 | 2 | y | 6 | 1 |
| HPX | Hemopexin | P02790 | LWWLDLK | 487.3 | 674.4 | 2 | y | 5 | 1 |
| HPX | Hemopexin | P02790 | LWWLDLK | 487.3 | 488.3 | 2 | y | 4 | 1 |
| HPX | Hemopexin | P02790 | LWWLDLK | 487.3 | 375.2 | 2 | y | 3 | 1 |
| HPX | Hemopexin | P02790 | SGAQATWTELPWPHEK | 613.3 | 793.4 | 3 | y | 6 | 1 |
| HPX | Hemopexin | P02790 | SGAQATWTELPWPHEK | 613.3 | 510.3 | 3 | y | 4 | 1 |
| HPX | Hemopexin | P02790 | SGAQATWTELPWPHEK | 613.3 | 811.9 | 3 | y | 13 | 2 |
| HPX | Hemopexin | P02790 | SGAQATWTELPWPHEK | 613.3 | 397.2 | 3 | y | 6 | 2 |
| IFNA6 | Interferon alpha-6 | P05013 | ISLFSCLK | 484.3 | 854.4 | 2 | y | 7 | 1 |
| IFNA6 | Interferon alpha-6 | P05013 | ISLFSCLK | 484.3 | 767.4 | 2 | y | 6 | 1 |
| IFNA6 | Interferon alpha-6 | P05013 | ISLFSCLK | 484.3 | 654.3 | 2 | y | 5 | 1 |
| IGFALS | Insulin-like growth factor-binding protein complex acid labile subunit | P35858 | DFALQNPSAVPR | 657.8 | 981.5 | 2 | y | 9 | 1 |
| IGFALS | Insulin-like growth factor-binding protein complex acid labile subunit | P35858 | DFALQNPSAVPR | 657.8 | 868.5 | 2 | y | 8 | 1 |
| IGFALS | Insulin-like growth factor-binding protein complex acid labile subunit | P35858 | DFALQNPSAVPR | 657.8 | 740.4 | 2 | y | 7 | 1 |
| IGFALS | Insulin-like growth factor-binding protein complex acid labile subunit | P35858 | DFALQNPSAVPR | 657.8 | 626.4 | 2 | y | 6 | 1 |
| IL1RAP | Interleukin-1 receptor accessory protein | Q9NPH3 | VAFPLEVVQK | 565.3 | 812.5 | 2 | y | 7 | 1 |
| IL1RAP | Interleukin-1 receptor accessory protein | Q9NPH3 | VAFPLEVVQK | 565.3 | 602.4 | 2 | y | 5 | 1 |
| IL1RAP | Interleukin-1 receptor accessory protein | Q9NPH3 | VAFPLEVVQK | 565.3 | 374.2 | 2 | y | 3 | 1 |
| IL1RAP | Interleukin-1 receptor accessory protein | Q9NPH3 | VAFPLEVVQK | 565.3 | 406.7 | 2 | y | 7 | 2 |
| INHBC | Inhibin beta C chain | P55103 | LDFHFSSDR | 375.2 | 611.3 | 3 | y | 5 | 1 |
| INHBC | Inhibin beta C chain | P55103 | LDFHFSSDR | 375.2 | 464.2 | 3 | y | 4 | 1 |
| INHBC | Inhibin beta C chain | P55103 | LDFHFSSDR | 375.2 | 505.7 | 3 | y | 8 | 2 |
| INHBC | Inhibin beta C chain | P55103 | LDFHFSSDR | 375.2 | 448.2 | 3 | y | 7 | 2 |
| ITIH2 | Inter-alpha-trypsin inhibitor heavy chain H2 | P19823 | VQFELHYQEVK | 473.9 | 666.3 | 3 | y | 5 | 1 |
| ITIH2 | Inter-alpha-trypsin inhibitor heavy chain H2 | P19823 | VQFELHYQEVK | 473.9 | 660.8 | 3 | y | 10 | 2 |
| ITIH2 | Inter-alpha-trypsin inhibitor heavy chain H2 | P19823 | VQFELHYQEVK | 473.9 | 596.8 | 3 | y | 9 | 2 |
| ITIH2 | Inter-alpha-trypsin inhibitor heavy chain H2 | P19823 | VQFELHYQEVK | 473.9 | 402.2 | 3 | y | 6 | 2 |
| ITIH4 | Inter-alpha-trypsin inhibitor heavy chain H4 | Q14624 | ILDDLSPR | 464.8 | 815.4 | 2 | y | 7 | 1 |
| ITIH4 | Inter-alpha-trypsin inhibitor heavy chain H4 | Q14624 | ILDDLSPR | 464.8 | 702.3 | 2 | y | 6 | 1 |
| ITIH4 | Inter-alpha-trypsin inhibitor heavy chain H4 | Q14624 | ILDDLSPR | 464.8 | 587.3 | 2 | y | 5 | 1 |
| ITIH4 | Inter-alpha-trypsin inhibitor heavy chain H4 | Q14624 | ILDDLSPR | 464.8 | 472.3 | 2 | y | 4 | 1 |
| ITIH4 | Inter-alpha-trypsin inhibitor heavy chain H4 | Q14624 | LGVYELLLK | 524.3 | 934.6 | 2 | y | 8 | 1 |
| ITIH4 | Inter-alpha-trypsin inhibitor heavy chain H4 | Q14624 | LGVYELLLK | 524.3 | 778.5 | 2 | y | 6 | 1 |
| ITIH4 | Inter-alpha-trypsin inhibitor heavy chain H4 | Q14624 | LGVYELLLK | 524.3 | 615.4 | 2 | y | 5 | 1 |
| ITIH4 | Inter-alpha-trypsin inhibitor heavy chain H4 | Q14624 | LGVYELLLK | 524.3 | 486.4 | 2 | y | 4 | 1 |
| JSRP1 | Junctional sarcoplasmic reticulum protein 1 | Q96MG2 | DAVPGEAALQAR | 599.3 | 815.4 | 2 | y | 8 | 1 |
| JSRP1 | Junctional sarcoplasmic reticulum protein 1 | Q96MG2 | DAVPGEAALQAR | 599.3 | 629.4 | 2 | y | 6 | 1 |
| JSRP1 | Junctional sarcoplasmic reticulum protein 1 | Q96MG2 | DAVPGEAALQAR | 599.3 | 558.3 | 2 | y | 5 | 1 |
| JSRP1 | Junctional sarcoplasmic reticulum protein 1 | Q96MG2 | DAVPGEAALQAR | 599.3 | 456.7 | 2 | y | 9 | 2 |
| KIT | Mast/stem cell growth factor receptor Kit | P10721 | LVVQSSIDSSAFK | 690.9 | 1168.6 | 2 | y | 11 | 1 |
| KIT | Mast/stem cell growth factor receptor Kit | P10721 | LVVQSSIDSSAFK | 690.9 | 1069.5 | 2 | y | 10 | 1 |
| KIT | Mast/stem cell growth factor receptor Kit | P10721 | LVVQSSIDSSAFK | 690.9 | 941.5 | 2 | y | 9 | 1 |
| KIT | Mast/stem cell growth factor receptor Kit | P10721 | LVVQSSIDSSAFK | 690.9 | 584.8 | 2 | y | 11 | 2 |
| KNG1 | Kininogen-1 | P01042 | DIPTNSPELEETLTHTITK | 713.7 | 956.0 | 3 | y | 17 | 2 |
| KNG1 | Kininogen-1 | P01042 | DIPTNSPELEETLTHTITK | 713.7 | 856.9 | 3 | y | 15 | 2 |
| KNG1 | Kininogen-1 | P01042 | DIPTNSPELEETLTHTITK | 713.7 | 756.4 | 3 | y | 13 | 2 |
| KNG1 | Kininogen-1 | P01042 | DIPTNSPELEETLTHTITK | 713.7 | 637.7 | 3 | y | 17 | 3 |
| KNG1 | Kininogen-1 | P01042 | YFIDFVAR | 515.8 | 867.5 | 2 | y | 7 | 1 |
| KNG1 | Kininogen-1 | P01042 | YFIDFVAR | 515.8 | 720.4 | 2 | y | 6 | 1 |
| KNG1 | Kininogen-1 | P01042 | YFIDFVAR | 515.8 | 607.3 | 2 | y | 5 | 1 |
| KNG1 | Kininogen-1 | P01042 | YFIDFVAR | 515.8 | 492.3 | 2 | y | 4 | 1 |
| KRT9 | Keratin, type I cytoskeletal 9 | P35527 | TLLDIDNTR | 530.8 | 846.4 | 2 | y | 7 | 1 |
| KRT9 | Keratin, type I cytoskeletal 9 | P35527 | TLLDIDNTR | 530.8 | 733.3 | 2 | y | 6 | 1 |
| KRT9 | Keratin, type I cytoskeletal 9 | P35527 | TLLDIDNTR | 530.8 | 618.3 | 2 | y | 5 | 1 |

| | | | | | | | | | |
|---|---|---|---|---|---|---|---|---|---|
| KRT9 | Keratin, type I cytoskeletal 9 | P35527 | TLLDIDNTR | 530.8 | 505.2 | 2 | y | 4 | 1 |
| KRTDAP | Keratinocyte differentiation-associated protein | P60985 | LPFLNWDAFPK | 674.4 | 990.5 | 2 | y | 8 | 1 |
| KRTDAP | Keratinocyte differentiation-associated protein | P60985 | LPFLNWDAFPK | 674.4 | 877.4 | 2 | y | 7 | 1 |
| KRTDAP | Keratinocyte differentiation-associated protein | P60985 | LPFLNWDAFPK | 674.4 | 617.8 | 2 | y | 10 | 2 |
| LCAT | Phosphatidylcholine-sterol acyltransferase | P04180 | SSGLVSNAPGVQIR | 692.9 | 1040.6 | 2 | y | 9 | 1 |
| LCAT | Phosphatidylcholine-sterol acyltransferase | P04180 | SSGLVSNAPGVQIR | 692.9 | 941.5 | 2 | y | 7 | 1 |
| LCAT | Phosphatidylcholine-sterol acyltransferase | P04180 | SSGLVSNAPGVQIR | 692.9 | 740.4 | 2 | y | 6 | 1 |
| LCAT | Phosphatidylcholine-sterol acyltransferase | P04180 | SSGLVSNAPGVQIR | 692.9 | 669.4 | 2 | y | 6 | 1 |
| LCP1 | Plastin-2 | P13796 | AYYHLLEQVAPK | 477.9 | 315.2 | 3 | y | 3 | 1 |
| LCP1 | Plastin-2 | P13796 | AYYHLLEQVAPK | 477.9 | 244.2 | 3 | y | 2 | 1 |
| LCP1 | Plastin-2 | P13796 | AYYHLLEQVAPK | 477.9 | 680.9 | 3 | y | 11 | 2 |
| LCP1 | Plastin-2 | P13796 | AYYHLLEQVAPK | 477.9 | 599.3 | 3 | y | 10 | 2 |
| LCP1 | Plastin-2 | P13796 | VYALPEDLVEVNPK | 793.4 | 1139.6 | 2 | y | 10 | 1 |
| LCP1 | Plastin-2 | P13796 | VYALPEDLVEVNPK | 793.4 | 358.2 | 2 | y | 3 | 1 |
| LCP1 | Plastin-2 | P13796 | VYALPEDLVEVNPK | 793.4 | 244.2 | 2 | y | 2 | 1 |
| LCP1 | Plastin-2 | P13796 | VYALPEDLVEVNPK | 793.4 | 570.3 | 2 | y | 10 | 2 |
| MAP1A | Microtubule-associated protein 1A | P78559 | VVSNTIEPLTLFHK | 533.3 | 855.5 | 3 | y | 7 | 1 |
| MAP1A | Microtubule-associated protein 1A | P78559 | VVSNTIEPLTLFHK | 533.3 | 645.4 | 3 | y | 5 | 1 |
| MAP1A | Microtubule-associated protein 1A | P78559 | VVSNTIEPLTLFHK | 533.3 | 700.4 | 3 | y | 12 | 2 |
| MAP1A | Microtubule-associated protein 1A | P78559 | VVSNTIEPLTLFHK | 533.3 | 428.3 | 3 | y | 7 | 2 |
| MAP2 | Microtubule-associated protein 2 | P11137 | EEFVETCPSEHK | 497.9 | 597.3 | 3 | y | 5 | 1 |
| MAP2 | Microtubule-associated protein 2 | P11137 | EEFVETCPSEHK | 497.9 | 617.3 | 3 | y | 10 | 2 |
| MAP2 | Microtubule-associated protein 2 | P11137 | EEFVETCPSEHK | 497.9 | 429.7 | 3 | y | 7 | 2 |
| MAP2 | Microtubule-associated protein 2 | P11137 | EEFVETCPSEHK | 497.9 | 379.2 | 3 | y | 6 | 2 |
| MB | Myoglobin | P02144 | GHHEAEIKPLAQSHATK | 464.2 | 543.3 | 4 | y | 5 | 1 |
| MB | Myoglobin | P02144 | GHHEAEIKPLAQSHATK | 464.2 | 476.8 | 4 | y | 9 | 2 |
| MB | Myoglobin | P02144 | GHHEAEIKPLAQSHATK | 464.2 | 554.0 | 4 | y | 15 | 3 |
| MB | Myoglobin | P02144 | GHHEAEIKPLAQSHATK | 464.2 | 398.6 | 4 | y | 11 | 3 |
| MYH7 | Myosin-7 | P12883 | DFELNALNAR | 581.8 | 771.4 | 2 | y | 7 | 1 |
| MYH7 | Myosin-7 | P12883 | DFELNALNAR | 581.8 | 658.4 | 2 | y | 6 | 1 |
| MYH7 | Myosin-7 | P12883 | DFELNALNAR | 581.8 | 544.3 | 2 | y | 5 | 1 |
| MYL1 | Myosin light chain 1/3, skeletal muscle isoform | P05976 | IPFQSHLPIQAAWR | 555.3 | 841.5 | 3 | y | 7 | 1 |
| MYL1 | Myosin light chain 1/3, skeletal muscle isoform | P05976 | IPFQSHLPIQAAWR | 555.3 | 775.9 | 3 | y | 13 | 2 |
| MYL1 | Myosin light chain 1/3, skeletal muscle isoform | P05976 | IPFQSHLPIQAAWR | 555.3 | 727.4 | 3 | y | 12 | 2 |
| MYL1 | Myosin light chain 1/3, skeletal muscle isoform | P05976 | IPFQSHLPIQAAWR | 555.3 | 421.2 | 3 | y | 7 | 2 |
| MYL3 | Myosin light chain 3 | P08590 | ITYGQCGDVLR | 641.3 | 1067.5 | 2 | y | 9 | 1 |
| MYL3 | Myosin light chain 3 | P08590 | ITYGQCGDVLR | 641.3 | 904.4 | 2 | y | 8 | 1 |
| MYL3 | Myosin light chain 3 | P08590 | ITYGQCGDVLR | 641.3 | 719.4 | 2 | y | 6 | 1 |
| MYL3 | Myosin light chain 3 | P08590 | ITYGQCGDVLR | 641.3 | 534.3 | 2 | y | 9 | 2 |
| MYOM2 | Myomesin-2 | P54296 | IESNYGVHTLEINR | 548.9 | 882.5 | 3 | y | 7 | 1 |
| MYOM2 | Myomesin-2 | P54296 | IESNYGVHTLEINR | 548.9 | 766.4 | 3 | y | 13 | 2 |
| MYOM2 | Myomesin-2 | P54296 | IESNYGVHTLEINR | 548.9 | 701.9 | 3 | y | 12 | 2 |
| MYOM2 | Myomesin-2 | P54296 | IESNYGVHTLEINR | 548.9 | 601.3 | 3 | y | 10 | 2 |
| MYOM2 | Myomesin-2 | P54296 | YPVTGLFEGR | 569.8 | 878.5 | 2 | y | 8 | 1 |
| MYOM2 | Myomesin-2 | P54296 | YPVTGLFEGR | 569.8 | 779.4 | 2 | y | 7 | 1 |
| MYOM2 | Myomesin-2 | P54296 | YPVTGLFEGR | 569.8 | 678.4 | 2 | y | 6 | 1 |
| MYOM2 | Myomesin-2 | P54296 | YPVTGLFEGR | 569.8 | 488.3 | 2 | y | 9 | 2 |
| NSG2 | Neuronal vesicle trafficking-associated protein 2 | Q9Y328 | FYTVISHYSVAK | 472.3 | 791.4 | 3 | y | 7 | 1 |
| NSG2 | Neuronal vesicle trafficking-associated protein 2 | Q9Y328 | FYTVISHYSVAK | 472.3 | 552.8 | 3 | y | 10 | 2 |
| NSG2 | Neuronal vesicle trafficking-associated protein 2 | Q9Y328 | FYTVISHYSVAK | 472.3 | 502.3 | 3 | y | 9 | 2 |
| NSG2 | Neuronal vesicle trafficking-associated protein 2 | Q9Y328 | FYTVISHYSVAK | 472.3 | 452.7 | 3 | y | 8 | 2 |
| NTM | Neurotrimin | Q9P121 | GTLQCEASAVPSAEFQWYK | 724.7 | 1155.5 | 3 | y | 9 | 1 |
| NTM | Neurotrimin | Q9P121 | GTLQCEASAVPSAEFQWYK | 724.7 | 1058.5 | 3 | y | 8 | 1 |
| NTM | Neurotrimin | Q9P121 | GTLQCEASAVPSAEFQWYK | 724.7 | 900.4 | 3 | y | 6 | 1 |
| NTM | Neurotrimin | Q9P121 | GTLQCEASAVPSAEFQWYK | 724.7 | 578.3 | 3 | y | 9 | 2 |
| OBSCN | Obscurin | Q5VST9 | ATLLNVLEGR | 543.3 | 800.5 | 2 | y | 7 | 1 |
| OBSCN | Obscurin | Q5VST9 | ATLLNVLEGR | 543.3 | 687.4 | 2 | y | 6 | 1 |
| OBSCN | Obscurin | Q5VST9 | ATLLNVLEGR | 543.3 | 573.3 | 2 | y | 5 | 1 |
| OLFM1 | Noelin | Q99784 | LTGISDPVTVK | 565.3 | 915.5 | 2 | y | 9 | 1 |
| OLFM1 | Noelin | Q99784 | LTGISDPVTVK | 565.3 | 745.4 | 2 | y | 7 | 1 |
| OLFM1 | Noelin | Q99784 | LTGISDPVTVK | 565.3 | 543.4 | 2 | y | 5 | 1 |
| OLFM1 | Noelin | Q99784 | LTGISDPVTVK | 565.3 | 347.2 | 2 | y | 3 | 1 |
| PCDH11X | Protocadherin-11 X-linked | Q9BZA7 | VIPLTTFTPR | 572.8 | 932.5 | 2 | y | 8 | 1 |
| PCDH11X | Protocadherin-11 X-linked | Q9BZA7 | VIPLTTFTPR | 572.8 | 722.4 | 2 | y | 6 | 1 |
| PCDH11X | Protocadherin-11 X-linked | Q9BZA7 | VIPLTTFTPR | 572.8 | 621.3 | 2 | y | 5 | 1 |

| | | | | | | | | | |
|---|---|---|---|---|---|---|---|---|---|
| PCDH11X | Protocadherin-11 X-linked | Q9BZA7 | VIPLTTFTPR | 572.8 | 466.8 | 2 | y | 8 | 2 |
| PF4 | Platelet factor 4 | P02776 | AGPHCPTAQLIATLK | 526.6 | 786.5 | 3 | y | 7 | 1 |
| PF4 | Platelet factor 4 | P02776 | AGPHCPTAQLIATLK | 526.6 | 658.4 | 3 | y | 6 | 1 |
| PF4 | Platelet factor 4 | P02776 | AGPHCPTAQLIATLK | 526.6 | 545.4 | 3 | y | 5 | 1 |
| PF4 | Platelet factor 4 | P02776 | AGPHCPTAQLIATLK | 526.6 | 432.3 | 3 | y | 4 | 1 |
| PF4 | Platelet factor 4 | P02776 | ICLDLQAPLYK | 667.4 | 1060.6 | 2 | y | 9 | 1 |
| PF4 | Platelet factor 4 | P02776 | ICLDLQAPLYK | 667.4 | 947.5 | 2 | y | 8 | 1 |
| PF4 | Platelet factor 4 | P02776 | ICLDLQAPLYK | 667.4 | 591.4 | 2 | y | 5 | 1 |
| PF4 | Platelet factor 4 | P02776 | ICLDLQAPLYK | 667.4 | 520.3 | 2 | y | 4 | 1 |
| PGLYRP2 | N-acetylmuramoyl-L-alanine amidase | Q96PD5 | EFTEAFLGCPAIHPR | 582.3 | 690.4 | 3 | y | 6 | 1 |
| PGLYRP2 | N-acetylmuramoyl-L-alanine amidase | Q96PD5 | EFTEAFLGCPAIHPR | 582.3 | 619.8 | 3 | y | 11 | 2 |
| PGLYRP2 | N-acetylmuramoyl-L-alanine amidase | Q96PD5 | EFTEAFLGCPAIHPR | 582.3 | 584.3 | 3 | y | 10 | 2 |
| PGLYRP2 | N-acetylmuramoyl-L-alanine amidase | Q96PD5 | EFTEAFLGCPAIHPR | 582.3 | 345.7 | 3 | y | 6 | 2 |
| PGLYRP2 | N-acetylmuramoyl-L-alanine amidase | Q96PD5 | GCPDVQASLPDAK | 679.3 | 1140.6 | 2 | y | 11 | 1 |
| PGLYRP2 | N-acetylmuramoyl-L-alanine amidase | Q96PD5 | GCPDVQASLPDAK | 679.3 | 829.4 | 2 | y | 8 | 1 |
| PGLYRP2 | N-acetylmuramoyl-L-alanine amidase | Q96PD5 | GCPDVQASLPDAK | 679.3 | 430.2 | 2 | y | 4 | 1 |
| PGLYRP2 | N-acetylmuramoyl-L-alanine amidase | Q96PD5 | GCPDVQASLPDAK | 679.3 | 570.8 | 2 | y | 11 | 2 |
| PKLR | Pyruvate kinase PKLR | P30613 | GDLGIEIPAEK | 571.3 | 856.5 | 2 | y | 8 | 1 |
| PKLR | Pyruvate kinase PKLR | P30613 | GDLGIEIPAEK | 571.3 | 686.4 | 2 | y | 6 | 1 |
| PKLR | Pyruvate kinase PKLR | P30613 | GDLGIEIPAEK | 571.3 | 557.3 | 2 | y | 5 | 1 |
| PKLR | Pyruvate kinase PKLR | P30613 | GDLGIEIPAEK | 571.3 | 444.2 | 2 | y | 4 | 1 |
| PLG | Plasminogen | P00747 | CTTPPPSSGPTYQCLK | 598.6 | 909.4 | 3 | y | 7 | 1 |
| PLG | Plasminogen | P00747 | CTTPPPSSGPTYQCLK | 598.6 | 711.3 | 3 | y | 5 | 1 |
| PLG | Plasminogen | P00747 | CTTPPPSSGPTYQCLK | 598.6 | 483.7 | 3 | y | 8 | 2 |
| PLG | Plasminogen | P00747 | CTTPPPSSGPTYQCLK | 598.6 | 455.2 | 3 | y | 7 | 2 |
| PLG | Plasminogen | P00747 | EAQLPVIENK | 570.8 | 812.5 | 2 | y | 7 | 1 |
| PLG | Plasminogen | P00747 | EAQLPVIENK | 570.8 | 699.4 | 2 | y | 6 | 1 |
| PLG | Plasminogen | P00747 | EAQLPVIENK | 570.8 | 503.3 | 2 | y | 4 | 1 |
| PLG | Plasminogen | P00747 | EAQLPVIENK | 570.8 | 350.2 | 2 | y | 6 | 2 |
| PNCK | Calcium/calmodulin-dependent protein kinase type 1B | Q6P2M8 | HLWISGDTAFDR | 473.2 | 609.3 | 3 | y | 5 | 1 |
| PNCK | Calcium/calmodulin-dependent protein kinase type 1B | Q6P2M8 | HLWISGDTAFDR | 473.2 | 508.3 | 3 | y | 4 | 1 |
| PNCK | Calcium/calmodulin-dependent protein kinase type 1B | Q6P2M8 | HLWISGDTAFDR | 473.2 | 434.7 | 3 | y | 8 | 2 |
| PON1 | Serum paraoxonase/arylesterase 1 | P27169 | IFFYDSENPPASEVLR | 628.6 | 868.5 | 3 | y | 8 | 1 |
| PON1 | Serum paraoxonase/arylesterase 1 | P27169 | IFFYDSENPPASEVLR | 628.6 | 771.4 | 3 | y | 7 | 1 |
| PON1 | Serum paraoxonase/arylesterase 1 | P27169 | IFFYDSENPPASEVLR | 628.6 | 434.7 | 3 | y | 8 | 2 |
| PON1 | Serum paraoxonase/arylesterase 1 | P27169 | IFFYDSENPPASEVLR | 628.6 | 386.2 | 3 | y | 7 | 2 |
| PON1 | Serum paraoxonase/arylesterase 1 | P27169 | IQNILTEEPK | 592.8 | 943.5 | 2 | y | 8 | 1 |
| PON1 | Serum paraoxonase/arylesterase 1 | P27169 | IQNILTEEPK | 592.8 | 829.5 | 2 | y | 7 | 1 |
| PON1 | Serum paraoxonase/arylesterase 1 | P27169 | IQNILTEEPK | 592.8 | 716.4 | 2 | y | 6 | 1 |
| PON1 | Serum paraoxonase/arylesterase 1 | P27169 | IQNILTEEPK | 592.8 | 603.3 | 2 | y | 5 | 1 |
| PPBP | Platelet basic protein | P02775 | GTHCNQVEVIATLK | 523.9 | 773.5 | 3 | y | 7 | 1 |
| PPBP | Platelet basic protein | P02775 | GTHCNQVEVIATLK | 523.9 | 644.4 | 3 | y | 6 | 1 |
| PPBP | Platelet basic protein | P02775 | GTHCNQVEVIATLK | 523.9 | 545.4 | 3 | y | 5 | 1 |
| PPBP | Platelet basic protein | P02775 | GTHCNQVEVIATLK | 523.9 | 432.3 | 3 | y | 4 | 1 |
| PPBP | Platelet basic protein | P02775 | ICLDPDAPR | 528.8 | 783.4 | 2 | y | 7 | 1 |
| PPBP | Platelet basic protein | P02775 | ICLDPDAPR | 528.8 | 670.3 | 2 | y | 6 | 1 |
| PPBP | Platelet basic protein | P02775 | ICLDPDAPR | 528.8 | 555.3 | 2 | y | 5 | 1 |
| PPBP | Platelet basic protein | P02775 | ICLDPDAPR | 528.8 | 343.2 | 2 | y | 3 | 1 |
| PRC1 | Protein regulator of cytokinesis 1 | O43663 | AWTDVLPWK | 558.3 | 1044.6 | 2 | y | 8 | 1 |
| PRC1 | Protein regulator of cytokinesis 1 | O43663 | AWTDVLPWK | 558.3 | 858.5 | 2 | y | 7 | 1 |
| PRC1 | Protein regulator of cytokinesis 1 | O43663 | AWTDVLPWK | 558.3 | 757.4 | 2 | y | 6 | 1 |
| PRC1 | Protein regulator of cytokinesis 1 | O43663 | AWTDVLPWK | 558.3 | 642.4 | 2 | y | 5 | 1 |
| PRG4 | Proteoglycan 4 | Q92954 | DQYYNIDVPSR | 685.3 | 800.4 | 2 | y | 7 | 1 |
| PRG4 | Proteoglycan 4 | Q92954 | DQYYNIDVPSR | 685.3 | 573.3 | 2 | y | 5 | 1 |
| PRG4 | Proteoglycan 4 | Q92954 | DQYYNIDVPSR | 685.3 | 458.3 | 2 | y | 4 | 1 |
| PRG4 | Proteoglycan 4 | Q92954 | DQYYNIDVPSR | 685.3 | 359.2 | 2 | y | 3 | 1 |
| PRG4 | Proteoglycan 4 | Q92954 | GLPNVVTSAISLPNIR | 826.0 | 1071.6 | 2 | y | 10 | 1 |
| PRG4 | Proteoglycan 4 | Q92954 | GLPNVVTSAISLPNIR | 826.0 | 970.6 | 2 | y | 9 | 1 |
| PRG4 | Proteoglycan 4 | Q92954 | GLPNVVTSAISLPNIR | 826.0 | 499.3 | 2 | y | 4 | 1 |
| PRG4 | Proteoglycan 4 | Q92954 | GLPNVVTSAISLPNIR | 826.0 | 740.9 | 2 | y | 14 | 2 |
| PTGDS | Prostaglandin-H2 D-isomerase | P41222 | AQGFTEDTIVFLPQTDK | 955.5 | 947.5 | 2 | y | 8 | 1 |
| PTGDS | Prostaglandin-H2 D-isomerase | P41222 | AQGFTEDTIVFLPQTDK | 955.5 | 848.5 | 2 | y | 7 | 1 |
| PTGDS | Prostaglandin-H2 D-isomerase | P41222 | AQGFTEDTIVFLPQTDK | 955.5 | 701.4 | 2 | y | 6 | 1 |
| PTGDS | Prostaglandin-H2 D-isomerase | P41222 | AQGFTEDTIVFLPQTDK | 955.5 | 588.3 | 2 | y | 5 | 1 |
| RAB11B | Ras-related protein Rab-11B | Q15907 | AITSAYYR | 472.7 | 873.4 | 2 | y | 7 | 1 |

| | | | | | | | | | |
|---|---|---|---|---|---|---|---|---|---|
| RAB11B | Ras-related protein Rab-11B | Q15907 | AITSAYYR | 472.7 | 760.4 | 2 | y | 6 | 1 |
| RAB11B | Ras-related protein Rab-11B | Q15907 | AITSAYYR | 472.7 | 659.3 | 2 | y | 5 | 1 |
| RAB11B | Ras-related protein Rab-11B | Q15907 | AITSAYYR | 472.7 | 572.3 | 2 | y | 4 | 1 |
| RAB11B | Ras-related protein Rab-11B | B4DMK0 | AQIWDTAGQER | 637.8 | 962.4 | 2 | y | 8 | 1 |
| RAB11B | Ras-related protein Rab-11B | B4DMK0 | AQIWDTAGQER | 637.8 | 776.4 | 2 | y | 7 | 1 |
| RAB11B | Ras-related protein Rab-11B | B4DMK0 | AQIWDTAGQER | 637.8 | 560.3 | 2 | y | 5 | 1 |
| RAB11B | Ras-related protein Rab-11B | B4DMK0 | AQIWDTAGQER | 637.8 | 489.2 | 2 | y | 4 | 1 |
| RGS20 | Regulator of G-protein signaling 20 | O76081 | LFGLLSSPLSSLAR | 730.9 | 1030.6 | 2 | y | 10 | 1 |
| RGS20 | Regulator of G-protein signaling 20 | O76081 | LFGLLSSPLSSLAR | 730.9 | 917.5 | 2 | y | 9 | 1 |
| RGS20 | Regulator of G-protein signaling 20 | O76081 | LFGLLSSPLSSLAR | 730.9 | 830.5 | 2 | y | 8 | 1 |
| RGS20 | Regulator of G-protein signaling 20 | O76081 | LFGLLSSPLSSLAR | 730.9 | 743.4 | 2 | y | 7 | 1 |
| RUNDC3A | RUN domain-containing protein 3A | Q59EK9 | TPVVIDYTPYLK | 704.9 | 1012.5 | 2 | y | 8 | 1 |
| RUNDC3A | RUN domain-containing protein 3A | Q59EK9 | TPVVIDYTPYLK | 704.9 | 899.5 | 2 | y | 7 | 1 |
| RUNDC3A | RUN domain-containing protein 3A | Q59EK9 | TPVVIDYTPYLK | 704.9 | 784.4 | 2 | y | 6 | 1 |
| RUNDC3A | RUN domain-containing protein 3A | Q59EK9 | TPVVIDYTPYLK | 704.9 | 520.3 | 2 | y | 4 | 1 |
| S100A9 | Protein S100-A9 | P06702 | LGHPDTLNQGEFK | 485.9 | 722.3 | 3 | y | 6 | 1 |
| S100A9 | Protein S100-A9 | P06702 | LGHPDTLNQGEFK | 485.9 | 608.3 | 3 | y | 5 | 1 |
| S100A9 | Protein S100-A9 | P06702 | LGHPDTLNQGEFK | 485.9 | 294.2 | 3 | y | 2 | 1 |
| SAA1 | Serum amyloid A-1 protein | P0DJI8 | GPGGVWAAEAISDAR | 728.9 | 1089.5 | 2 | y | 10 | 1 |
| SAA1 | Serum amyloid A-1 protein | P0DJI8 | GPGGVWAAEAISDAR | 728.9 | 903.5 | 2 | y | 9 | 1 |
| SAA1 | Serum amyloid A-1 protein | P0DJI8 | GPGGVWAAEAISDAR | 728.9 | 832.4 | 2 | y | 8 | 1 |
| SAA1 | Serum amyloid A-1 protein | P0DJI8 | GPGGVWAAEAISDAR | 728.9 | 761.4 | 2 | y | 7 | 1 |
| SAA2 | Serum amyloid A-2 protein | P0DJI9 | EANYIGSDK | 498.7 | 682.3 | 2 | y | 6 | 1 |
| SAA2 | Serum amyloid A-2 protein | P0DJI9 | EANYIGSDK | 498.7 | 519.3 | 2 | y | 5 | 1 |
| SAA2 | Serum amyloid A-2 protein | P0DJI9 | EANYIGSDK | 498.7 | 406.2 | 2 | y | 4 | 1 |
| SAA2 | Serum amyloid A-2 protein | P0DJI9 | EANYIGSDK | 498.7 | 349.2 | 2 | y | 3 | 1 |
| SCGB1A1 | Uteroglobin | P11684 | LVDTLPQKPR | 389.6 | 482.8 | 3 | y | 9 | 2 |
| SCGB1A1 | Uteroglobin | P11684 | LVDTLPQKPR | 389.6 | 625.4 | 3 | y | 5 | 1 |
| SCGB1A1 | Uteroglobin | P11684 | LVDTLPQKPR | 389.6 | 527.3 | 3 | y | 9 | 2 |
| SCGB1A1 | Uteroglobin | P11684 | LVDTLPQKPR | 389.6 | 313.2 | 3 | y | 5 | 2 |
| SELL | L-selectin | P14151 | SLTEEAENWGDGEPNNK | 630.6 | 830.4 | 3 | y | 8 | 1 |
| SELL | L-selectin | P14151 | SLTEEAENWGDGEPNNK | 630.6 | 773.3 | 3 | y | 7 | 1 |
| SELL | L-selectin | P14151 | SLTEEAENWGDGEPNNK | 630.6 | 472.3 | 3 | y | 4 | 1 |
| SERPINA10 | Protein Z-dependent protease inhibitor | Q9UK55 | IFSPFADLSELSATGR | 855.9 | 1119.6 | 2 | y | 11 | 1 |
| SERPINA10 | Protein Z-dependent protease inhibitor | Q9UK55 | IFSPFADLSELSATGR | 855.9 | 1048.5 | 2 | y | 10 | 1 |
| SERPINA10 | Protein Z-dependent protease inhibitor | Q9UK55 | IFSPFADLSELSATGR | 855.9 | 820.4 | 2 | y | 8 | 1 |
| SERPINA10 | Protein Z-dependent protease inhibitor | Q9UK55 | IFSPFADLSELSATGR | 855.9 | 682.3 | 2 | y | 13 | 2 |
| SERPINA12 | Serpin A12 | Q8IW75 | IFEEHGDLTK | 396.9 | 538.3 | 3 | y | 9 | 2 |
| SERPINA12 | Serpin A12 | Q8IW75 | IFEEHGDLTK | 396.9 | 464.7 | 3 | y | 8 | 2 |
| SERPINA12 | Serpin A12 | Q8IW75 | IFEEHGDLTK | 396.9 | 400.2 | 3 | y | 7 | 2 |
| SERPINA7 | Thyroxine-binding globulin | P05543 | GWVDLFVPK | 530.8 | 343.2 | 2 | y | 3 | 1 |
| SERPINA7 | Thyroxine-binding globulin | P05543 | GWVDLFVPK | 530.8 | 817.5 | 2 | y | 7 | 1 |
| SERPINA7 | Thyroxine-binding globulin | P05543 | GWVDLFVPK | 530.8 | 718.4 | 2 | y | 6 | 1 |
| SERPINA7 | Thyroxine-binding globulin | P05543 | GWVDLFVPK | 530.8 | 603.4 | 2 | y | 5 | 1 |
| SERPINC1 | Antithrombin-III | P01008 | TSDQIHFFFAK | 447.6 | 796.4 | 3 | y | 6 | 1 |
| SERPINC1 | Antithrombin-III | P01008 | TSDQIHFFFAK | 447.6 | 659.4 | 3 | y | 5 | 1 |
| SERPINC1 | Antithrombin-III | P01008 | TSDQIHFFFAK | 447.6 | 620.3 | 3 | y | 10 | 2 |
| SERPINC1 | Antithrombin-III | P01008 | TSDQIHFFFAK | 447.6 | 576.8 | 3 | y | 9 | 2 |
| SERPINF2 | Alpha-2-antiplasmin | P08697 | WFLLEQPEIQVAHFPFK | 710.4 | 1312.7 | 3 | y | 11 | 1 |
| SERPINF2 | Alpha-2-antiplasmin | P08697 | WFLLEQPEIQVAHFPFK | 710.4 | 898.5 | 3 | y | 15 | 2 |
| SERPINF2 | Alpha-2-antiplasmin | P08697 | WFLLEQPEIQVAHFPFK | 710.4 | 785.4 | 3 | y | 13 | 2 |
| SERPINF2 | Alpha-2-antiplasmin | P08697 | WFLLEQPEIQVAHFPFK | 710.4 | 656.9 | 3 | y | 11 | 2 |
| SERPING1 | Plasma protease C1 inhibitor | P05155 | GVTSVSQIFHSPDLAIR | 609.7 | 835.9 | 3 | y | 15 | 2 |
| SERPING1 | Plasma protease C1 inhibitor | P05155 | GVTSVSQIFHSPDLAIR | 609.7 | 785.4 | 3 | y | 14 | 2 |
| SERPING1 | Plasma protease C1 inhibitor | P05155 | GVTSVSQIFHSPDLAIR | 609.7 | 741.9 | 3 | y | 13 | 2 |
| SERPING1 | Plasma protease C1 inhibitor | P05155 | GVTSVSQIFHSPDLAIR | 609.7 | 692.4 | 3 | y | 12 | 2 |
| SERPING1 | Plasma protease C1 inhibitor | P05155 | TNLESILSYPK | 632.8 | 936.5 | 2 | y | 8 | 1 |
| SERPING1 | Plasma protease C1 inhibitor | P05155 | TNLESILSYPK | 632.8 | 807.5 | 2 | y | 7 | 1 |
| SERPING1 | Plasma protease C1 inhibitor | P05155 | TNLESILSYPK | 632.8 | 494.3 | 2 | y | 4 | 1 |
| SERPING1 | Plasma protease C1 inhibitor | P05155 | TNLESILSYPK | 632.8 | 407.2 | 2 | y | 3 | 1 |
| SHC3 | SHC-transforming protein 3 | Q92529 | GAPHASDQVLGPGVTYVVK | 632.3 | 1032.6 | 3 | y | 10 | 1 |
| SHC3 | SHC-transforming protein 3 | Q92529 | GAPHASDQVLGPGVTYVVK | 632.3 | 919.5 | 3 | y | 9 | 1 |
| SHC3 | SHC-transforming protein 3 | Q92529 | GAPHASDQVLGPGVTYVVK | 632.3 | 609.4 | 3 | y | 5 | 1 |
| SHC3 | SHC-transforming protein 3 | Q92529 | GAPHASDQVLGPGVTYVVK | 632.3 | 460.3 | 3 | y | 9 | 2 |
| SLC36A2 | Proton-coupled amino acid transporter 2 | Q495M3 | WALPLDLSIR | 592.3 | 926.6 | 2 | y | 8 | 1 |

| | | | | | | | | | |
|---|---|---|---|---|---|---|---|---|---|
| SLC36A2 | Proton-coupled amino acid transporter 2 | Q495M3 | WALPLDLSIR | 592.3 | 813.5 | 2 | y | 7 | 1 |
| SLC36A2 | Proton-coupled amino acid transporter 2 | Q495M3 | WALPLDLSIR | 592.3 | 407.2 | 2 | y | 7 | 2 |
| SLC5A1 | Sodium/glucose cotransporter 1 | P13866 | TTAVTRPVETHELIR | 431.5 | 541.3 | 4 | y | 14 | 3 |
| SLC5A1 | Sodium/glucose cotransporter 1 | P13866 | TTAVTRPVETHELIR | 431.5 | 507.6 | 4 | y | 13 | 3 |
| SLC5A1 | Sodium/glucose cotransporter 1 | P13866 | TTAVTRPVETHELIR | 431.5 | 483.9 | 4 | y | 12 | 3 |
| SLC5A1 | Sodium/glucose cotransporter 1 | P13866 | TTAVTRPVETHELIR | 431.5 | 450.9 | 4 | y | 11 | 3 |
| SRL | Sarcalumenin | Q86TD4 | DFFGINPISSFK | 686.4 | 1109.6 | 2 | y | 10 | 1 |
| SRL | Sarcalumenin | Q86TD4 | DFFGINPISSFK | 686.4 | 962.5 | 2 | y | 9 | 1 |
| SRL | Sarcalumenin | Q86TD4 | DFFGINPISSFK | 686.4 | 792.4 | 2 | y | 7 | 1 |
| SRL | Sarcalumenin | Q86TD4 | DFFGINPISSFK | 686.4 | 678.4 | 2 | y | 6 | 1 |
| STX1B | Syntaxin-1B | P61266 | DSDDEEEVVHVDR | 515.2 | 724.4 | 3 | y | 6 | 1 |
| STX1B | Syntaxin-1B | P61266 | DSDDEEEVVHVDR | 515.2 | 625.3 | 3 | y | 5 | 1 |
| STX1B | Syntaxin-1B | P61266 | DSDDEEEVVHVDR | 515.2 | 671.3 | 3 | y | 11 | 2 |
| TBX5 | T-box transcription factor TBX5 | Q99593 | LPYQHFSAHFTSGPLVPR | 514.3 | 725.4 | 4 | y | 7 | 1 |
| TBX5 | T-box transcription factor TBX5 | Q99593 | LPYQHFSAHFTSGPLVPR | 514.3 | 291.2 | 4 | y | 5 | 2 |
| TBX5 | T-box transcription factor TBX5 | Q99593 | LPYQHFSAHFTSGPLVPR | 514.3 | 647.7 | 4 | y | 17 | 3 |
| TEAD1 | Transcriptional enhancer factor TEF-1 | P28347 | GPQNAFFLVK | 560.8 | 653.4 | 2 | y | 5 | 1 |
| TEAD1 | Transcriptional enhancer factor TEF-1 | P28347 | GPQNAFFLVK | 560.8 | 506.3 | 2 | y | 4 | 1 |
| TEAD1 | Transcriptional enhancer factor TEF-1 | P28347 | GPQNAFFLVK | 560.8 | 359.3 | 2 | y | 3 | 1 |
| TEAD1 | Transcriptional enhancer factor TEF-1 | P28347 | GPQNAFFLVK | 560.8 | 246.2 | 2 | y | 2 | 1 |
| TEAD1 | Transcriptional enhancer factor TEF-1 | P28347 | IILSDEGK | 437.7 | 761.4 | 2 | y | 7 | 1 |
| TEAD1 | Transcriptional enhancer factor TEF-1 | P28347 | IILSDEGK | 437.7 | 648.3 | 2 | y | 6 | 1 |
| TEAD1 | Transcriptional enhancer factor TEF-1 | P28347 | IILSDEGK | 437.7 | 535.2 | 2 | y | 5 | 1 |
| TEAD1 | Transcriptional enhancer factor TEF-1 | P28347 | IILSDEGK | 437.7 | 204.1 | 2 | y | 2 | 1 |
| TYRP1 | 5,6-dihydroxyindole-2-carboxylic acid oxidase | P17643 | YNADISTFPLENAPIGHNR | 710.4 | 926.5 | 3 | y | 17 | 2 |
| TYRP1 | 5,6-dihydroxyindole-2-carboxylic acid oxidase | P17643 | YNADISTFPLENAPIGHNR | 710.4 | 776.9 | 3 | y | 14 | 2 |
| TYRP1 | 5,6-dihydroxyindole-2-carboxylic acid oxidase | P17643 | YNADISTFPLENAPIGHNR | 710.4 | 682.9 | 3 | y | 12 | 2 |
| TYRP1 | 5,6-dihydroxyindole-2-carboxylic acid oxidase | P17643 | YNADISTFPLENAPIGHNR | 710.4 | 609.3 | 3 | y | 11 | 2 |
| VASP | Vasodilator-stimulated phosphoprotein | P50552 | VQIYHNPTANSFR | 516.3 | 792.4 | 3 | y | 7 | 1 |
| VASP | Vasodilator-stimulated phosphoprotein | P50552 | VQIYHNPTANSFR | 516.3 | 724.4 | 3 | y | 12 | 2 |
| VASP | Vasodilator-stimulated phosphoprotein | P50552 | VQIYHNPTANSFR | 516.3 | 660.3 | 3 | y | 11 | 2 |
| VASP | Vasodilator-stimulated phosphoprotein | P50552 | VQIYHNPTANSFR | 516.3 | 522.3 | 3 | y | 9 | 2 |
| VASP | Vasodilator-stimulated phosphoprotein | P50552 | YNQATPNFHQWR | 521.2 | 642.8 | 3 | y | 10 | 2 |
| VASP | Vasodilator-stimulated phosphoprotein | P50552 | YNQATPNFHQWR | 521.2 | 578.8 | 3 | y | 9 | 2 |
| VASP | Vasodilator-stimulated phosphoprotein | P50552 | YNQATPNFHQWR | 521.2 | 543.3 | 3 | y | 8 | 2 |
| VASP | Vasodilator-stimulated phosphoprotein | P50552 | YNQATPNFHQWR | 521.2 | 492.7 | 3 | y | 7 | 2 |
| VTN | Vitronectin | P04004 | VDTVDPPYPR | 579.8 | 744.4 | 2 | y | 6 | 1 |
| VTN | Vitronectin | P04004 | VDTVDPPYPR | 579.8 | 629.3 | 2 | y | 5 | 1 |
| VTN | Vitronectin | P04004 | VDTVDPPYPR | 579.8 | 532.3 | 2 | y | 4 | 1 |
| VTN | Vitronectin | P04004 | VDTVDPPYPR | 579.8 | 315.2 | 2 | y | 5 | 2 |

**Table S4.** Serum level changes of the 10 individual proteins (***t*-Test Set**) in seronegative and seropositive subgroups in the SLICE cohort.

T-Test *P* values in RED: $P < 0.05$; C: control (n=20); Sero-: seronegative (n=17); Sero+: seropositive (n=23)

| Protein | Peptide | Average FC to healthy controls | | | Original *t* test *P* value | | | Correlation coefficient to duration of illness |
|---|---|---|---|---|---|---|---|---|
| | | Sero - | Sero + | Control | Sero - vs. C | Sero + vs. C | Sero + vs. Sero - | |
| AFM | LPNNVLQEK | 0.73 ± 0.29 | 0.87 ± 0.4 | 1 ± 0.36 | 2.31E-03 | 2.15E-01 | 2.33E-01 | -0.02 |
| ALDOB | ALQASALAA\ | 1.65 ± 1.39 | 2.12 ± 2.18 | 1 ± 0.54 | 4.70E-02 | 5.01E-02 | 4.75E-01 | 0.06 |
| APOA4 | LGPHAGDVE( | 0.66 ± 0.42 | 0.6 ± 0.29 | 1 ± 0.47 | 5.25E-03 | 4.65E-04 | 7.55E-01 | 0.03 |
| APOA4 | SELTQQLNAL | 0.67 ± 0.48 | 0.6 ± 0.36 | 1 ± 0.42 | 9.64E-03 | 7.69E-04 | 6.71E-01 | 0.17 |
| C9 | LSPIYNLVPVK | 1.73 ± 1.24 | 2.21 ± 1.49 | 1 ± 0.25 | 1.20E-02 | 3.67E-03 | 3.32E-01 | -0.17 |
| CRP | ESDTSYVSLK | 21.44 ± 23.8: | 42.56 ± 68.2: | 1 ± 1.33 | 6.10E-04 | 2.25E-02 | 2.62E-01 | -0.09 |
| CRP | GYSIFSYATK | 17.53 ± 19.8 | 31.56 ± 51.6: | 1 ± 1.22 | 7.99E-04 | 2.58E-02 | 3.32E-01 | -0.09 |
| CST6 | AQSQLVAGIK | 0.61 ± 0.24 | 0.7 ± 0.38 | 1 ± 0.26 | 4.19E-07 | 3.80E-03 | 4.17E-01 | -0.16 |
| FBP1 | EAVLDVIPTDI | 1.77 ± 1.53 | 1.83 ± 1.62 | 1 ± 0.51 | 3.12E-02 | 5.28E-02 | 9.72E-01 | -0.17 |
| ITIH2 | VQFELHYQE\ | 0.75 ± 0.36 | 0.76 ± 0.32 | 1 ± 0.27 | 8.27E-03 | 6.97E-03 | 9.89E-01 | -0.07 |
| PGLYRP2 | EFTEAFLGCP. | 0.69 ± 0.31 | 0.81 ± 0.38 | 1 ± 0.33 | 6.64E-04 | 6.36E-02 | 3.18E-01 | -0.12 |
| PGLYRP2 | GCPDVQASLI | 0.82 ± 0.34 | 0.77 ± 0.28 | 1 ± 0.23 | 3.40E-02 | 3.51E-03 | 5.69E-01 | -0.18 |
| S100A9 | LGHPDTLNQ( | 1.46 ± 1.01 | 1.85 ± 1.01 | 1 ± 0.47 | 4.33E-02 | 5.63E-03 | 3.45E-01 | -0.27 |

**Table S5**. Performance of the **MVA Panel** in stratifying seronegative and seropositive LD subgroups from healthy controls in both SLICE and NYMC sets

Lyme: All Lyme disease patients in the cohort; Sero -: seronegative; Sero +: seropositive; Cntl: control

| Dataset | Accuracy | Error Rate | Sensitivity | Specificity | AUC |
|---|---|---|---|---|---|
| **JHU_SLICE (10-protein MVA panel)** | | | | | |
| Lyme *vs* Cntl | 0.912 | 0.088 | 0.95 | 0.925 | 0.983 |
| Sero - *vs* Cntl | 0.929 | 0.071 | 0.864 | 1 | 0.911 |
| Sero + *vs* Cntl | 0.816 | 0.184 | 0.833 | 0.8 | 0.851 |
| **NYMC (9-protein MVA panel)** | | | | | |
| Lyme *vs* Cntl | 0.838 | 0.162 | 0.75 | 0.9 | 0.832 |
| Sero - *vs* Cntl | 0.868 | 0.132 | 0.947 | 0.789 | 0.904 |
| Sero + *vs* Cntl | 0.816 | 0.184 | 0.684 | 0.947 | 0.842 |
